# Sleep increases firing rate modulation during interictal epileptic activities in mesial temporal structures

**DOI:** 10.1101/2022.12.30.522096

**Authors:** Stephen Whitmarsh, Vi-Huong Nguyen-Michel, Katia Lehongre, Bertrand Mathon, Claude Adam, Virginie Lambrecq, Valerio Frazzini, Vincent Navarro

## Abstract

Epileptic seizures and interictal epileptiform discharges (IEDs) are strongly influenced by sleep and circadian rhythms. However, human data on the effect of sleep on neuronal behavior during interictal activity have been lacking. We analyzed EEG data from epileptic patients implanted with macro and micro electrodes targeting mesial temporal structures. Sleep staging was performed on concomitantly recorded polysomnography and video-EEG. Automated IED detection identified thousands of IEDs per patient. Both the rate and amplitude of IEDs were increased with deeper stages of NREM sleep. Single unit activity (SUA) and multi-unit activity (MUA) increased their firing during the IED spike, and strongly decreased during the subsequent slow wave. These time-locked firing rate modulations were shown to increase during deeper stages of NREM sleep. Finally, during resting behaviour, neuronal firing rate, bursting rate and firing regularity were all shown to progressively decrease with deeper stages of NREM sleep.

## Introduction

Early modern research recognized that epileptic activity occurs in circadian and ultradian cycles (1; 2; 3). Since the advent of electroencephalography (EEG) and polysomnography (PSG), it is now known that many types of epilepsy (focal and idiopathic generalized) have the highest seizure preponderance during light non-REM sleep (NREM), and the lowest during REM sleep (4; 5; 6; 7; 8; 9; 10; 11; 12; 13). In temporal epilepsy, focal seizures were also found to be more likely to evolve into bilateral tonic-clonic seizures during sleep than during wakefulness (9; 8; 6). In addition, sleep deprivation is the most common trigger for spontaneous “breakthrough seizures” (14; 15; 16; 17; 18) and seizures in the epilepsy clinic (19; 20), regardless of the type of epilepsy or the epilepsy syndrome. Furthermore, sleep disorders such as sleep apneas, that strongly disturb the quality of sleep, likely worsen epilepsy (21; 22).

Interictal epileptiform discharges (IEDs) are brief non-symptomatic events observed in the EEG of patients predisposed to spontaneous seizures (23). IEDs are known to be strongly influenced by sleep and circadian rhythms as well. (24) were the first to demonstrate that the rate of IED occurrence increases with sleep, even if the waking EEG was normal. Using invasive EEG, (25) later reported an increase of IEDs when certain patients became drowsy or fell asleep. Since then, IEDs have been consistently found to occur more often during sleep (26; 27; 28; 29; 30; 31; 32; 33; 34; 35). Indeed, long-term recordings have found robust 24 hour periods, peaking between midnight and 5pm (36; 33; 37). Generally, deeper stages of non-REM (NREM) sleep seem to promote interictal activity, while lighter stages may promote seizures (9; 11), especially in temporal lobe epilepsy (26; 31; 32). The average IED rate is lower in rapid eye movement (REM) sleep than in wakefulness, and higher in slow wave sleep (SWS) than in REM (38). Furthermore, IED rates are positively correlated with power in the delta (1 Hz to 4 Hz) frequency range (27; 39; 40).

IEDs are understood to be generated by synchronous neuronal discharges, occurring during depolarization of the neuronal membrane (41; 42; 43), consistent with paroxysmal depolarizing shifts (PDS) studied in animal models of epilepsy, in which a large depolarization phase is followed by a long hyperpolarization (44; 45). In humans, microelectrode recordings have shown that firing rates increase during the interictal spike, and are subsequently reduced (and often completely silenced) during the subsequent slow wave (46; 47; 43; 48). The effect of sleep on seizures and IEDs are hypothesized to be due to enhanced brain synchronization during SWS, and decreased synchronization during REM (49; 50). Consistent with sleep-related changes in cortical excitability, human cortical and hippocampal neurons have been shown to have a higher propensity for bursting during NREM, with the lowest bursting rates during REM (51; 52; 53). Although these findings provide a pathophysiological model on the level of single unit and population activity, no studies have yet demonstrated an effect of sleep on neuronal firing behavior during IEDs in human data.

Several studies have shown the clinical potential of basic morphological metrics of IEDs. Aanestad et al. (54; 55) showed a statistically significant reduction in the amplitude of spikes and slow waves with age. In epilepsy, increased IED amplitudes during sleep and after awakening have been anecdotally reported in benign epilepsy with centrotemporal spikes (BECTS) (56; 57; 58; 59). (60) later observed that the highest spike amplitude and shortest spike duration occurred within 2 cm from the site of seizure origin in 24 (75%) and 22 (69%) out of 32 patients, respectively. (39) and (32) reported that during REM spikes were often of lower voltage than those recorded during wakefulness or NREM sleep. (39) reported fluctuations in the sharpness and amplitude of IEDs during NREM, even within the same 30 seconds epoch. However, neither (32) nor (39) provided quantification or statistical comparisons between sleep stages. (61) reported a statistically significant increase in spike amplitudes in NREM versus REM sleep. (62) reported a statistical increase in the amplitude of cortical and mesial temporal spikes during N2 versus wakefulness, but these findings were based on a very limited number of observations. Recently, (35) investigated differences in spike morphology using a feature extraction method (63), reporting amplitude differences between brain regions and SOZ, but no statistically significant differences between sleep stages. A non-significant trend towards increased amplitudes in deeper stages of sleep could be observed, but only within median brain regions, and only from the SOZ. Together, while previous studies suggest that IED amplitudes are increased during deeper stages of sleep, consistent and statistically robust comparisons of IED amplitudes across all sleep stages have not yet been reported.

The current study investigated the influence of sleep on short and long-term dynamics of interictal activities, i.e. on time-locked neuronal activity during interictal events as well as on baseline activity without interictal events. For this purpose we analysed recordings from intracerebral electrodes containing both macro and micro contacts implanted in the hippocampal-amygdala complex of patients with focal epilepsy. An automatic template matching procedure was used to objectively identify IEDs in continuous long-term (2-3 weeks) recordings. Implanted micro electrodes allowed for the recording of neuronal action potentials during the full sleep and wake cycle. Polysomnography allowed detailed multilevel analyses during different stages of sleep.

We expected deeper stages of NREM sleep to increase the rate of IEDs, as well as their amplitude, while increasing baseline firing rates and bursting propensity. During the IEDs, we expected to find increased neuronal firing rates during IED spikes, and decreased neuronal firing rates during subsequent slow waves. Importantly, we expected these firing rate modulations during IEDs to increase with deeper stages of NREM sleep compared to REM and wake. Finally, we explored the effect of sleep on the regularity of neuronal firing with CV2 (64).

## Results

### IED detection

Manual annotation of IEDs resulted in an average detection of 5667 IEDs per patient (2106 to 12912) during the first 24 hours (Table S3). Automatic detection resulted in a similar average of 5618 IED detections per patient (3054 to 12811) in the same 24 hours, showing an average hit-rate of 98.0%, and an average false-alarm (false positive) rate of 6.5%. The total continuous recording time was on average 314 hours, i.e., just over 13 days, in which the automatic procedure detected an average of 50271 IEDs per patient (9224 to 113832, Table S3). In all patients, the six templates showed morphologies consistent with spike-wave interictal discharges (65; 66; 42; 48), with only modest variations in the amplitude of sharp peaks and slow waves (Figure 1).

**Fig. 1.**
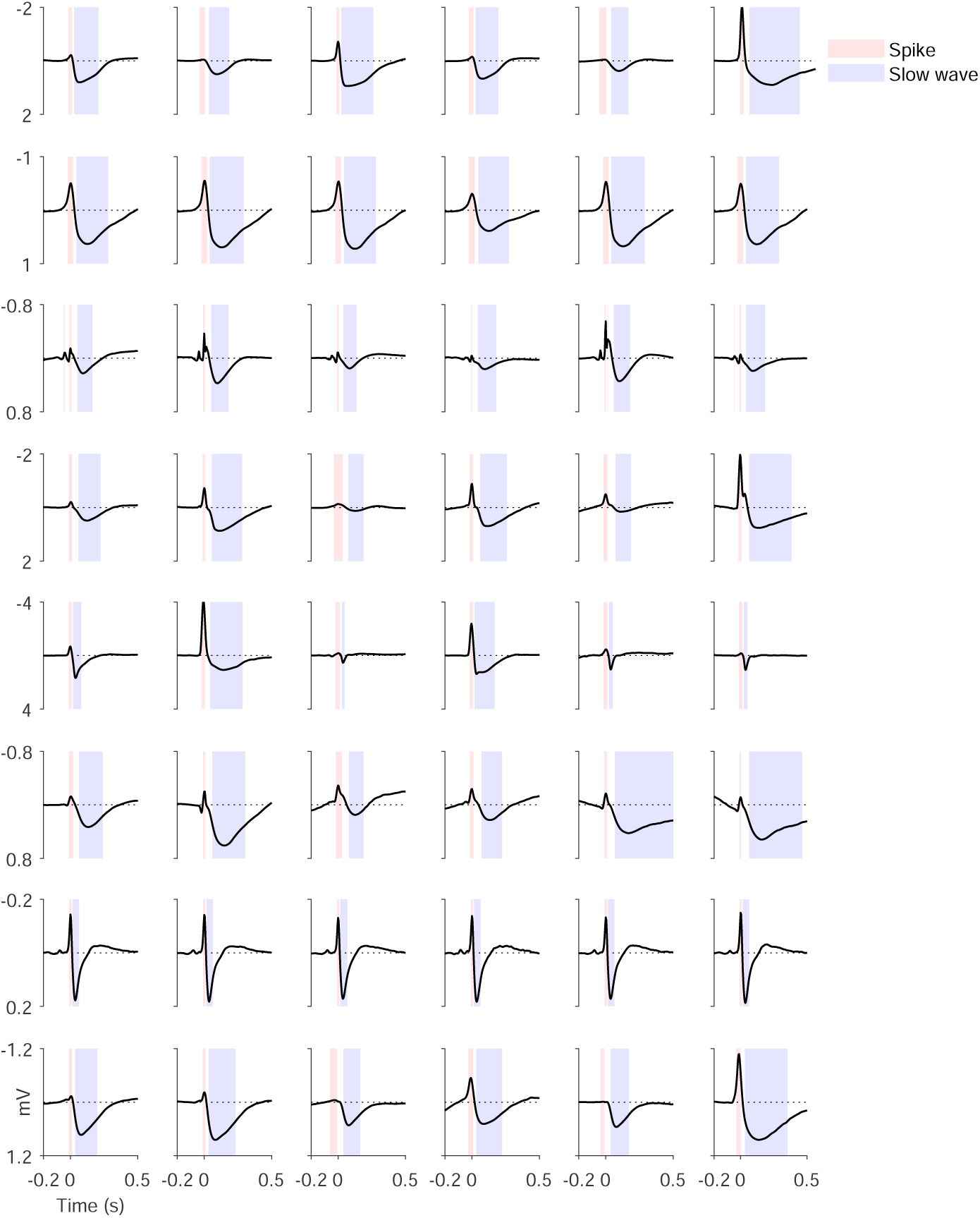
Templates for automatic detection of IEDs and statistical time intervals. Each row represents the six templates of one patient (from top to bottom: Patient 1 to 8). Black lines show the template at the channel with the largest response over templates. Red overlays show the spike period [*<* 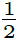 *min*(*V*)], blue overlays the slow wave period [*>* 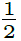 *max*(*V*)]. Note the clinical direction of the current (top = negative).

### Circadian distribution of interictal and ictal activity

IED rates showed a significant deviation from a uniform circadian distribution in all patients (Figure 2A & S5). Seven out of eight patients show a median phase between 01:00 and 03:30 at night, and the remaining patient presented a median phase at 17:25.

**Fig. 2.**
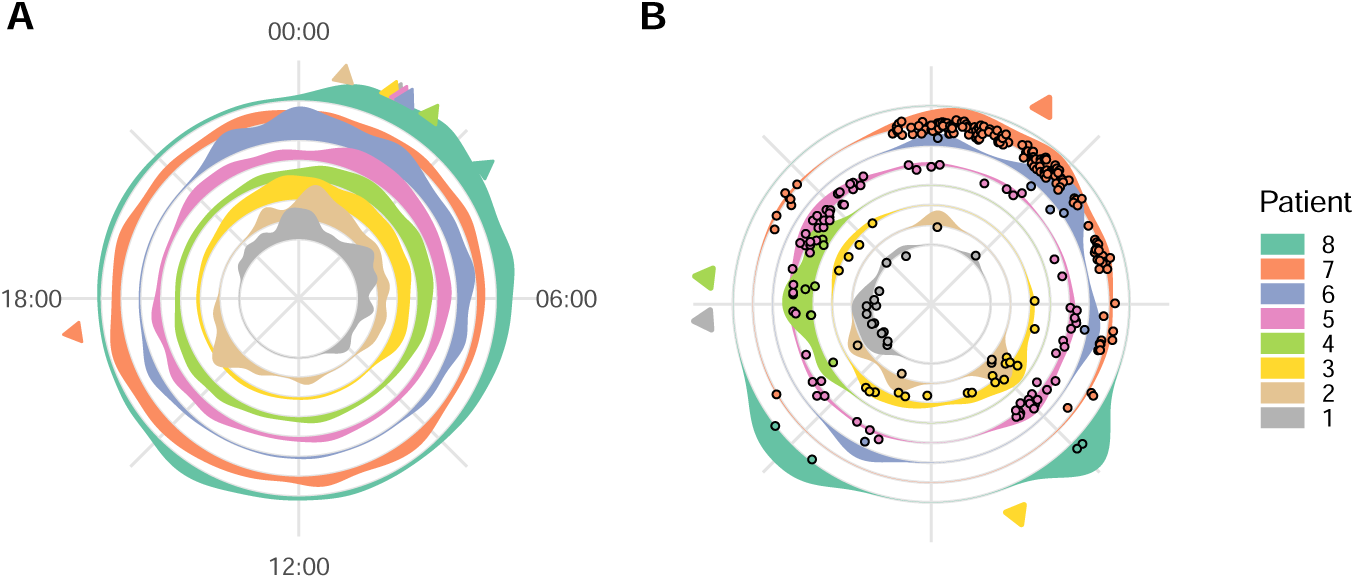
Polar representations of 24 hours circadian distributions of (**A**) interictal activity, (**B**) seizures. Significant mean angles of interictal density and seizure occurrence are shown by arrows. See Table S5 for details

Seizure occurrence showed a significant deviation from a uniform circadian distribution in four of eight patients (Figure 2B & Figure S5). In contrast to IED rates, the median phases of seizures were distributed throughout the day/night cycle. Of those patients who’s phase differed significantly from a flat distribution, the medium phase was found at 17:43, 10:31, 18:26 & 01:59.

### Increased IED rate with deeper stages of NREM sleep

IED rates increased significantly from wake to sleep (Figure 3A,B & S7), showing higher IED rates with increasing depth of NREM sleep (*S*3 *> S*2 *> S*1 *> Wake*). REM and wake did not differ significantly in IED rates.

**Fig. 3.**
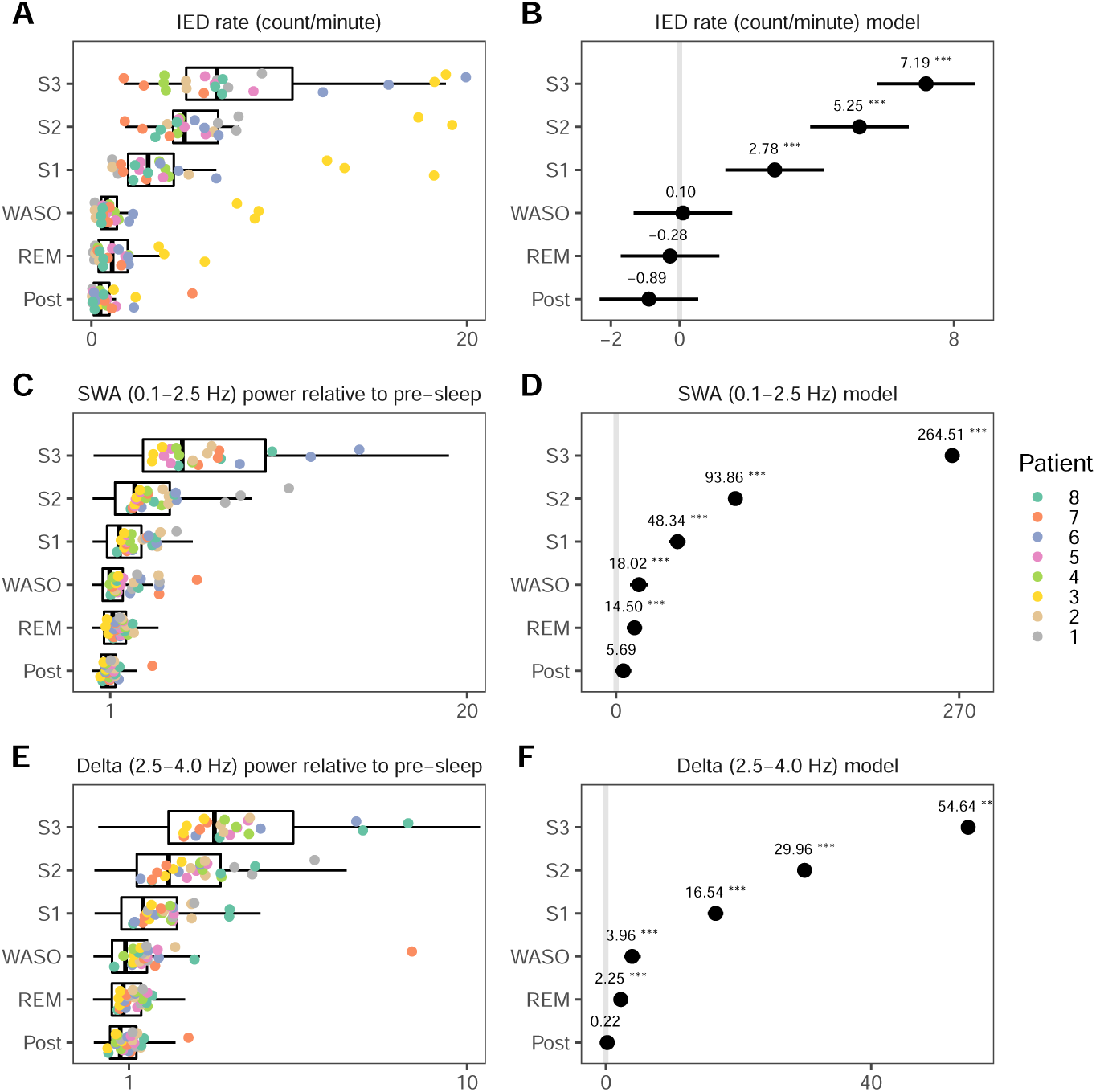
Summary statistics of IED rate, slow wave activity (SWA) and delta power per sleep stage, with box plots overlaid by values representing averages per patient (colored) and night. **A** and **B** show the rate of IEDs relative to the Pre-sleep period, and the fixed effects estimations, respectively. See Table S7 for statistical tests. **C** and **D** show the power in the SWA band (0.1 Hz to 2.5 Hz) relative to the pre-sleep period, and the fixed effects estimations, respectively. See Table S9 for statistical tests. **E** and **F** show the power in the delta band (2.5 Hz to 4 Hz) relative to the pre-sleep period, and the fixed effects estimations, respectively. See Table S10 for statistical tests.

### Increased IED rate with increased slow wave activity and delta power

For each patient, LFP data were epoched in 10-second non-overlapping intervals for the whole, and labeled according to the sleep stage, while excluding those that overlapped with IEDs or artefacts. Power in 0.1 Hz to 5 Hz was calculated in steps of 0.1 Hz, then averaged for each sleep stage, as well as over all sleep stages. Power at each frequency was then calculated relative to the pre-sleep period. Relative power showed a peak in the SWA frequency range (0.1 Hz to 2.5 Hz) which increased from 0.7 Hz in S3, to 0.8 Hz in S2, 0.9 Hz in S1, and 1.2 Hz in REM sleep (Figure S1). Mixed effects analyses verified that variance in both SWA and delta power were significantly explained by sleep stage (Figure 3C-F, Table S9 & S10), with post-hoc comparisons confirming that both SWA and delta power increased with depth of sleep (*Pre < REM < S*1 *< S*2 *< S*3). Variations in IED rate were significantly explained by power in both the SWA (0.1 Hz to 2.5 Hz), and delta range (2.5 Hz to 4 Hz) (Figure S2 & Table S8).

### Increased IED amplitudes with deeper stages of NREM sleep

Figure 4 shows the LFP of all detected IEDs, averaged per sleep stage, showing that the magnitude of both the peak and the slow wave components increased with decreasing stages of NREM sleep, in all but one patient (patient 7).

**Fig. 4.**
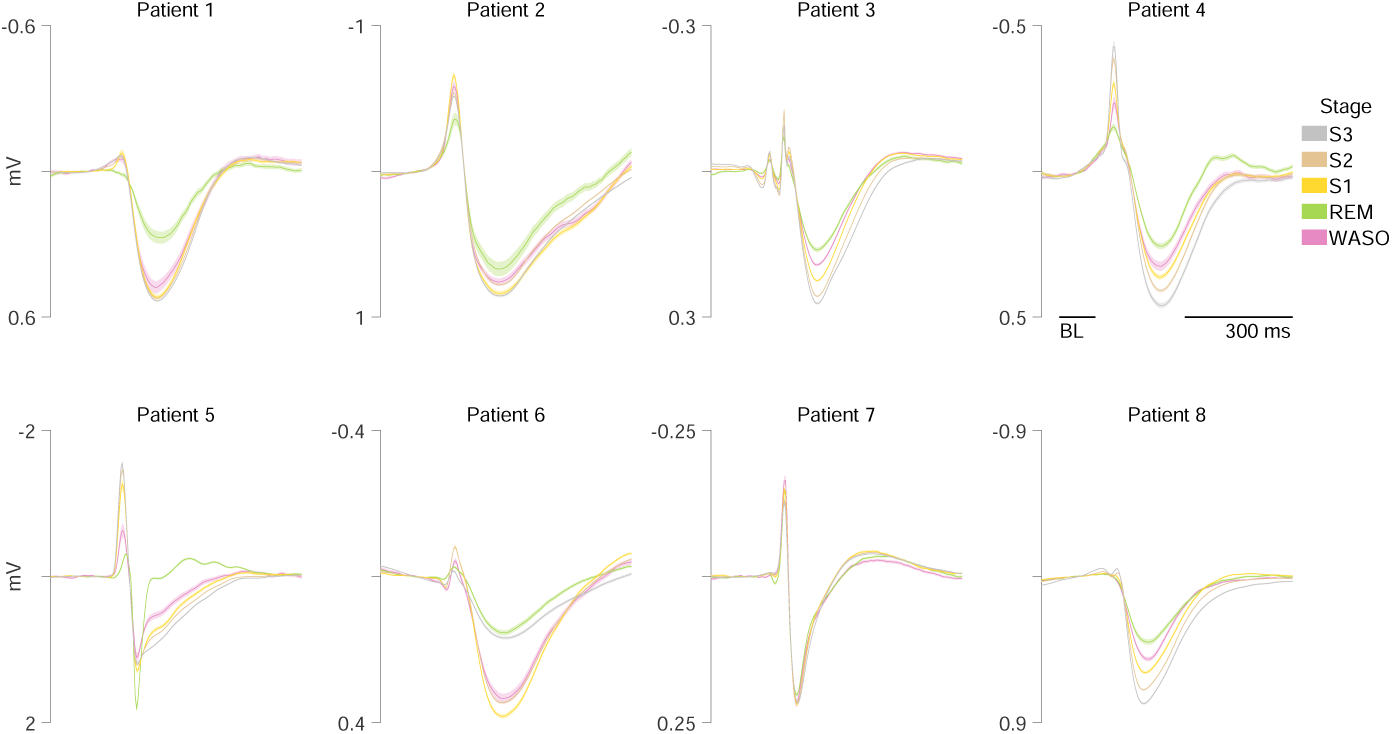
Average IED LFP for each patient (panel) and sleep stage (color).

To statistically test the amplitude modulation of the spike and slow wave components of the IEDs, the average amplitude during their respective intervals, as determined earlier (Figure 1), were entered into a mixed-effects analysis with patient and template as a random factor, and pre-sleep as the reference. Post-hoc comparisons allowed the evaluation of differences between sleep stages. The amplitudes of the IED spikes were increased in SWS compared to pre-sleep, WASO and REM, and increased progressively with increased depth within SWS (*S*3 *> S*2 *> S*1, Figure 5A,B & Figure S12). Slow wave amplitudes decrease further (i.e. increased the deflection) during SWS compared to WASO and REM sleep, but were not significantly different within SWS stages. (Figure 5C,D). When comparing the difference in peak and slow wave amplitudes (*V ^peak^ − V ^SW^*), a clear gradual increase with decreasing arousal could be seen (Figure 5E,F), with post-hoc tests showing a gradual increase of spike-wave deflection with increasing depth of sleep (*S*3 *> S*2 *> S*1). Neither the spike, wave or difference, was different between pre-sleep and REM.

**Fig. 5.**
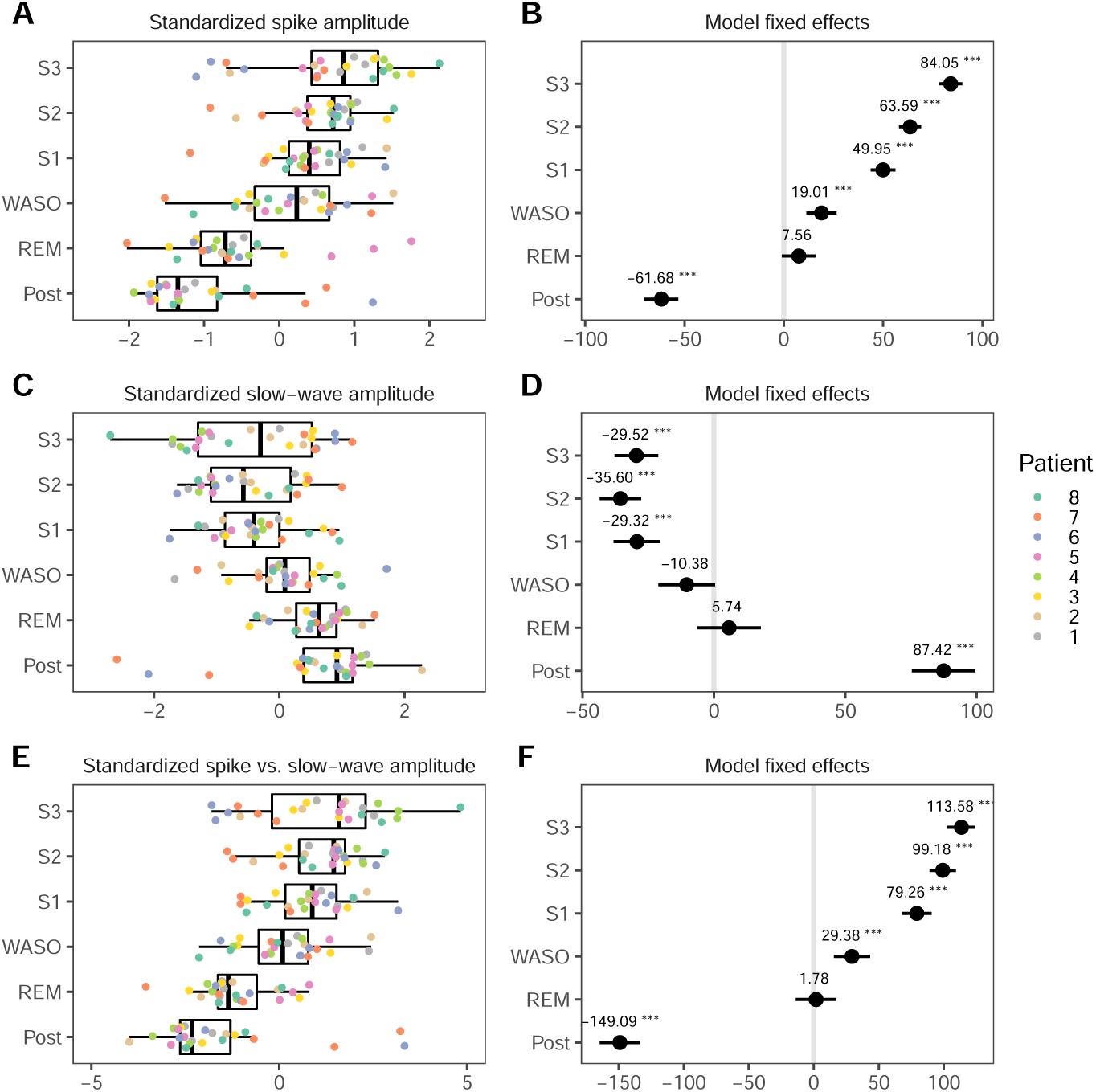
Summary statistics of IED amplitude per sleep stage, with box plots overlaid by values representing averages per night, per patient (colored). **A** & **B** shows the average amplitude during the IED spike relative to those during the pre-sleep period, and the fixed effects estimations of the comparisons, respectively. **C** & **D** shows the average amplitude during the IED slow wave relative to those during the pre-sleep period, and the fixed effects estimations of the comparisons, respectively. **E** & **F** show the total deflection of the spike and wave (*V ^peak^ − V ^SW^*) relative to the pre-sleep period, and the fixed effects estimations, respectively. See Table S12 for a summary of statistics.

### Firing rate modulation during IEDs increases deeper stages of NREM sleep

Over all patients and nights, 112 putative single units (SUA), and 131 multiunits (MUA) were extracted (Table S11). In all patients, individual peri-stimulus time histograms (PSTHs) showed increases in firingrates time-locked to the IED spike, as well as more gradual and sustained decreases during the subsequent slow wave period (Figure S12-S21). Some units only responded with an increase during the spike, while some responded with only a decrease during the slow wave. Many units showed both, however, and the average response showed a strong modulation consistent with the LFP (Figure 6). Together, the majority of units were shown to be responsive to IEDs, i.e. showed significant changes in firing rates during IEDs versus baseline (SUA: 75.9%, MUA: 65.6%, Table S11).

**Fig. 6.**
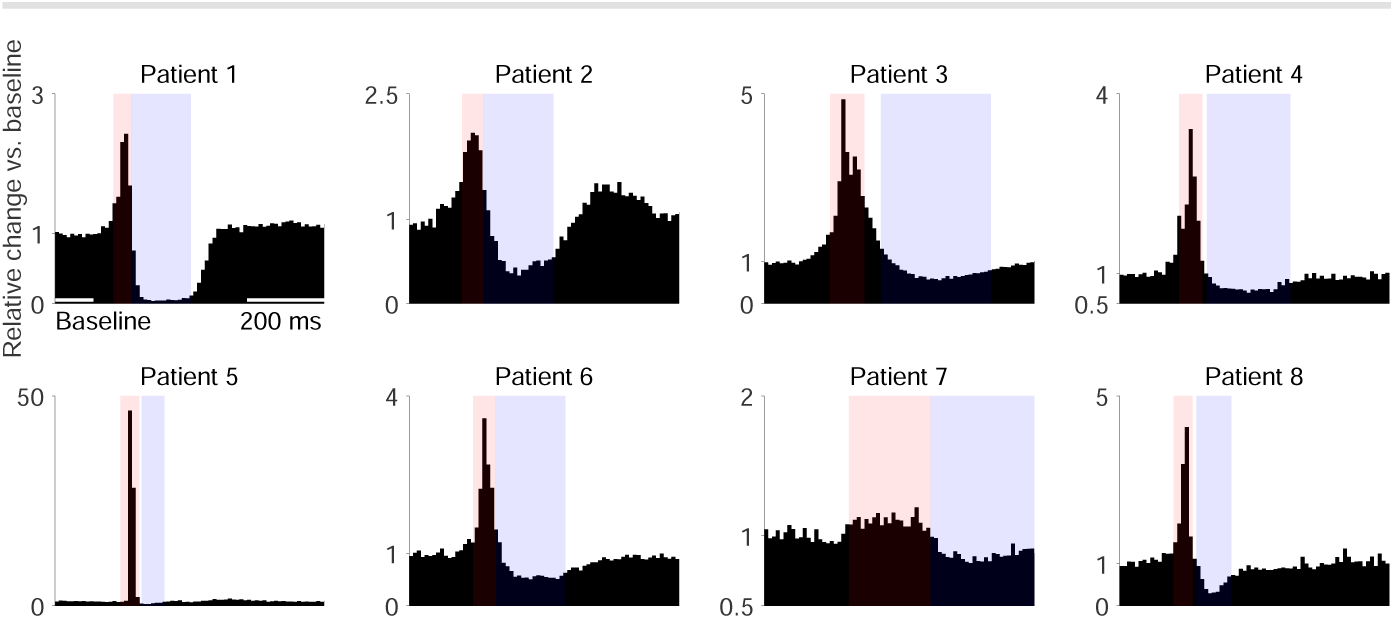
Peri-stimulus time histograms (PSTH) of all responsive units, templates and sleep stages, separately for each patient, including the average LFP showing the consistency between the PSTH and the LFP. For each patient, template and unit, firing rates were standardized by dividing the PSTH by the mean firing rate during baseline (indicated in the first panel). Red and blue overlays indicate intervals for statistical tests of the sharp wave and slow wave, based on the width at half-prominence (Figure 7 & Table S13).

To statistically test the effect of sleep stages on firing rate modulation during IEDs, the average firing rates of each unit during the spike or slow wave periods (Figure 6) were entered into a mixed model. Firing rates during IED spikes increased significantly during NREM sleep (Figure 7A&B, Figure S13) compared to the period before and after sleep. Firing rates during IED spikes were also slightly increased during wake periods sleep onset (WASO). No differences were found between NREM and REM sleep, nor between REM and wake periods. During the subsequent slow wave, NREM (as well as WASO) also significantly decreased firing rates further, compared to waking periods (Figure 7C & D). REM was not found to affect firing rates during either the peak or slow wave. Finally, the normalized difference in firing rates between peak and slow waves (Figure 7E&F) summarized the effect of NREM sleep on firing rate modulation during IEDs. During NREM sleep, as well as during WASO, the firing rate difference between peak and slow wave were increased. None of comparisons showed differences between S1, S2 and S3, and in all comparisons REM sleep did not differ significantly from waking periods.

**Fig. 7.**
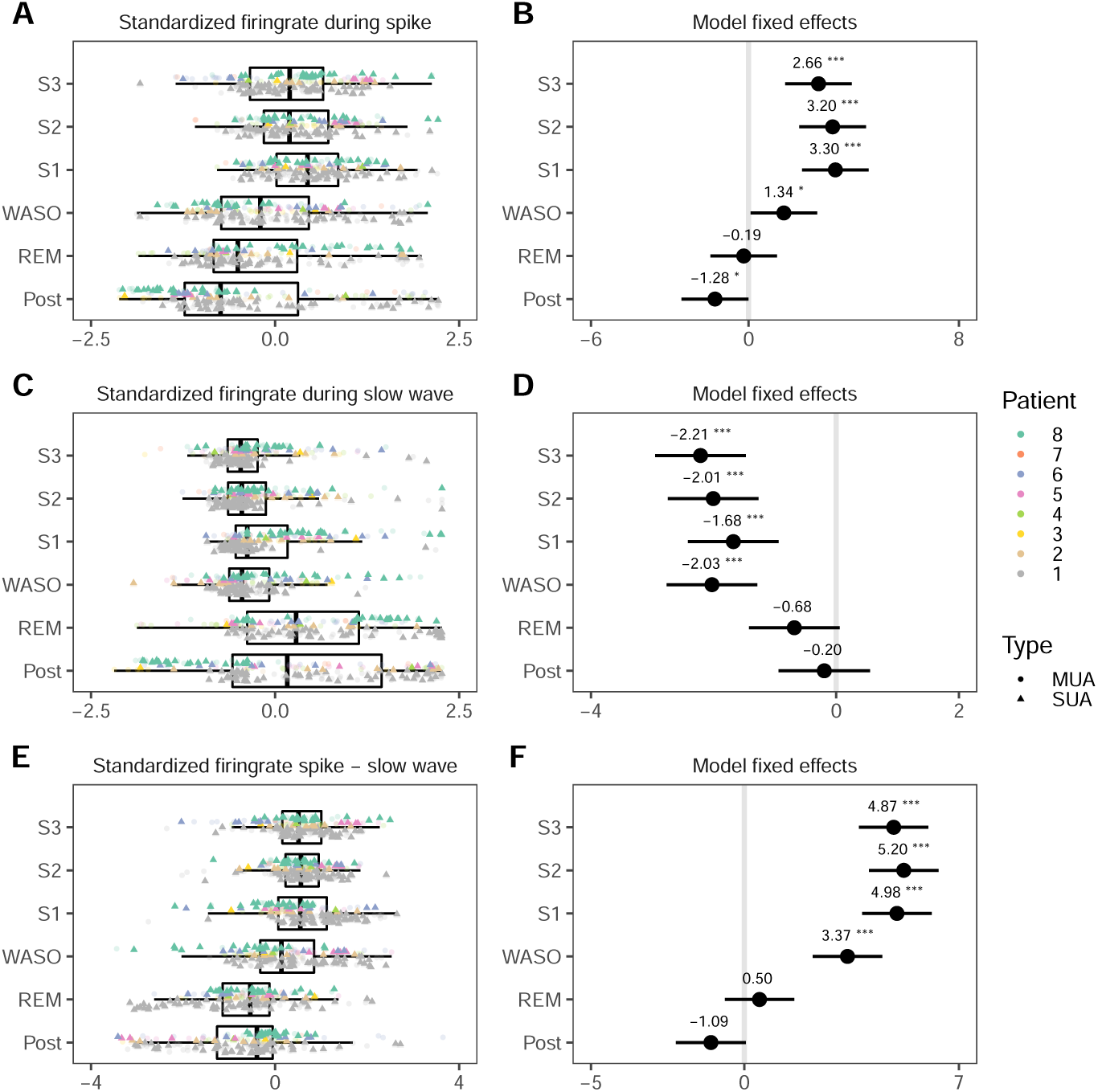
Unit firing rates during the spike and slow wave periods of the interictal discharges. **A**, **C** & **E** show box plots overlaid by values representing averages per patient (colored) and unit. Pyramid shapes and dots represent SUA and MUA, respectively. For display purposes, values were normalized over all sleep stages. **B**, **D** & **F** show model estimates of fixed effects, including annotations for significant differences versus pre-sleep (*p <* 0.05., *p <* 0.01*^∗^*, *p <* 0.001*^∗∗^*, *p <* 0.0001*^∗∗∗^*). See Table S13 for a summary of the statistics.

### Unit resting behavior changes with deeper stages of NREM sleep

Resting baseline activity of units, i.e. during periods in which no IEDs occurred, was investigated by epoching the first 72 hours in 10-second non-overlapping intervals. These epochs were labeled according to the stage of sleep in which they occurred, and excluded if they overlapped with IEDs or artefacts.

Mixed models showed that SUA and MUA firing rates, as well as bursting rates decreased significantly with increasing depth of sleep, with post-hoc tests verifying a progressive increase (*pre > REM > S*1 *> S*2 *> S*3, Figure 8A-F & Figure S14). CV2 also increased significantly, in an almost linear fashion until a large increase during S3 (*S*3 *>> S*2 *> S*1 *> Wake > REM*), indicating decreasing regularity of action potential firing with decreasing depth of sleep (Figure 9A & B). Finally, the amplitudes of the action potential increased significantly during all stages of sleep compared to both the pre and post period (Figure 9C & D), in which post-hoc comparisons showed a progressive increase wioth *S*3 *> S*2 *> S*1.

**Fig. 8.**
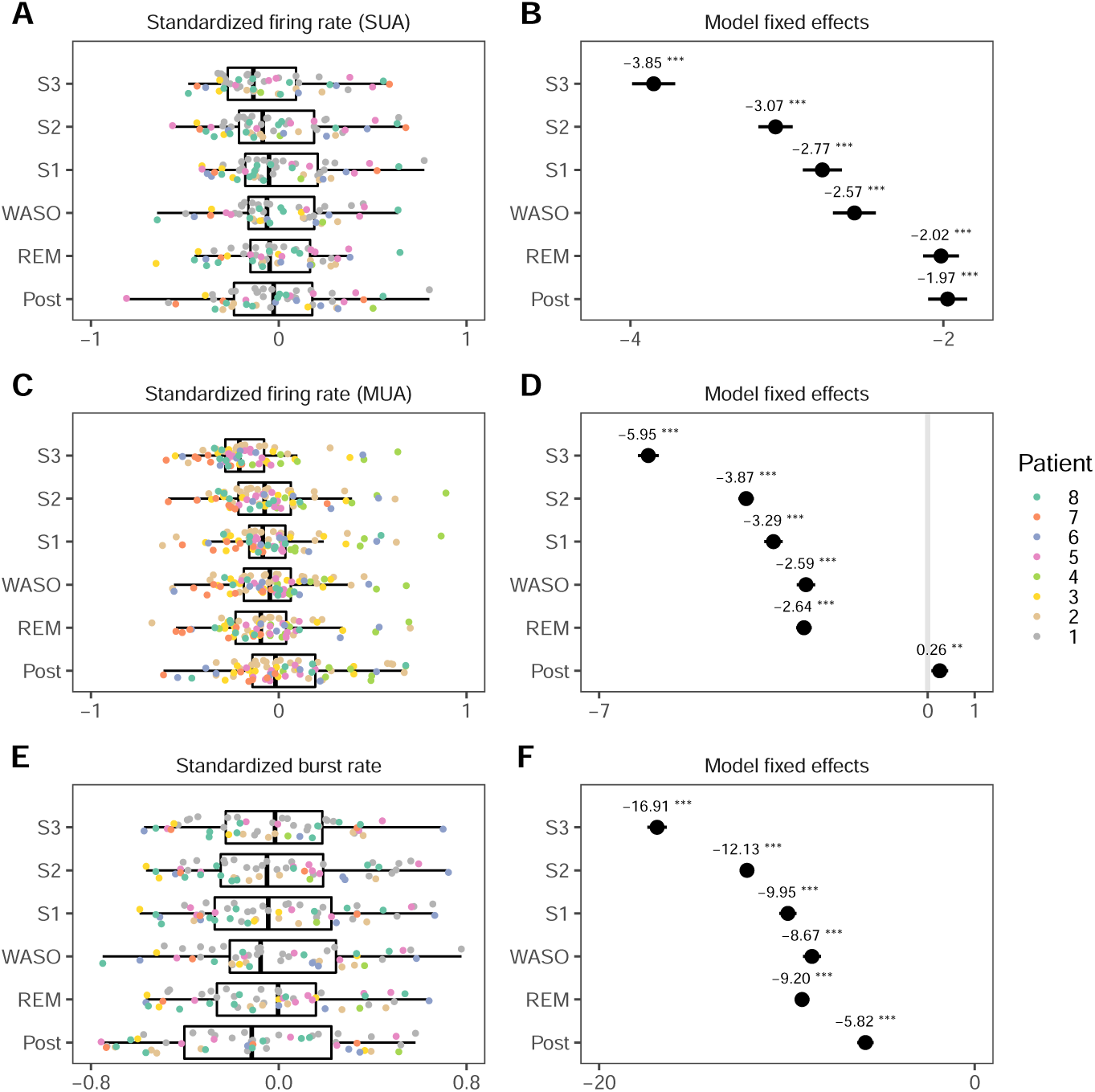
Summary statistics of average unit characteristics during rest, in different sleep stages. **A**, **C** & **E** show box plots overlaid by values representing averages per patient (colored) and unit. For display purposes, values were standardized as relative change versus pre-sleep according to 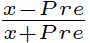). **B**, **D** & **F** show model estimates of fixed effects, including annotations for significant differences versus pre-sleep (*p <* 0.05., *p <* 0.01*^∗^*, *p <* 0.001*^∗∗^*, *p <* 0.0001*^∗∗∗^*). Only SUA were included in the calculation of burst rates. See S14 for a summary of statistics.

**Fig. 9.**
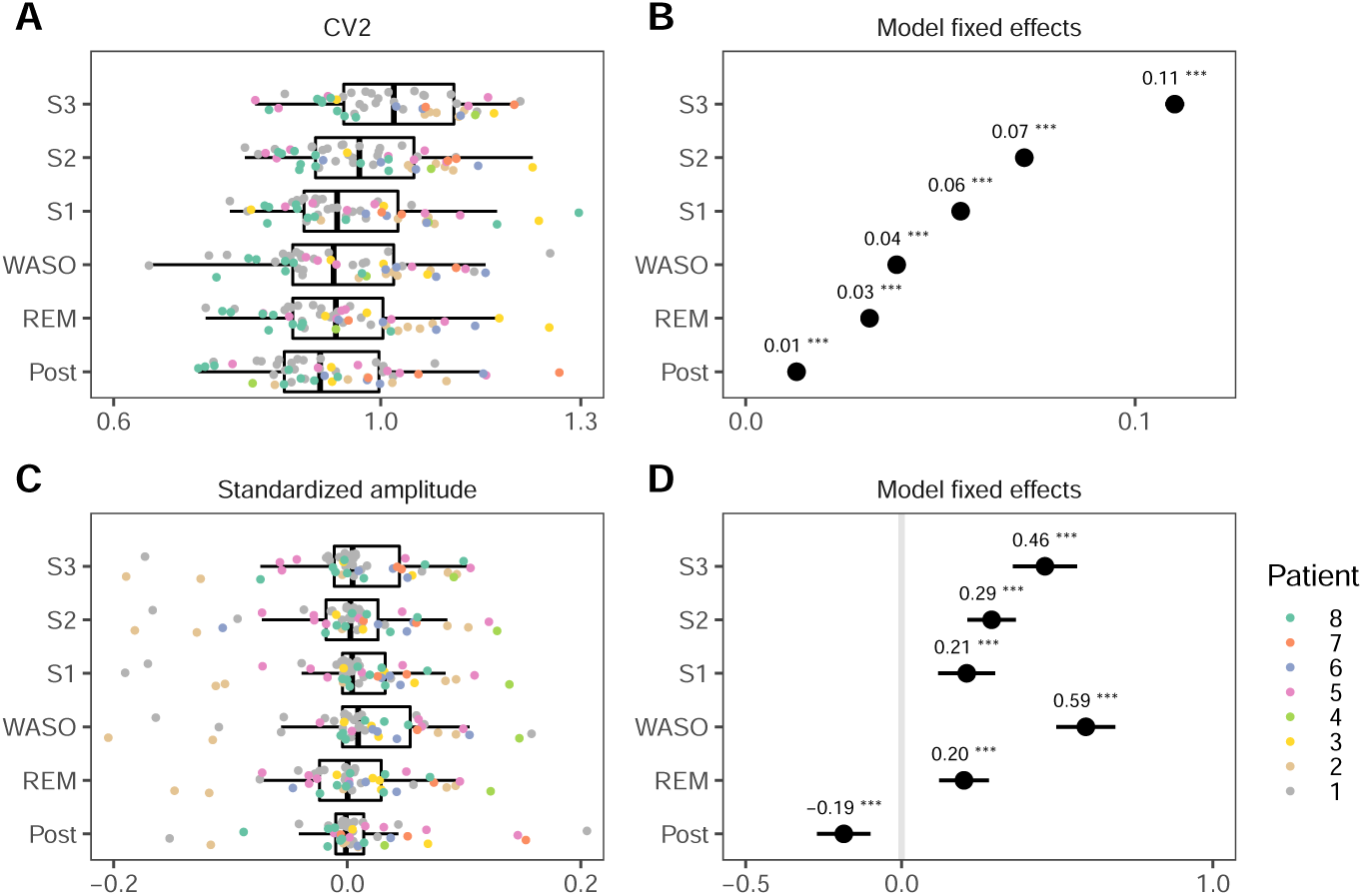
Summary statistics of average unit characteristics during rest, for different sleep stages. **A** & **C** show box plots overlaid by values representing averages per patient (colored) and unit. For display purposes, values were standardized as relative change versus pre-sleep: 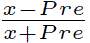). **B** & **D** show model estimates of fixed effects, including annotations for significant differences versus pre-sleep (*p <* 0.05., *p <* 0.01*^∗^*, *p <* 0.001*^∗∗^*, *p <* 0.0001*^∗∗∗^*). Only SUA were included. See Table S14 for a summary of statistics.

**Fig. 10.**
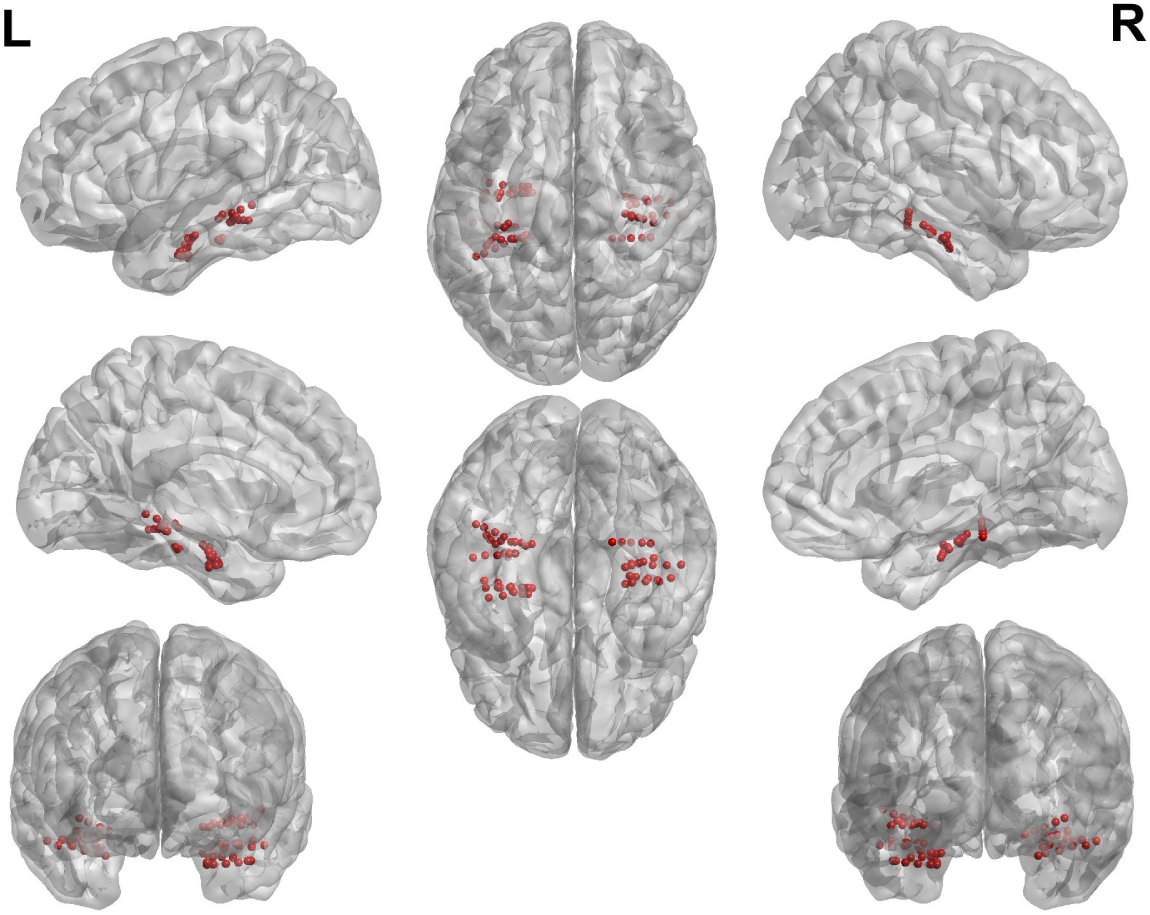
Locations of each analysed electrode contact, transformed to MNI space. See S4 for MNI coordinates.

**Fig. 11.**
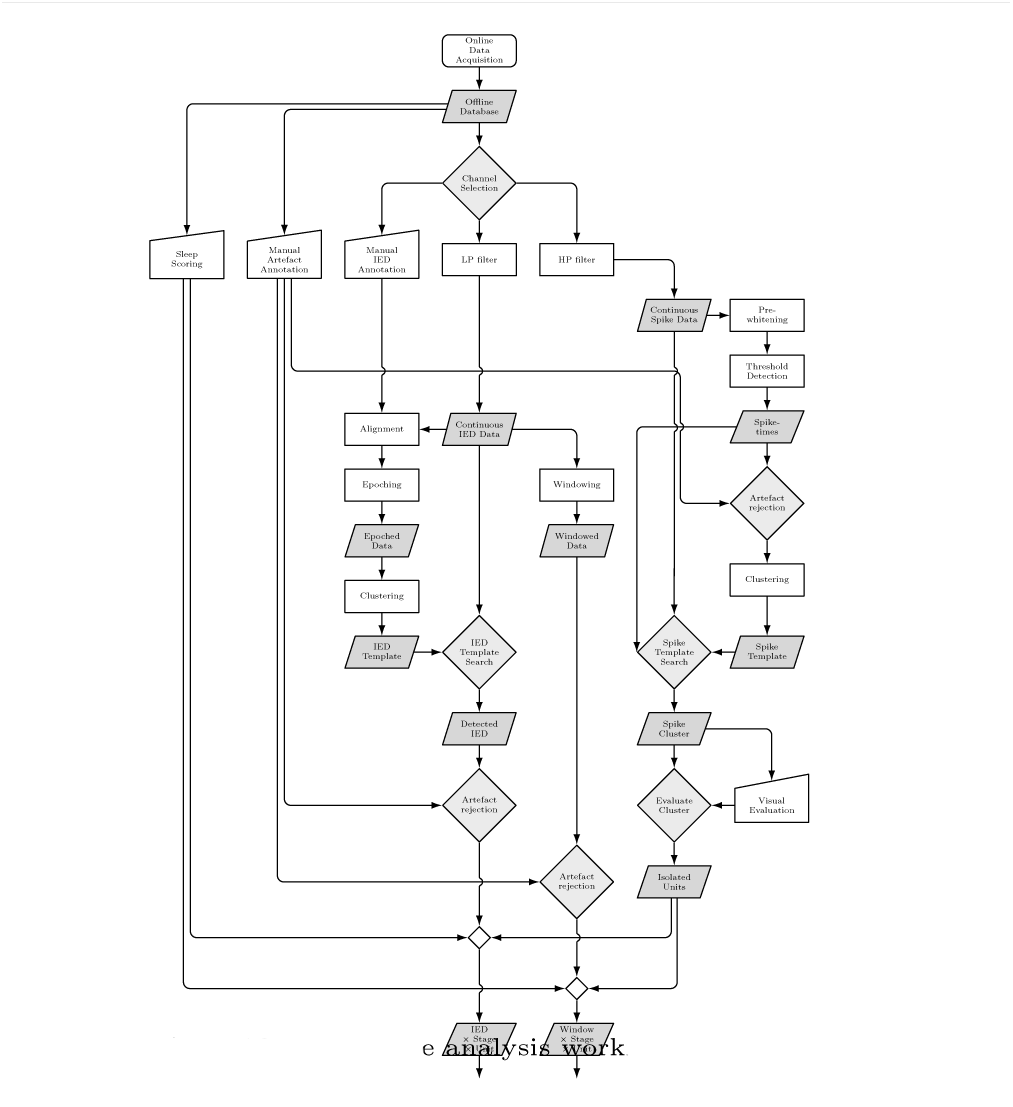
Visual depiction of the LFP and spike analysis workflow. See the Methods section for details.

## Discussion

### Interictal rate is under circadian influence

All eight patients showed a circadian distribution of IED rates, with seven peaking between 01:00 and 03:30 at night, consistent with previous reports (11:00-05:00 in (36), 02:21-04:12 in (33) and one of the clusters at 04:00, 02:00, and 06:00 in (37)). While our population was heterogeneous with regard to the seizure onset zone (SOZ), including prefrontal, temporal, multifocal, and amydalo-hippocampal regions, all showed IEDs in mesial structures. Only one patient, with a SOZ in the left parietal region, did not show a nightly peak in IED rates from mesial structures. Although the sample size of our study prevent conclusions on the effect of the SOZ, previous studies have shown that the seizure onset zone (SOZ) and anatomical location do not effect circadian influences on interictal activity (26; 31; 32; 12; 19; 67; 22; 68; 69), but see (34).

### Seizure occurrence is under circadian influence

Several patients showed a circadian influence on the occurrence of seizures. However, peaks of seizure occurrence were spread out along the day/night cycle. Patient 1, 3, 4 & 7 showed a circadian peak in the early evening, late morning, early evening, and night, respectively. This did not seem to be associated with a particular SOZ, which were frontal, temporal, temporal and parietal, respectively. Patient 5, and possibly Patient 3, showed a bimodal distribution, with a peak in the morning and evening, and a multifocal and temporal SOZ, respectively. The number of seizures of Patient 2, 6 & 8 were not sufficient to identify any potential circadian pattern. These results are in line with previous literature, showing circadian peaks in seizure occurrence (36; 70; 71; 72; 73; 74; 75) with frontal lobe seizures generally peaking very early in the morning, and mesial temporal lobe seizures showing a more bimodal distribution, with a primary peak in the late afternoon and a secondary peak in the morning (76; 77; 70; 73; 72; 74; 78; 33). The exact timing of the peaks are known to vary considerably between studies and patients, likely due to inconsistencies in protocols and analyses, as well as factors such as samples size, sample demographic, geographic location, seizure focus, and individual differences in circadian rhythms (79; 67; 80). Our results are consistent with varied circadian patterns in seizure occurrence, showing that temporal seizures display single and bimodal distributions throughout the day/night cycle, although the fact that our patient population had heterogeneous SOZ location might also have contributed to the variation in circadian distributions.

### IED rate and amplitude are modulated by sleep stage

Consistent with previous studies, the IED rate increased progressively during NREM sleep (*S*3 *> S*2 *> S*1 *> Wake*) (26; 31; 32; 27; 28; 30; 33; 34) and no differences in IED rates between REM and wake (32), although the latter has been occasionally reported (38; 31). Consistent with previous reports (39; 27; 40), we found that IED rates were positively correlated with power in the slow wave (0.1 Hz to 2.5 Hz) and delta (1 Hz to 4 Hz) frequency range.

The increase in IED rates during SWS and their suppression during REM sleep has been hypothesized to be due to increased brain synchronization during SWS and decreased synchronization during REM sleep (49; 50). In fact, the same networks of cortical neurons involved in synchronizing thalamically generated delta oscillations during SWS may serve as the preferential substrate for epileptiform spike-wave activity (81; 82; 83; 84). Several studies reported an increase in the spatial distribution of IEDs during SWS and a decrease during REM sleep (32; 50; 10; 85; 13; 62; 86; 87; 88; 49; 89; 90). However, differences in IED amplitudes were rarely reported and in none of the aforementioned studies. An increase in IED amplitudes could result in increases in electrical field spread, which could be misinterpreted as an increase in spike propagation. Furthermore, the extent of the reported changes in the spatial distribution has generally been small and not unequivocally found.

For example, (13) found that while interictal activity shifted from anterior to posterior and from mesial to lateral during sleep, the source of interictal activity remained localized within the amygdalo-hippocampal complex. (91) found no effects of sleep on the spatial distribution, and (89) found spatial distribution not to be predictive of seizure occurrence. Finally, (34) found an increase in co-occurrence of different interictal sources during SWS, but only for neocortical spikes, not mesial-temporal spikes. Together, the reported changes in the spatial distribution of IEDs are small and inconsistent, while robust studies of the effect of sleep on the amplitude of IEDs have been missing. The current study analyzed an unprecedented number of IEDs (3500 to 12811 per patient) from continuous data recorded over three day/night cycles and found clear evidence of an increase in IED amplitude during sleep. The LFP deflections of both the spike and slow wave were found to be gradually increased with deeper stages of sleep.

### Interictal activity modulates firing rates of the majority of neurons

Early studies found that some neocortical neurons discharged in time with interictal spikes (92). These have been termed “positive” (93) or “involved” neurons (94), and were considered to reflect the intensity of epileptic activity. More recent studies have consistently found that a proportion of units increase their firing rate during the interictal spike, which is then reduced during the subsequent slow wave (95; 43; 48; 47; 46).

Our results show a consistent pattern across all patients. Units increased their firing rate sharply during, or right before, the upward flank of the sharp LFP spike, then reduced their firing rates to below baseline, highly consistent with the shape of the LFP slow wave. Only one patient (Patient 7) did not show fast and strong firing rate modulation during the IED spike. However, in the rest of the patients, the majority of units showed significant changes in firing rates versus baseline (SUA: 75.9%, MUA: 65.5%). This high proportion of “involved” units is somewhat higher than the 49% found by (95), the 48% by (47), and the 40% by (46). However, methods to determine whether a unit modulates its firing rate during IEDs depend on the sensitivity of the design and statistical methods. The early work by (95) was visually performed based on a relatively small number of IEDs, the number of which were not reported but indicated to be less than 50 in the representative figures. (47) removed sleep periods in their analysis and reported a large variability in the number of manually annotated IEDs per patient (31–608). In the current study, IEDs were automatically detected on three days, resulting in 2106 to 12912 IEDs *per patient*. While it stands to reason that the increased number of data points increased the sensitivity, clinical differences could also explain the higher rate of responsive neurons. In (95) and (46) clinical information was not provided. In (47), 269 out of 363 units were recorded from dysplasic cortex, of which 26 had a cryptogenic origin, and the remaining 73 units came from patients with various lesions. Furthermore, different recording methods were used to record LFP and spike data from diverse regions and with a large variation in the distance to the SOZ. Although our population also showed heterogeneous SOZs, recordings of LFP and spike data were all restricted to the hippocampal-amygala complex. The hippocampal formation has anatomical and physiological mechanisms that promote neuronal synchronization, making this region very prone to epileptic discharges. It should be noted that using a similarly sensitive statistical approach in a recent study on periventricular nodular heterotopia, all units were shown to respond to interictal events (96). Together, our findings are consistent with the paroxysmal depolarizing shift (PDS) mechanism in which a large depolarization phase is followed by a long hyperpolarization found in animal (44; 45), and consistent with *in vitro* studies on hippocampal slices from human patients with temporal lobe epilepsy (97; 98; 47).

### Firing rates modulation increases with deeper stages of NREM sleep

While modulation of firing rates during IEDs has been established and replicated here, it is unknown whether this modulation is under the influence of sleep. Our results show for the first time that the firing rate during the sharp LFP peak is significantly increased during sleep compared to waking periods, while firing rates during the subsequent slow wave are significantly reduced.

The first part of the PDS depolarization phase is believed to be generated by intrinsic membrane conductances (99), and the later from feedback recurrent synaptic excitation mediated by AMPA and NMDA receptor subtypes, and glutamate receptor-coupled calcium conductances (100). The subsequent hyperpolarization represents GABA-mediated recurrent inhibition, as well as Ca2+-dependent outwards K+ currents. Our results suggest that sleep increases excitability and synchronization of neuronal populations, increasing both the probability of the occurrence of IEDs, as well the degree of de-facilitation by subsequent hyperpolarization.

### Neuronal baseline activity is modulated by sleep stages

Neuronal baseline activity, i.e. resting behaviour in the absence of IEDs, was investigated by means of firing rates, burst rates and CV2. These were calculated from a large number of non-overlapping 10-second time windows in 72 hours, allowing robust estimations of conditional means within patients. By using mixed models, we dealt with both between-subject and within-subject variance. Firing rates, burst rates and CV2 were all shown to be lower under increasing depth of sleep, and especially pronounced during SWS. A reduction in the firing rate during SWS is consistent with the periodic suppression of neuronal firing due to SWA during NREM sleep, as shown in animal *in vivo* intracellular recordings (51; 53). Although measures of regularity were not reported in these studies, decreased regularity with decreased depth of sleep could be explained by such “clumping together” (51) of action potentials.

These studies, as well as our own findings, are in partial disagreement with (52), who reported a reduced firing rate in REM versus wake and SWS, a higher propensity for bursting during SWS versus wake and NREM, and increased variability (CV2) in SWS versus wake. However, these analyses were performed on only 23 neurons and, as the authors themselves noted, potential influences from interictal discharges were not considered. Another study of the same lab included a larger number of isolated neurons (72), and reported higher burst rates in MTL regions, but only during sleep (SWS and REM), and lacking consistent difference in firing rates between sleep stages or between epileptic versus non-epileptic regions (101).

Although early human studies found indications of bursting in epileptic regions (102; 103), others found no differences in firing patterns such as bursting between neurons in the epileptic focus or surrounding area (92). In fact, while some studies continued to associate bursting behavior with epilepsy in both animal (104) and human in vitro studies (105; 106), microelectrode recordings in epileptic patients have so far returned contradictory results, showing both increased (101; 107) and decreased (108) bursting in the SOZ of MTL patients.

### Automatic detection of IEDs

The current study exploited the possibility of analyzing a very large number of IEDs by using automatic detection of IEDs. Many algorithms have been proposed for IED detection over the last decades (109; 110; 111; 112), with recent implementations showing good, or better, performance than human experts (113; 114; 115; 116). Recent examples include the use of adaptive morphological filters (117), signal envelope distribution modeling (118; 119), convolutional neural networks (CNN) and deep learning (120; 121; 122; 123; 124; 125), long short-term memory (LSTM) neural networks (126) and generative adversarial networks (GANs) (127; 128). Two-step methods have been proposed to reduce the need to manually optimize parameters per different datasets (63; 129). Furthermore, machine learning models need to be trained on large standardized datasets of annotated data to deal with the challenge of generalizing models over different patients. Such datasets currently do not exist, but are necessary to objectively compare the performance of the available machine learning approaches (109; 111; 112). Furthermore, the inherent unbalanced nature of interictal data, i.e. the small ratio of positive events within large periods of background activity, has only recently been addressed (130).

The use of LFP templates provide a more robust and transparent alternative, and have been shown to outperform any classifier trained on morphological features (131). Template matching approaches have also been shown to perform very well even with high noise situations such as recordings taken from within an fMRI scanner (132). Some studies have used templates derived from databases (133), used patient-specific annotations (134), or performed clustering of patient-specific IEDs for rapid visual inspection and validation (124). In the current study, we chose to employ template matching based on patient-specific IED templates, extracted from visual annotations of 24 hours. The approach showed to be highly effective, while remaining relatively straightforward. Importantly, the templates can be visually evaluated, in contrast to “black-box” approaches using machine learning. By expanding the IEDs into six templates per patient, potential variations in the morphology of IEDs were taken into account as well.

### Conclusion

The current study evaluated for the first time the influence of sleep stages on neuronal firing during interictal activity, by means of *in vivo* recordings from hippocampal structures in epileptic patients. This was done with an unprecedented large dataset of continuous recordings of several nights, and thousands of IEDs per patient. The IED rate as well as amplitude were found to be increased with deeper stages of NREM sleep. Neuronal firing rates were found to increase during the IED spike, and decreased during the slow wave. Importantly, these firing rate modulations during IEDs were increased with deeper stages of NREM (SWS) sleep, versus REM and wake. Finally, during resting default behaviour, the neuronal firing rate, bursting rate and firing regularity were all shown to progressively decrease with deeper stages of NREM sleep. Together, this study provides the first evidence from human *in vivo* recordings, showing that sleep increases neuronal synchronization and subsequent de-facilitation during interictal epileptiform activity.

## Methods and Materials

### Patients

Eight drug-resistant focal epilepsy patients were selected from our database (135), with the following criteria: 1) Implantation of macroelectrodes targeting the mesial temporal structures, 2) microelectrode recordings with visible MUA, and 3) polysomnography recordings during the first three nights. See S1 for clinical details.

### SEEG recording

Patients were implanted with intracerebral depth electrodes (Ad-Tech^®^, Oak Creek, Wisconsin, USA), consisting of 4 to 8 platinum macroelectrode contacts, 2.41 mm long (Ø=1.1 mm). Each patient was also implanted with one to three Behnke-Fried type macro-micro electrodes (Ø=1.3 mm) that included eight additional microelectrodes and a reference wire of *≈* 2 mm long (Ø=40 µm), extending from the tip of the electrode shaft and capable of recording non-overlapping cell populations (136). Trajectories, anatomical targets, number, type and the number of contacts were all determined according to the clinical practice and the epilepsy features of the patients. Implantation was performed in the Department of Neurosurgery of the Pitié-Salpêtrière Hospital using a Leksell Model G stereotactic system (Elekta, Inc., Norcross, GA) or using a robotic assistant device (ROSA^®^ Brain, Medtech, France). For more details and evaluation of the implantation procedures, see (137) and (135).

The locations of the anatomical electrodes after implantation were determined by VF (S2), based on pre-implantation 3 T 3D-MRI, post-implantation 1.5 T 3D-MRI and post-implantation CT scan, combined using the EpiLoc plugin (138) for 3D-Slicer (139), developed by the STIM engineering facility at the Paris Brain Institute. Spatial locations of the electrodes were automatically computed in native space using the EpiLoc plugin, and visualized (10) using BrainNet Viewer (140), after transformation from to MNI-space (S4) using the Freesurfer image analysis suite integrated in Epiloc.

All patients gave their written informed consent (project C11-16 conducted by INSERM and approved by the local ethics committee, CPP Paris VI).

### Polysomnography

During long-term video-EEG monitoring, video-EEG and polysomnography (PSG) were simultaneously recorded for the first three nights post-implantation. When the first night was too disturbed, an extra night was scored and the first night was not analyzed. PSG included three EEG channels (Fp2, C4 & O2, or Fp1, C3 & O1, referenced to linked mastoids M1 or M2, respectively), right and left electro-oculography (EOG) channels, chin electromyography (EMG) and electrocardiography (ECG). Sleep scoring using PSG recording was performed by VH-NM, according to the American Academy of Sleep Medicine (AASM) 2017 criteria (141). Before sleep onset, and after the final awakening, a continuous period of two hours was labelled *pre-sleep* (or *pre*) and *post-sleep* (or *post*), respectively.

Sleep quality was varied among patients and nights (S6). However, all patients had a sufficient amount of sleep per night (average = 8 hrs., range = 6.5-11.9 hrs.). Light sleep (S1) somewhat increased above normal (15% vs. <5%), while deep sleep (average S3 = 20%) and REM (average = 19%) were normal (142). Hypnograms show a frequent sleep fragmentation (S3), which might be explained by multiple factors, including the post-surgery and hospitalized condition.

### Analysis

Analyses were performed with a combination of FieldTrip functions (143), custom MATLAB scripts (Version 2020b, The Mathworks Inc., Natick, Massachusetts). All analysis scripts are available here.

### Artefact detection

In the first three days (72 hours) of continuous recording, periods with movement artefacts or instrumentation noise in macro-or microelectrode recordings were manually annotated using software developed in-house (MUSE) for synchronous visualization of macro– and microelectrode signals (135). Data from one night (second) of a patient (patient 3) was replaced by a fourth night, due to excessive artefacts, and the polysomnography was then performed on that night as well.

### IED detection

IEDs are fast (<200 ms) high-amplitude (>50 mV) EEG transients, defined as interictal spikes (42), habitually followed by a slow wave lasting several hundreds of milliseconds (42; 23). For each patient, IEDs were manually annotated for the first 24 hours, according to standard guidelines (144; 145) using in-house developed software (MUSE) (135). IEDs were subsequently automatically detected for the full duration of the recording during (2-3 weeks) based on the following procedure (11):

1. The time-courses (trials) of the five deepest contacts were extracted (*−*500 ms to 1000 ms) at the original sample rate of 4 kHz according to manual annotations.
2. Trials were temporally aligned (c.f. (96)) based on the cross-correlation between each trial and the average of all trials. This procedure was performed iteratively by calculating a new average after all trials were shifted to their peak cross-correlation lag, until either the cross-correlation did not improve or a maximum of 10 iterations was reached. To calculate the cross-correlation lag over channels simultaneously, signals from the five electrode contacts were concatenated over time.
3. Once aligned, the trials were clustered in six k-mediods clusters using the MATLAB *kmedoids* function (The Mathworks Inc., Natick, Massachusetts).
4. If a cluster contained less than 2.5% of the total number of trials in that patient, these trials were automatically removed from the clustering procedure and the k-mediods clustering was repeated.
5. Each cluster was aligned to the LFP average of the cluster, similar to (2), then re-averaged, resulting in a patient-specific Channel *×* Time IED template.
6. Normalized cross-correlation was calculated between each template and the original data using the MATLAB *normxcorr2* function (The Mathworks Inc., Natick, Massachusetts).
7. Patient-specific correlation thresholds were visually determined based on the distribution of correlation values over time. IEDs were then detected when the peaks in the correlation values exceeded the threshold, using the MATLAB *findpeaks* function (The Mathworks Inc., Natick, Massachusetts)
8. IED detections that overlapped within a *±*50 ms period were assigned to the template with the highest normalized cross-correlation value.

The above automatic IED detection procedure effectively dealt with the within-subject variability of IED morphologies by using six clusters rather than a single template, as well as their spatial distribution by using normalized correlation over five contacts. Only a single patient-specific parameter (z-threshold) was needed. Importantly, this parameter was unbiased with regard to sleep stages. The procedure was straightforward and allowed the visual inspection of templates. To test the performance of the automatic detection procedure, the hit and false alarm rates were determined using a tolerance of *±*50 ms to allow for small differences in timing between manual and automatic detection.

### Spike sorting

After selecting electrodes that visually showed multi-unit activity (MUA) activity, clusters of action-potentials were automatically detected and clustered using Spyking Circus (146). Data was first high-pass filtered as 300 Hz using a 3^rd^ order Butterworth filter, then temporally whitened. Action potentials were detected at a user-defined 6 to 9 median absolute deviation, depending on the signal-to-noise level of the recording. Action potentials that occurred during artefacted periods were ignored. A combination of density-based clustering and template matching algorithms was used to cluster the detected action potentials (146). Clusters that did not show a clear morphology of action potentials or that were not stable across the recording were discarded manually. Finally, the clusters were labeled as putative single-unit activity (SUA) when they reflected well-isolated activity, based on the inter-spike interval (ISI), the percentage of violations of the refractory period (*RPV* = *ISI <* 1 ms), and action potential morphology. Otherwise, the clusters were labeled multiunit activity (MUA).

### Time-locked analyses

For each IED, the average LFP amplitudes of the spike and slow wave were extracted at latencies based on the patient average, i.e. independently from any analysis of sleep stages. LFPs were first baseline-corrected at *−*0.15 s to *−*0.05 s. For the sharp (positive) peaks, the latency was defined as periods between *−*0.15 s to 0.15 s at which the average LFP exceeded half of the maximum amplitude. For slow (negative) waves, the latency was defined as periods between *−*0.15 s to 0.05 s at which the average LFP was less than half of the minimum amplitude.

Spike times were epoched and time-locked to time periods identical to the LFP. For each IED, peri-stimulus time histogram (PSTH) spike rates were calculated with a bin size of 10 ms. To control for multiple comparisons and non-normal distributions of firing-rates, non-parametric cluster-based permutation tests (147) were used to determine time-periods where firing-rates changed significantly from baseline (*−*0.3 s to 0.1 s). A threshold of *p <* 0.01 (first-level t-test) was used to determine contiguous temporal clusters, after which a threshold of *p <* 0.05 (one-sided correction) determined whether the clusters could be explained by permutation (sum of t values, *n* = 10.000). Those clusters that significantly increased or decreased their firing rate time-locked to the IEDs were considered responsive. Clusters that showed spurious statistically significant results due to lack of data were manually changed to non-responsive. To increase the robustness of the PSTH spike rates during IEDs, templates were combined. Latencies for comparing the PSTH spike rates between sleep stages of the peak and slow wave were determined on the basis of the width at half-prominence of the average PSTH, i.e. independently from sleep stage. Pearson’s correlations between LFP and PSTH of each unit were calculated after downsampling LFP to the PSTH time axis (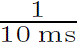 = 100 Hz).

### Sliding window analyses

Analyses of continuous data were based on segmentation of the data into non-overlapping windows of 10 seconds for both LFP and spike data. The number of IEDs that occurred during these time periods was determined. If a window overlapped *>* 50% with a sleep stage, it was labeled with the corresponding sleep stage. Windows that overlapped with an artefacted period were removed from further analyses. Oscillatory power was calculated for every window using a Hanning tapered fast Fourier transform (FFT). The 10 s windows provided a frequency resolution of 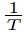 = 0.1 Hz. Slow wave activity (SWA) (148; 149) was defined as the mean power in the range of 0.1 Hz to 2.5 Hz, including the 0.5 Hz to 2 Hz range as suggested by the guidelines of the American Academy of Sleep (150). Delta activity was defined as the average power in the 2.5 Hz to 4 Hz range. For each window, the firing behavior of the unit was explored. Besides firing rate analyses, inter-spike variability was determined by means of CV2 (151; 152) and bursts were detected according to (108; 52), using a cut-off inter-spike intervals <5 ms. Interspike intervals and firing rates were corrected for bursts by removing bursting action potentials beyond the first action potential in a burst.

### Statistics

Statistical test were conducted with R (153; 154), version 4.1.2. Circular statistics were performed with the circular R package (155). Statistical plots were created with the ggplot R package (156). Models of the relationships between physiological measures were tested using mixed-effects linear models (157), using the lmer R package (158), with patient, template and night (LFP analyzes) or unit (spike analyzes) as random effects. Tukey’s post hoc tests were performed using the emmeans R package (159). The R scripts for all statistical analyzes and graphs can be found together with the analysis scripts here.

## Data availability

All scripts are made available here. Anonymous electrophysiology data can be made available on reasonable request for research purposes, but cannot be publicly shared due to legal constraints.

## Article and author information

According to CRediT standard:

**Stephen Whitmarsh**

Sorbonne Université, Institut du Cerveau – Paris Brain Institute – ICM, Inserm, CNRS, APHP, Pitié-Salpêtrière Hospital, Paris, France

**Contributions:** Conceptualization, Methodology, Software, Validation, Formal Analysis, Investigation, Original Draft Preparation, Review & Editing, Visualization

**Competing interests:** No competing interests declared

**Vi-Huong Nguyen-Michel**

AP-HP, Pitié-Salpêtrière Hospital, Epilepsy Unit and Reference Center for Rare Epilepsies, F-75013, Paris, France

**Contributions:** Methodology, Validation, Investigation, Review & Editing

**Competing interests:** No competing interests declared

**Katia Lehongre**

Sorbonne Université, Institut du Cerveau – Paris Brain Institute – ICM, Inserm, CNRS, APHP, Pitié-Salpêtrière Hospital, Paris, France

**Contributions:** Data-Curation, Software, Methodology, Review & Editing

**Competing:** interests: No competing interests declared

**Bertrand Mathon**

Sorbonne Université, Institut du Cerveau – Paris Brain Institute – ICM, Inserm, CNRS, APHP, Pitié-Salpêtrière Hospital, Paris, France

AP-HP, Department of Neurosurgery, Pitié-Salpêtrière Hospital and Sorbonne Université, Paris, France

**Contributions:** Validation, Methodology, Review & Editing

**Competing interests:** No competing interests declared

**Claude Adam**

AP-HP, Pitié-Salpêtrière Hospital, Epilepsy Unit and Reference Center for Rare Epilepsies, F-75013, Paris, France

**Contributions**: Validation, Investigation, Project Administration, Review & Editing

**Competing interests**: No competing interests declared

**Virginie Lambrecq**

Sorbonne Université, Institut du Cerveau – Paris Brain Institute – ICM, Inserm, CNRS, APHP, Pitié-Salpêtrière Hospital, Paris, France

AP-HP, Pitié-Salpêtrière Hospital, Epilepsy Unit and Reference Center for Rare Epilepsies, F-75013, Paris, France

**Contributions**: Validation, Investigation, Review & Editing

**Competing** interests: No competing interests declared

**Valerio Frazzini**

Sorbonne Université, Institut du Cerveau – Paris Brain Institute – ICM, Inserm, CNRS, APHP, Pitié-Salpêtrière Hospital, Paris, France

AP-HP, Pitié-Salpêtrière Hospital, Epilepsy Unit and Reference Center for Rare Epilepsies, F-75013, Paris, France

**Contributions**: Validation, Formal Analysis, Investigation, Supervision, Review & Editing

**Competing interests**: No competing interests declared

**Vincent Navarro**

Sorbonne Université, Institut du Cerveau – Paris Brain Institute – ICM, Inserm, CNRS, APHP, Pitié-Salpêtrière Hospital, Paris, France

AP-HP, Pitié-Salpêtrière Hospital, Epilepsy Unit and Reference Center for Rare Epilepsies, F-75013, Paris, France

**Contributions**: Conceptualization, Validation, Investigation, Resources, Review & Editing, Supervision, Project Administration, Funding Acquisition

**Competing interests**: No competing interests declared

## Acknowledgements

Part of this work was carried out on the CENIR-STIM platform of the Paris Brain Institute (ICM). We would like to thank the Reddit communities r/LaTeX and r/stat for their help on plotting and statistics.

## Supplementary Materials

**Fig. S1.**
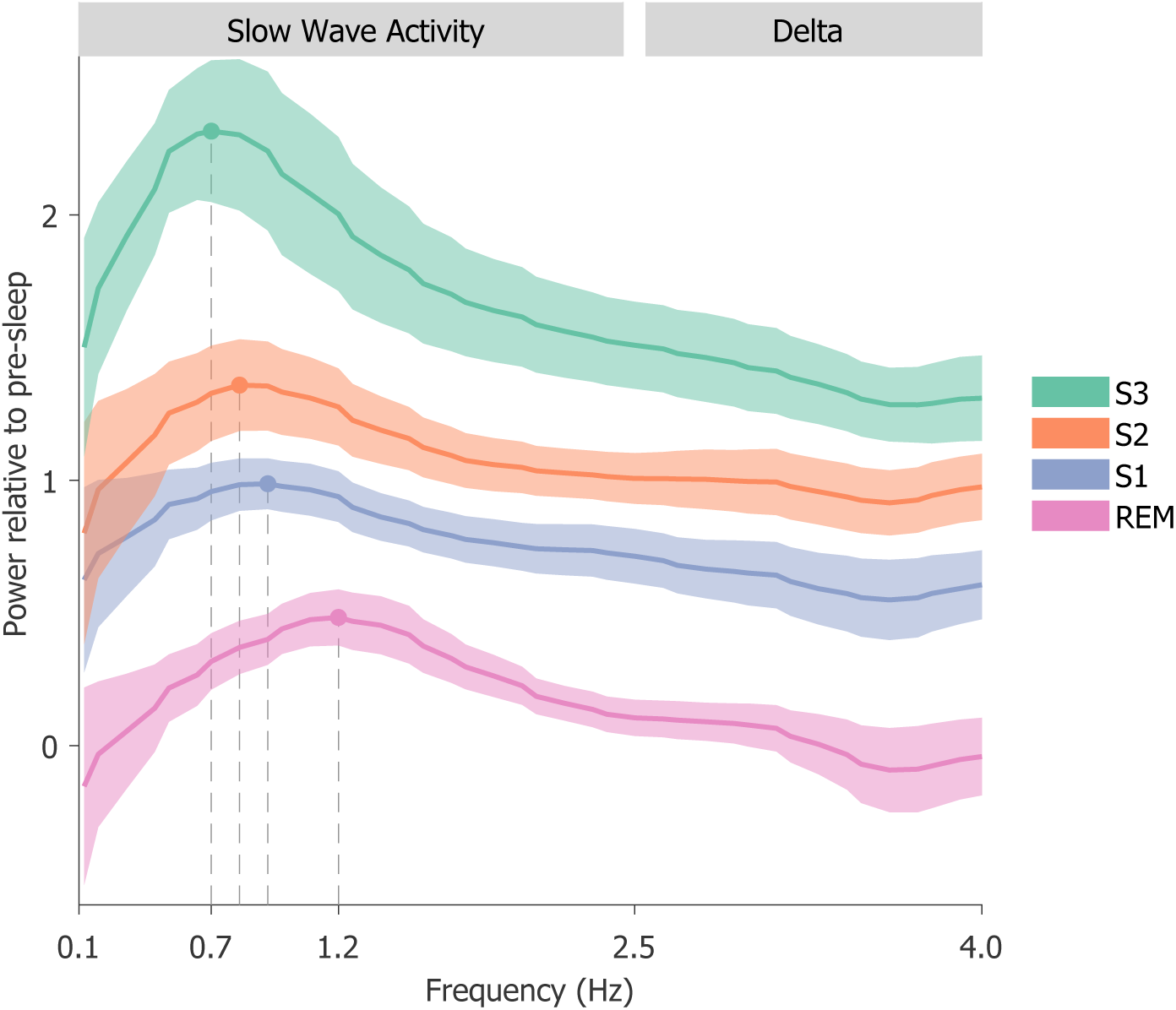
Power in the SWA and delta range, relative to the pre-sleep period, and separated per sleep stage. Lines show average power over patients per sleep stage, with shaded regions showing the standard error of the mean. Relative power showed a peak in the SWA frequency range (0.1 Hz to 2.5 Hz) which increased from 0.7 Hz in S3, to 0.8 Hz in S2, 0.9 Hz in S1, and 1.2 Hz in REM sleep.

**Fig. S2.**
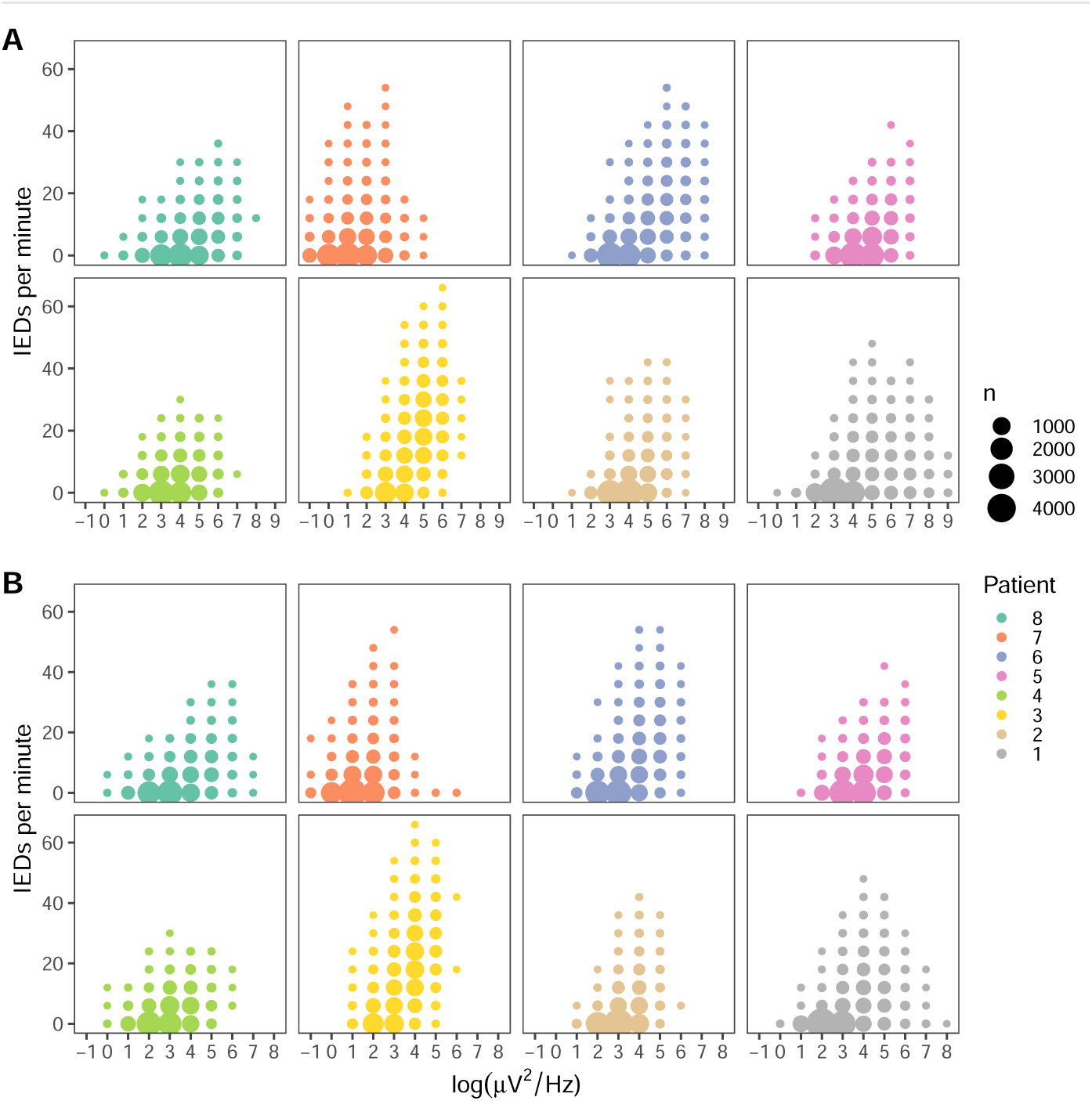
IED rates increase with increasing (**A**) slow wave activity (0.1 Hz to 2.5 Hz), and (**B**) delta (2.5 Hz to 4 Hz) power. For visualization, power was log-transformed, and both IED rates and power were binned. The size of the dots are proportional to the total number of observations in each bin. See Table S8 for statistical test.

**Table S1.** Clinical summary.

| Patient | Sex | Age | Onset | Type | MRI | PET | SPECT | Medication | Implantation | SOZ | Surgery |
| --- | --- | --- | --- | --- | --- | --- | --- | --- | --- | --- | --- |
| 1 | F | 18 | 10 | FSLA, FTBS | Normal | Mesial prefrontal cortex (R) | No localization | LTG, OXC | Fronto-Temporal (R) | Mesial prefrontal cortex (R) | Yes |
| 2 | M | 25 | 14 | FSLA | Normal | Temporo-polar and temporo-mesial (L) | Amygdala, anterior insula, and frontal opercular cortex (L) | ESL, TPM, PER | Temporo-Fronto-Insular (L) | Amygdalo-hippocampal region (L) | Amygdalo-hippampectomy (L) |
| 3 | F | 34 | 25 | FSWLA, FSLA, FTBS | Right SH (already resected at implantation) | Not available | Not available | VPA, CBZ, LTG | Bi-temporal | Bitemporal. Previously SH. Independent SOC in hippocampus (L) | No |
| 4 | F | 47 | 24 | FSWLA, FSLA, FTBS | Normal | Temporal pole (L) | Temporo basal and Lateral T1 (L) | ESL, LTG | Bi-temporal | Temporal (L) | No |
| 5 | F | 30 | 14 | FSLA, FTBS | Temporo posterior PNH (R), temporal SNH and PMG | No localization | Temporo-polar cortex (R), PNH and PMG | LTG, ZNG, LCS | Temporo-Insular (R) | Multifocal | No |
| 6 | F | 27 | 11 | FSWLA, FSLA, FTBS | Occipital porencephalic region (L) | Anterior hippocampus (L) | Temporal (L) and occipital | LCS, LTG | Temporo-Occipital (L) | Unknown | No |
| 7 | F | 21 | 14 | FSWLA, FSLA | Parietal gliotic lesion (L) | Parietal, temporal pole and temporo-lateral (L) | Temporal pole (L) | LCS, LTG | Parieto-Temporal (L) | Parietal (L) | No |
| 8 | F | 38 | 35 | FSLA | Normal | Temporomesial (R) | lateral temporal cortex (L) | LTC, VPA, CBZ | Temporal (R) | Amigdalo-hippocampal (R) | Amygdalo-hippampectomy (R) |

PET findings indicate regions of hypometabolism. SOZ are based on sEEG evaluation, but not yet validated post-surgery. AED: Antiepileptic drugs, CBZ: Carbamazepine, ESL: Eslicarbazepine, FBTCS: Focal to Bilateral Tonic-Clonic Seizures, FIAS: Focal Impaired Awareness Seizure, FSWLA: Focal seizures without loss of awareness, IEDs: Interictal Epileptiform discharges, LCS: Lacosamide, LTG: Lamotrigine, MRI: Indications from Magnetic Resonance Imaging, OXC: Oxcarbazepine, PER: Perampanel, PMG: Polymicrogyria, PNH: Periventricular Nodular Heterotopia, SNH: Subcortical Nodular Heterotopia, SOZ: Seizure Onset Zone, TPM: Topiramate VPA: Valproic Acid, ZNG: Zonisamide.

**Table S2.** Anatomical locations of macro and micro electrodes.

| Patient | Side | Region | Subregion | SOZ |
| --- | --- | --- | --- | --- |
| **Macro contacts** |  |  |  |  |
| 1 | R | Hippocampal formation | Head of the Hippocampus | No |
| 2 | L | Hippocampal formation | Head of the Hippocampus | Yes |
|  | L | Hippocampal formation | Hippocampus/CA1 | Yes |
| 3 | L | Hippocampal formation | Head of the Hippocampus | Yes |
|  | L | Hippocampal formation | Entorhinal cortex | No |
| 4 | L | Hippocampal formation | Hippocampus | No |
|  | L | Hippocampal formation | Hippocampus/CA3 | No |
| 5 | R | Hippocampal formation | Subiculum | Yes |
|  | R | Nodule | Nodule | Yes |
| 6 | L | Hippocampal formation | Entorhinal cortex | No |
|  | L | Hippocampal formation | Parahippocampus | No |
| 7 | R | Hippocampal formation | Entorhinal cortex | Yes |
|  | R | Hippocampal formation | Hippocampus/CA1 | Yes |
| 8 | R | Hippocampal formation | Hippocampus/CA1 | Yes |
| **Micro electrodes** |  |  |  |  |
| 1 | R | Hippocampal formation | Entorhinal cortex | Yes |
| 2 | L | Hippocampal formation | Entorhinal cortex | Yes |
| 3 | R | Temporo-basal cortex | Parahippocampus | No |
|  | L | Temporo-basal cortex | Parahippocampus | Yes |
| 4 | L | Temporo-anterior lobe | Amygdala | No |
|  | L | Mesial temporal cortex | CA3-CA4 | No |
| 5 | R | Periventricular nodule | Nodule | Yes |
|  | R | Periventricular nodule | Nodule | Yes |
|  | R | Hippocampal formation | Subiculum | No |
| 6 | L | Hippocampal formation | Parahippocampal gyrus | No |
| 7 | R | Temporo-anterior lobe | Amygdala | Yes |
| 8 | R | Hippocampal formation | CA1/WM | Yes |
|  | R | Hippocampal formation | CA1 | Yes |

**Table S3.** Automatic IED detection performance.

| Patient | Visual | Automatic detection |  |  |  |  |
| --- | --- | --- | --- | --- | --- | --- |
|  | 24 hrs. | 24 hrs. | Hit (%) | FA (%) | Total | Total hrs. |
| 1 | 2106 | 3500 | 99.8 | 5.4 | 63345 | 453.5 |
| 2 | 3330 | 4618 | 99.3 | 6.9 | 9224 | 358.3 |
| 3 | 12912 | 12811 | 98.8 | 1.2 | 113832 | 305.6 |
| 4 | 3669 | 3664 | 97.9 | 11.0 | 43757 | 308.8 |
| 5 | 7086 | 4630 | 96.5 | 8.6 | 32782 | 237.2 |
| 6 | 3077 | 3054 | 98.9 | 3.8 | 29810 | 233.6 |
| 7 | 10439 | 3985 | 98.4 | 0.8 | 55465 | 305.6 |
| 8 | 2714 | 1965 | 94.1 | 14.3 | 53951 | 307.7 |
| *Mean* | 5667 | 4778 | 98.0 | 6.5 | 50271 | 313.8 |
| *Std.* | 4050 | 3359 | 1.9 | 4.7 | 30945 | 69.7 |

**Table S4:**
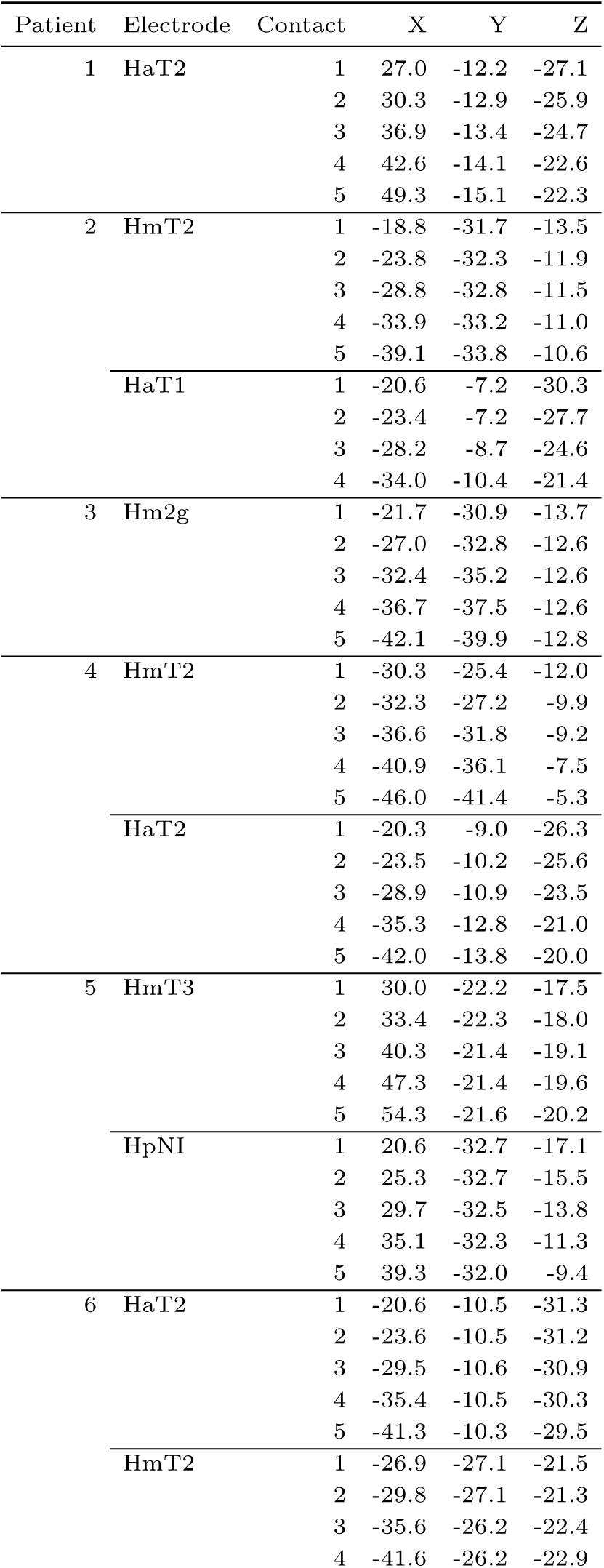
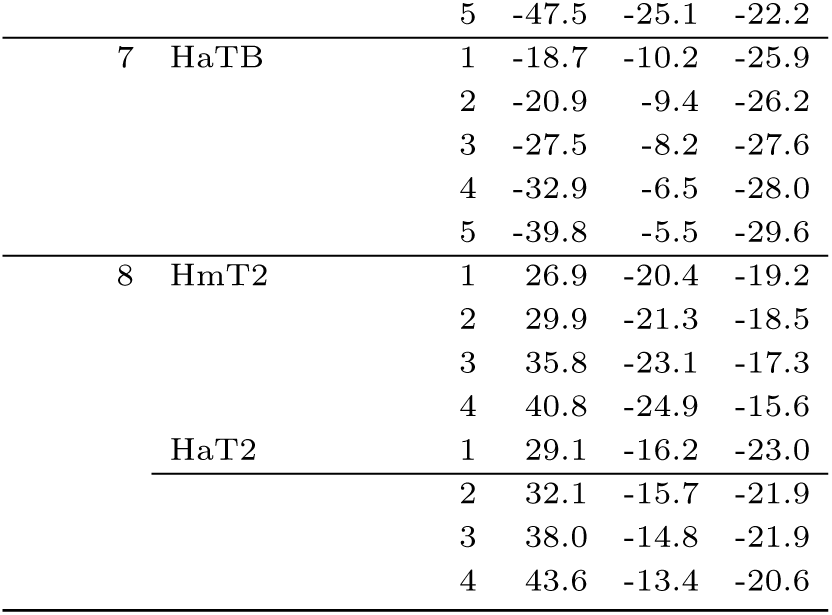
Anatomical locations of micro electrodes.

**Table S5.** Circular statistics of circadian epileptic activity.

| Patient | Interictal activity |  |  | Seizures |  |  |
| --- | --- | --- | --- | --- | --- | --- |
|  | Time | Rayleigh | *p* | Time | Rayleigh | *p* |
| 1 | 01:56 | 0.61 | <.0001*** | 17:43 | 0.76 | <.0001*** |
| 2 | 00:46 | 0.29 | <.0001*** | 08:56 | 0.36 | 0.55 |
| 3 | 01:37 | 0.50 | <.0001*** | 10:31 | 0.48 | <.01* |
| 4 | 02:23 | 0.26 | <.0001*** | 18:26 | 0.94 | <.001** |
| 5 | 01:48 | 0.28 | <.0001*** | 20:21 | 0.13 | 0.3 |
| 6 | 01:42 | 0.56 | <.0001*** | 03:33 | 0.57 | 0.14 |
| 7 | 17:25 | 0.08 | <.0001*** | 01:59 | 0.81 | <.0001*** |
| 8 | 03:38 | 0.41 | <.0001*** | 11:44 | 0.70 | 0.14 |

. = $p < 0.05$ , \* = $p < 0.01$ , \*\* = $p < 0.001$ , \*\*\* = $p < 0.0001$

**Table S6.** Time spend in sleep stages.

| Patient | Night | TST | % TST |  |  |  | WASO |
| --- | --- | --- | --- | --- | --- | --- | --- |
|  |  |  | S3 | S2 | S1 | REM |  |
| 1 | 1 | 11.9 | 13 | 44 | 21 | 22 | 1.6 |
|  | 2 | 8.7 | 14 | 42 | 18 | 25 | 0.9 |
|  | 3 | 7.5 | 22 | 42 | 17 | 19 | 3 |
| 2 | 1 | 8.5 | 26 | 52 | 7 | 15 | 2.3 |
|  | 2 | 7.6 | 33 | 39 | 8 | 19 | 0.4 |
|  | 3 | 8.1 | 24 | 52 | 8 | 16 | 0.4 |
| 3 | 1 | 9.2 | 13 | 45 | 26 | 16 | 1.7 |
|  | 2 | 7.2 | 27 | 33 | 24 | 16 | 1.7 |
|  | 3 | 7.2 | 28 | 33 | 27 | 12 | 1.5 |
| 4 | 1 | 7.5 | 23 | 38 | 26 | 13 | 1.7 |
|  | 2 | 6.7 | 16 | 41 | 26 | 17 | 1.3 |
|  | 3 | 6.5 | 22 | 27 | 19 | 31 | 0.9 |
| 5 | 1 | 9.6 | 9 | 67 | 9 | 15 | 1.3 |
|  | 2 | 7.2 | 26 | 42 | 10 | 22 | 0.4 |
|  | 3 | 7.2 | 15 | 46 | 9 | 30 | 0.2 |
| 6 | 1 | 7.5 | 15 | 43 | 25 | 17 | 1.2 |
|  | 2 | 8.2 | 21 | 35 | 15 | 29 | 0.9 |
|  | 3 | 8.9 | 18 | 38 | 18 | 26 | 1.3 |
| 7 | 1 | 6.9 | 31 | 41 | 14 | 14 | 0.4 |
|  | 2 | 9.6 | 28 | 46 | 7 | 19 | 0.5 |
|  | 3 | 10.3 | 35 | 37 | 8 | 20 | 0.9 |
| 8 | 1 | 7 | 16 | 54 | 10 | 19 | 0.4 |
|  | 2 | 7.1 | 10 | 52 | 16 | 22 | 0.6 |
|  | 3 | 6.8 | 15 | 50 | 20 | 15 | 3.7 |
| *Mean* |  | 8.0 | 20.8 | 43.3 | 16.2 | 19.5 | 1.2 |

TST = Total sleep time (hrs.), WASO = Wake after sleep onset (hrs.)

**Table S7.** Effect of sleep stage on IEDs rate.

| Predictor | Coef $\beta$ | SE($\beta$) | df | z | p |
| --- | --- | --- | --- | --- | --- |
| **Sleep stages** |  |  |  |  |  |
| *Intercept* | 0.26 | 0.23 | 7.03 | 1.12 | 0.3 |
| S3 | 1.13 | 0.01 | 93176.95 | 79.21 | <.0001*** |
| S2 | 0.90 | 0.01 | 93176.88 | 70.85 | <.0001*** |
| S1 | 0.54 | 0.02 | 93176.31 | 35.04 | <.0001*** |
| WASO | 0.10 | 0.02 | 93176.64 | 6.08 | <.0001*** |
| REM | 0.00 | 0.01 | 93176.50 | -0.31 | 0.76 |
| Post | 0.01 | 0.02 | 93176.92 | 0.43 | 0.67 |
| **Post-hoc comparisons** |  |  |  |  |  |
| Pre - S3 | -1.13 | 0.01 |  | -79.21 | <.0001*** |
| Pre - S2 | -0.90 | 0.01 |  | -70.85 | <.0001*** |
| Pre - S1 | -0.54 | 0.02 |  | -35.04 | <.0001*** |
| Pre - WASO | -0.10 | 0.02 |  | -6.08 | <.0001*** |
| Pre - REM | 0.00 | 0.01 |  | 0.31 | 1 |
| Pre - Post | -0.01 | 0.02 |  | -0.43 | 1 |
| S3 - S2 | 0.23 | 0.01 |  | 21.19 | <.0001*** |
| S3 - S1 | 0.59 | 0.01 |  | 41.55 | <.0001*** |
| S3 - WASO | 1.03 | 0.02 |  | 63.83 | <.0001*** |
| S3 - REM | 1.14 | 0.01 |  | 85.89 | <.0001*** |
| S3 - Post | 1.12 | 0.01 |  | 80.70 | <.0001*** |
| S2 - S1 | 0.36 | 0.01 |  | 28.35 | <.0001*** |

**Table S8.** Effect of Slow Wave activity (0.1-2.5Hz) and Delta power (2.5-4Hz) on rate of IEDs.

| | Coef $\beta$ | SE($\beta$) | df | z | p |
| --- | --- | --- | --- | --- | --- |
| *Intercept* | 0.412 | 0.217 | 7 | 1.90 | 0.1 |
| Slow Wave | 0.000 | 0.000 | 93182 | 14.02 | <.0001*** |
| Delta | 0.010 | 0.000 | 93184 | 101.95 | <.0001*** |

. = $p < 0.05$ , \* = $p < 0.01$ , \*\* = $p < 0.001$ , \*\*\* = $p < 0.0001$

**Table S9.** Effect of sleep stage on Slow Wave (0.1-2.5Hz) power.

| Predictor | Coef $\beta$ | SE($\beta$) | df | z | p |
| --- | --- | --- | --- | --- | --- |
| **Coefficients** |  |  |  |  |  |
| *Intercept* | 29.03 | 25.16 | 7.11 | 1.15 | 0.29 |
| S3 | 264.51 | 3.00 | 93179.16 | 88.30 | <.0001*** |
| S2 | 93.86 | 2.65 | 93178.96 | 35.37 | <.0001*** |
| S1 | 48.34 | 3.22 | 93177.11 | 15.00 | <.0001*** |
| WASO | 18.02 | 3.59 | 93178.22 | 5.02 | <.0001*** |
| REM | 14.50 | 3.05 | 93177.76 | 4.76 | <.0001*** |
| Post | 5.69 | 3.20 | 93179.09 | 1.78 | 0.08 |
| **Post-hoc comparisons** |  |  |  |  |  |
| Pre - S3 | -264.51 | 3.00 |  | -88.30 | <.0001*** |
| Pre - S2 | -93.86 | 2.65 |  | -35.37 | <.0001*** |
| Pre - S1 | -48.34 | 3.22 |  | -15.00 | <.0001*** |
| Pre - WASO | -18.02 | 3.59 |  | -5.02 | <.0001*** |
| Pre - REM | -14.50 | 3.05 |  | -4.76 | <.0001*** |
| Pre - Post | -5.69 | 3.20 |  | -1.78 | 0.56 |
| S3 - S2 | 170.65 | 2.33 |  | 73.38 | <.0001*** |
| S3 - S1 | 216.17 | 2.99 |  | 72.20 | <.0001*** |
| S3 - WASO | 246.49 | 3.38 |  | 73.03 | <.0001*** |
| S3 - REM | 250.02 | 2.77 |  | 90.14 | <.0001*** |
| S3 - Post | 258.82 | 2.92 |  | 88.53 | <.0001*** |
| S2 - S1 | 45.52 | 2.65 |  | 17.17 | <.0001*** |
| S2 - WASO | 75.84 | 3.07 |  | 24.72 | <.0001*** |
| S2 - REM | 79.36 | 2.40 |  | 33.11 | <.0001*** |
| S2 - Post | 88.17 | 2.58 |  | 34.22 | <.0001*** |
| S1 - WASO | 30.32 | 3.56 |  | 8.51 | <.0001*** |
| S1 - REM | 33.85 | 3.04 |  | 11.12 | <.0001*** |
| S1 - Post | 42.65 | 3.19 |  | 13.39 | <.0001*** |
| WASO - REM | 3.52 | 3.42 |  | 1.03 | 0.95 |
| WASO - Post | 12.33 | 3.54 |  | 3.48 | <.01* |
| REM - Post | 8.81 | 2.99 |  | 2.95 | <.05 |

. = $p < 0.05$ , \* = $p < 0.01$ , \*\* = $p < 0.001$ , \*\*\* = $p < 0.0001$

**Table S10.** Effect of sleep stage on Delta (2.5-4 Hz) power.

| Predictor | Coef $\beta$ | SE($\beta$) | df | z | p |
| --- | --- | --- | --- | --- | --- |
| **Coefficients** |  |  |  |  |  |
| *Intercept* | 15.24 | 6.58 | 7.05 | 2.31 | 0.05 |
| S3 | 54.64 | 0.55 | 93177.66 | 99.54 | <.0001*** |
| S2 | 29.96 | 0.49 | 93177.55 | 61.61 | <.0001*** |
| S1 | 16.54 | 0.59 | 93176.55 | 28.02 | <.0001*** |
| WASO | 3.96 | 0.66 | 93177.13 | 6.02 | <.0001*** |
| REM | 2.25 | 0.56 | 93176.89 | 4.03 | <.0001*** |
| Post | 0.22 | 0.59 | 93177.62 | 0.38 | 0.71 |
| **Post-hoc comparisons** |  |  |  |  |  |
| Pre - S3 | -54.64 | 0.55 |  | -99.54 | <.0001*** |
| Pre - S2 | -29.96 | 0.49 |  | -61.61 | <.0001*** |
| Pre - S1 | -16.54 | 0.59 |  | -28.02 | <.0001*** |
| Pre - WASO | -3.96 | 0.66 |  | -6.02 | <.0001*** |
| Pre - REM | -2.25 | 0.56 |  | -4.03 | <.01* |
| Pre - Post | -0.22 | 0.59 |  | -0.38 | 1 |
| S3 - S2 | 24.68 | 0.43 |  | 57.92 | <.0001*** |
| S3 - S1 | 38.09 | 0.55 |  | 69.44 | <.0001*** |
| S3 - WASO | 50.68 | 0.62 |  | 81.95 | <.0001*** |
| S3 - REM | 52.39 | 0.51 |  | 103.08 | <.0001*** |
| S3 - Post | 54.42 | 0.54 |  | 101.58 | <.0001*** |
| S2 - S1 | 13.42 | 0.49 |  | 27.63 | <.0001*** |
| S2 - WASO | 26.00 | 0.56 |  | 46.25 | <.0001*** |
| S2 - REM | 27.71 | 0.44 |  | 63.09 | <.0001*** |
| S2 - Post | 29.74 | 0.47 |  | 63.00 | <.0001*** |
| S1 - WASO | 12.59 | 0.65 |  | 19.28 | <.0001*** |
| S1 - REM | 14.30 | 0.56 |  | 25.62 | <.0001*** |
| S1 - Post | 16.32 | 0.58 |  | 27.96 | <.0001*** |
| WASO - REM | 1.71 | 0.63 |  | 2.73 | 0.09 |
| WASO - Post | 3.74 | 0.65 |  | 5.76 | <.0001*** |
| REM - Post | 2.03 | 0.55 |  | 3.70 | <.01* |

. = $p < 0.05$ , \* = $p < 0.01$ , \*\* = $p < 0.001$ , \*\*\* = $p < 0.0001$

**Table S11.**
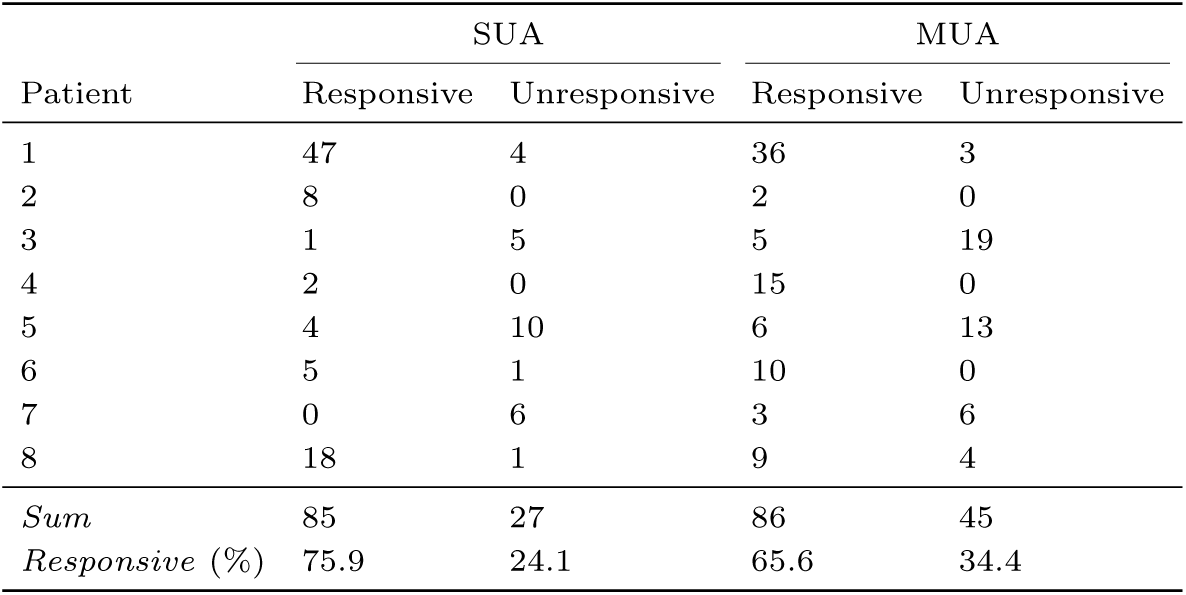
Number of responsive or unresponsive putatively isolated single units (SUA) and multiunits (MUA)

| Patient | SUA |  | MUA |  |
| --- | --- | --- | --- | --- |
|  | Responsive | Unresponsive | Responsive | Unresponsive |
| 1 | 47 | 4 | 36 | 3 |
| 2 | 8 | 0 | 2 | 0 |
| 3 | 1 | 5 | 5 | 19 |
| 4 | 2 | 0 | 15 | 0 |
| 5 | 4 | 10 | 6 | 13 |
| 6 | 5 | 1 | 10 | 0 |
| 7 | 0 | 6 | 3 | 6 |
| 8 | 18 | 1 | 9 | 4 |
| *Sum* | 85 | 27 | 86 | 45 |
| *Responsive (%)* | 75.9 | 24.1 | 65.6 | 34.4 |

**Table S12.** Effect of sleep stage on ERP peak amplitude.

|  | Spike amplitude |  |  |  |  | Slow wave amplitude |  |  |  |  | Difference amplitude |  |  |  |  |
| --- | --- | --- | --- | --- | --- | --- | --- | --- | --- | --- | --- | --- | --- | --- | --- |
| | Coef $\beta$ | SE($\beta$) | df | z | p | Coef $\beta$ | SE($\beta$) | df | z | p | Coef $\beta$ | SE($\beta$) | df | z | p |
| **Sleep stages** |  |  |  |  |  |  |  |  |  |  |  |  |  |  |  |
| *Intercept* | 425.78 | 108.40 | 7 | 3.93 | <.01* | -260.53 | 97.66 | 7 | -2.67 | <.05 | -260.53 | 97.66 | 7 | -2.67 | <.05 |
| S3 | 84.05 | 2.98 | 78969 | 28.19 | <.0001*** | -29.52 | 4.24 | 78968 | -6.96 | <.0001*** | -29.52 | 4.24 | 78968 | -6.96 | <.0001*** |
| S2 | 63.59 | 2.83 | 78967 | 22.46 | <.0001*** | -35.60 | 4.03 | 78967 | -8.85 | <.0001*** | -35.60 | 4.03 | 78967 | -8.85 | <.0001*** |
| S1 | 49.95 | 3.18 | 78966 | 15.70 | <.0001*** | -29.32 | 4.53 | 78966 | -6.48 | <.0001*** | -29.32 | 4.53 | 78966 | -6.48 | <.0001*** |
| WASO | 19.01 | 3.88 | 78965 | 4.91 | <.0001*** | -10.38 | 5.51 | 78966 | -1.88 | 0.06 | -10.38 | 5.51 | 78966 | -1.88 | 0.06 |
| Post | 7.56 | 4.35 | 78970 | 1.74 | 0.08 | 5.74 | 6.18 | 78969 | 0.93 | 0.35 | 5.74 | 6.18 | 78969 | 0.93 | 0.35 |
| REM | -61.68 | 4.33 | 78977 | -14.24 | <.0001*** | 87.42 | 6.16 | 78973 | 14.19 | <.0001*** | 87.42 | 6.16 | 78973 | 14.19 | <.0001*** |
| **Post-hoc comparisons** |  |  |  |  |  |  |  |  |  |  |  |  |  |  |  |
| Pre - S3 | -84.05 | 2.98 |  | -28.19 | <.0001*** | 29.52 | 4.24 |  | 6.96 | <.0001*** | -113.58 | 5.45 |  | -20.82 | <.0001*** |
| Pre - S2 | -63.59 | 2.83 |  | -22.46 | <.0001*** | 35.60 | 4.03 |  | 8.85 | <.0001*** | -99.18 | 5.18 |  | -19.15 | <.0001*** |
| Pre - S1 | -49.95 | 3.18 |  | -15.70 | <.0001*** | 29.32 | 4.53 |  | 6.48 | <.0001*** | -79.26 | 5.82 |  | -13.61 | <.0001*** |
| Pre - WASO | -19.01 | 3.88 |  | -4.91 | <.0001*** | 10.38 | 5.51 |  | 1.88 | 0.49 | -29.38 | 7.09 |  | -4.14 | <.001** |
| Pre - REM | -7.56 | 4.35 |  | -1.74 | 0.59 | -5.74 | 6.18 |  | -0.93 | 0.97 | -1.78 | 7.95 |  | -0.22 | 1 |
| Pre - Post | 61.68 | 4.33 |  | 14.24 | <.0001*** | -87.42 | 6.16 |  | -14.19 | <.0001*** | 149.09 | 7.92 |  | 18.81 | <.0001*** |
| S3 - S2 | 20.46 | 1.76 |  | 11.63 | <.0001*** | 6.08 | 2.50 |  | 2.43 | 0.19 | 14.40 | 3.22 |  | 4.47 | <.001** |
| S3 - S1 | 34.10 | 2.38 |  | 14.32 | <.0001*** | -0.21 | 3.39 |  | -0.06 | 1 | 34.32 | 4.36 |  | 7.88 | <.0001*** |
| S3 - WASO | 65.04 | 3.26 |  | 19.95 | <.0001*** | -19.14 | 4.63 |  | -4.13 | <.001** | 84.19 | 5.96 |  | 14.12 | <.0001*** |
| S3 - REM | 76.49 | 3.76 |  | 20.32 | <.0001*** | -35.26 | 5.35 |  | -6.59 | <.0001*** | 111.80 | 6.89 |  | 16.23 | <.0001*** |
| S3 - Post | 145.73 | 3.80 |  | 38.33 | <.0001*** | -116.94 | 5.41 |  | -21.63 | <.0001*** | 262.67 | 6.95 |  | 37.77 | <.0001*** |
| S2 - S1 | 13.64 | 2.17 |  | 6.30 | <.0001*** | -6.29 | 3.08 |  | -2.04 | 0.39 | 19.92 | 3.96 |  | 5.03 | <.0001*** |
| S2 - WASO | 44.58 | 3.12 |  | 14.29 | <.0001*** | -25.22 | 4.44 |  | -5.68 | <.0001*** | 69.80 | 5.71 |  | 12.23 | <.0001*** |
| S2 - REM | 56.03 | 3.66 |  | 15.32 | <.0001*** | -41.34 | 5.20 |  | -7.95 | <.0001*** | 97.40 | 6.69 |  | 14.55 | <.0001*** |
| S2 - Post | 125.27 | 3.69 |  | 33.91 | <.0001*** | -123.02 | 5.25 |  | -23.42 | <.0001*** | 248.27 | 6.76 |  | 36.73 | <.0001*** |
| S1 - WASO | 30.94 | 3.44 |  | 9.00 | <.0001*** | -18.93 | 4.89 |  | -3.87 | <.01* | 49.87 | 6.29 |  | 7.93 | <.0001*** |
| S1 - REM | 42.39 | 3.95 |  | 10.72 | <.0001*** | -35.05 | 5.62 |  | -6.23 | <.0001*** | 77.48 | 7.23 |  | 10.71 | <.0001*** |
| S1 - Post | 111.63 | 4.01 |  | 27.85 | <.0001*** | -116.73 | 5.70 |  | -20.48 | <.0001*** | 228.35 | 7.33 |  | 31.15 | <.0001*** |
| WASO - REM | 11.45 | 4.55 |  | 2.52 | 0.15 | -16.12 | 6.46 |  | -2.49 | 0.16 | 27.61 | 8.32 |  | 3.32 | <.05 |
| WASO - Post | 80.69 | 4.59 |  | 17.56 | <.0001*** | -97.80 | 6.53 |  | -14.97 | <.0001*** | 178.48 | 8.41 |  | 21.23 | <.0001*** |
| REM - Post | 69.23 | 4.95 |  | 13.98 | <.0001*** | -81.68 | 7.04 |  | -11.59 | <.0001*** | 150.87 | 9.06 |  | 16.65 | <.0001*** |

. = $p < 0.05$ , \* = $p < 0.01$ , \*\* = $p < 0.001$ , \*\*\* = $p < 0.0001$

**Table S13.** Effect of sleep stage on firing rates during IED.

|  | Spike |  |  |  |  | Wave |  |  |  |  | Ratio |  |  |  |  |
| --- | --- | --- | --- | --- | --- | --- | --- | --- | --- | --- | --- | --- | --- | --- | --- |
| | Coef $\beta$ | SE($\beta$) | df | z | p | Coef $\beta$ | SE($\beta$) | df | z | p | Coef $\beta$ | SE($\beta$) | df | z | p |
| **Sleep stages** |  |  |  |  |  |  |  |  |  |  |  |  |  |  |  |
| Intercept | 6.20 | 1.49 | 6 | 4.17 | <.01* | 11.05 | 3.08 | 7 | 3.59 | <.01* | 4.83 | 2.20 | 7 | 2.19 | 0.07 |
| S3 | -2.21 | 0.38 | 1237 | -5.88 | <.0001*** | 2.66 | 0.65 | 1237 | 4.12 | <.0001*** | 4.87 | 0.58 | 1237 | 8.45 | <.0001*** |
| S2 | -2.01 | 0.38 | 1237 | -5.33 | <.0001*** | 3.20 | 0.65 | 1237 | 4.95 | <.0001*** | 5.20 | 0.58 | 1237 | 9.02 | <.0001*** |
| S1 | -1.68 | 0.38 | 1237 | -4.45 | <.0001*** | 3.30 | 0.65 | 1237 | 5.11 | <.0001*** | 4.98 | 0.58 | 1237 | 8.63 | <.0001*** |
| WASO | -2.03 | 0.38 | 1237 | -5.38 | <.0001*** | 1.34 | 0.65 | 1237 | 2.08 | <.05* | 3.37 | 0.58 | 1237 | 5.84 | <.0001*** |
| Post | -0.68 | 0.38 | 1237 | -1.82 | 0.07 | -0.19 | 0.65 | 1237 | -0.29 | 0.77 | 0.50 | 0.58 | 1237 | 0.86 | 0.39 |
| REM | -0.20 | 0.38 | 1237 | -0.51 | 0.61 | -1.28 | 0.65 | 1237 | -1.97 | <.05* | -1.09 | 0.58 | 1237 | -1.87 | 0.06 |
| **Post-hoc comparisons** |  |  |  |  |  |  |  |  |  |  |  |  |  |  |  |
| Pre - S3 | 2.21 | 0.38 |  | 5.88 | <.0001*** | -2.66 | 0.65 |  | -4.12 | <.001** | -4.87 | 0.58 |  | -8.45 | <.0001*** |
| Pre - S2 | 2.01 | 0.38 |  | 5.33 | <.0001*** | -3.20 | 0.65 |  | -4.95 | <.0001*** | -5.20 | 0.58 |  | -9.02 | <.0001*** |
| Pre - S1 | 1.68 | 0.38 |  | 4.45 | <.001** | -3.30 | 0.65 |  | -5.11 | <.0001*** | -4.98 | 0.58 |  | -8.63 | <.0001*** |
| Pre - WASO | 2.03 | 0.38 |  | 5.38 | <.0001*** | -1.34 | 0.65 |  | -2.08 | 0.37 | -3.37 | 0.58 |  | -5.84 | <.0001*** |
| Pre - REM | 0.68 | 0.38 |  | 1.82 | 0.54 | 0.19 | 0.65 |  | 0.29 | 1 | -0.50 | 0.58 |  | -0.86 | 0.98 |
| Pre - Post | 0.20 | 0.38 |  | 0.51 | 1 | 1.28 | 0.65 |  | 1.97 | 0.44 | 1.09 | 0.58 |  | 1.87 | 0.5 |
| S3 - S2 | -0.21 | 0.38 |  | -0.55 | 1 | -0.54 | 0.65 |  | -0.83 | 0.98 | -0.33 | 0.58 |  | -0.57 | 1 |
| S3 - S1 | -0.54 | 0.38 |  | -1.42 | 0.79 | -0.64 | 0.65 |  | -0.99 | 0.96 | -0.11 | 0.58 |  | -0.18 | 1 |
| S3 - WASO | -0.19 | 0.38 |  | -0.50 | 1 | 1.32 | 0.65 |  | 2.05 | 0.39 | 1.51 | 0.58 |  | 2.61 | 0.12 |
| S3 - REM | -1.53 | 0.38 |  | -4.06 | <.01* | 2.85 | 0.65 |  | 4.41 | <.001** | 4.38 | 0.58 |  | 7.59 | <.0001*** |
| S3 - Post | -2.02 | 0.38 |  | -5.32 | <.0001*** | 3.94 | 0.65 |  | 6.06 | <.0001*** | 5.96 | 0.58 |  | 10.26 | <.0001*** |
| S2 - S1 | -0.33 | 0.38 |  | -0.87 | 0.98 | -0.10 | 0.65 |  | -0.16 | 1 | 0.22 | 0.58 |  | 0.39 | 1 |
| S2 - WASO | 0.02 | 0.38 |  | 0.05 | 1 | 1.86 | 0.65 |  | 2.88 | 0.06 | 1.84 | 0.58 |  | 3.18 | <.05* |
| S2 - REM | -1.32 | 0.38 |  | -3.51 | <.01* | 3.38 | 0.65 |  | 5.24 | <.0001*** | 4.70 | 0.58 |  | 8.16 | <.0001*** |
| S2 - Post | -1.81 | 0.38 |  | -4.77 | <.0001*** | 4.48 | 0.65 |  | 6.89 | <.0001*** | 6.29 | 0.58 |  | 10.83 | <.0001*** |
| S1 - WASO | 0.35 | 0.38 |  | 0.93 | 0.97 | 1.96 | 0.65 |  | 3.04 | <.05* | 1.61 | 0.58 |  | 2.80 | 0.08 |
| S1 - REM | -0.99 | 0.38 |  | -2.64 | 0.12 | 3.49 | 0.65 |  | 5.40 | <.0001*** | 4.48 | 0.58 |  | 7.77 | <.0001*** |
| S1 - Post | -1.48 | 0.38 |  | -3.91 | <.01* | 4.58 | 0.65 |  | 7.05 | <.0001*** | 6.07 | 0.58 |  | 10.44 | <.0001*** |
| WASO - REM | -1.34 | 0.38 |  | -3.56 | <.01* | 1.53 | 0.65 |  | 2.36 | 0.21 | 2.87 | 0.58 |  | 4.97 | <.0001*** |
| WASO - Post | -1.83 | 0.38 |  | -4.83 | <.0001*** | 2.62 | 0.65 |  | 4.03 | <.01* | 4.45 | 0.58 |  | 7.67 | <.0001*** |
| REM - Post | -0.49 | 0.38 |  | -1.29 | 0.86 | 1.09 | 0.65 |  | 1.68 | 0.63 | 1.58 | 0.58 |  | 2.73 | 0.09 |

**Table S14.**
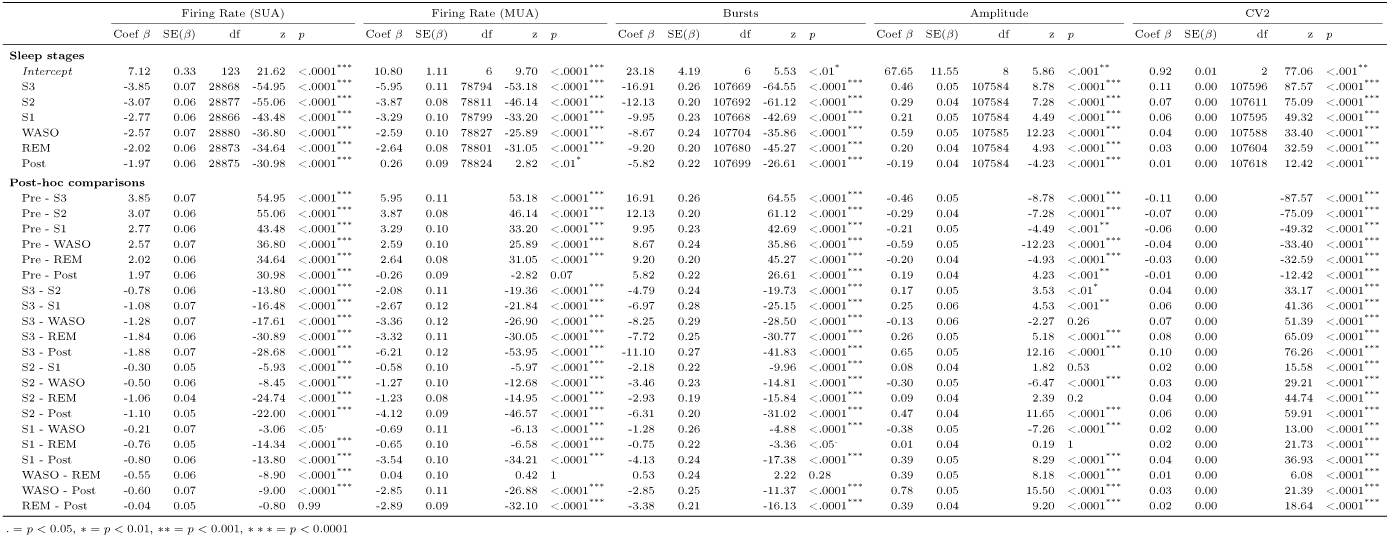
Effect of sleep stage on spontanious neuronal behaviour.

**Fig. S3.**
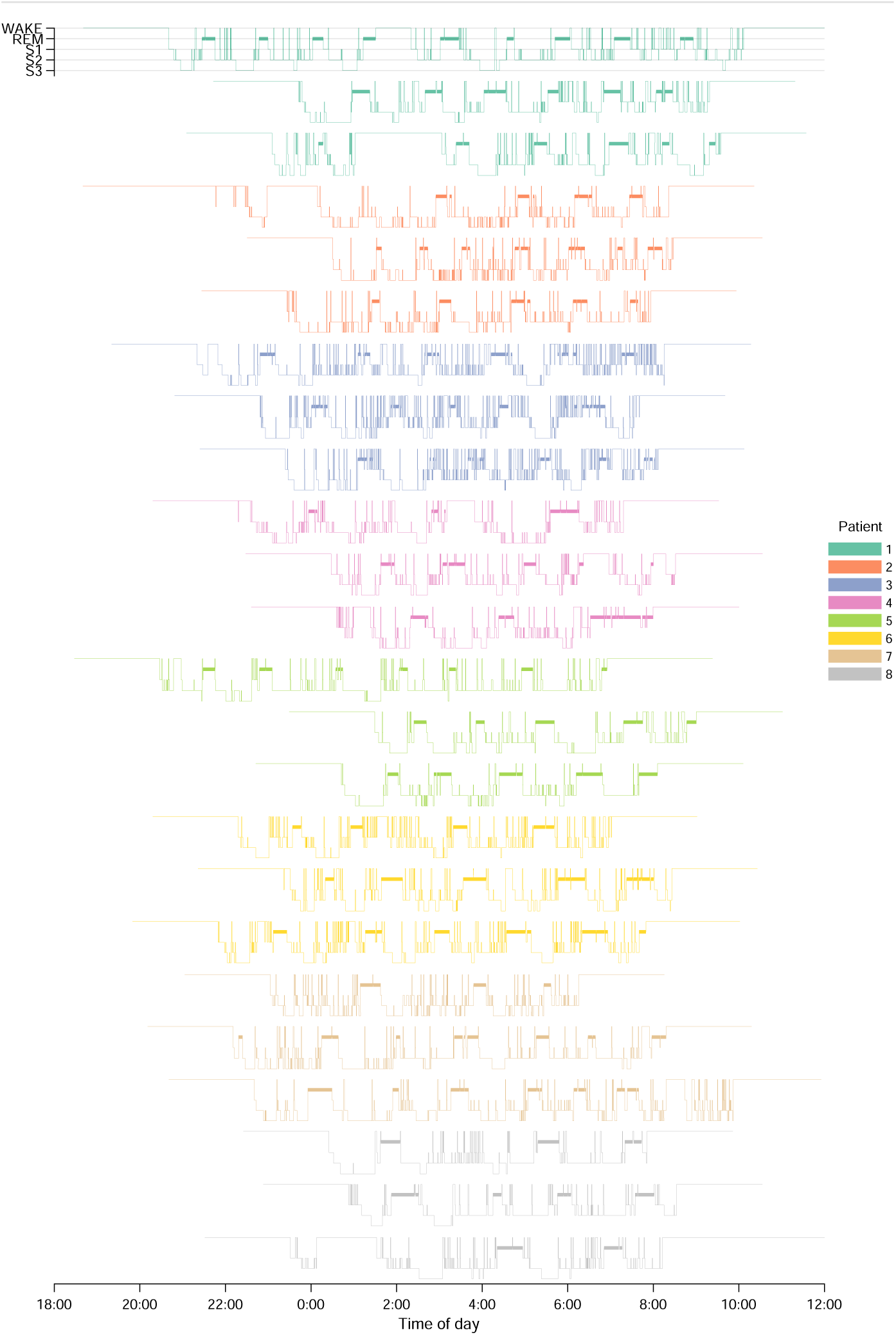
All hypnograms, including 2 hours pre-sleep and post-sleep.

**Fig. S4.**
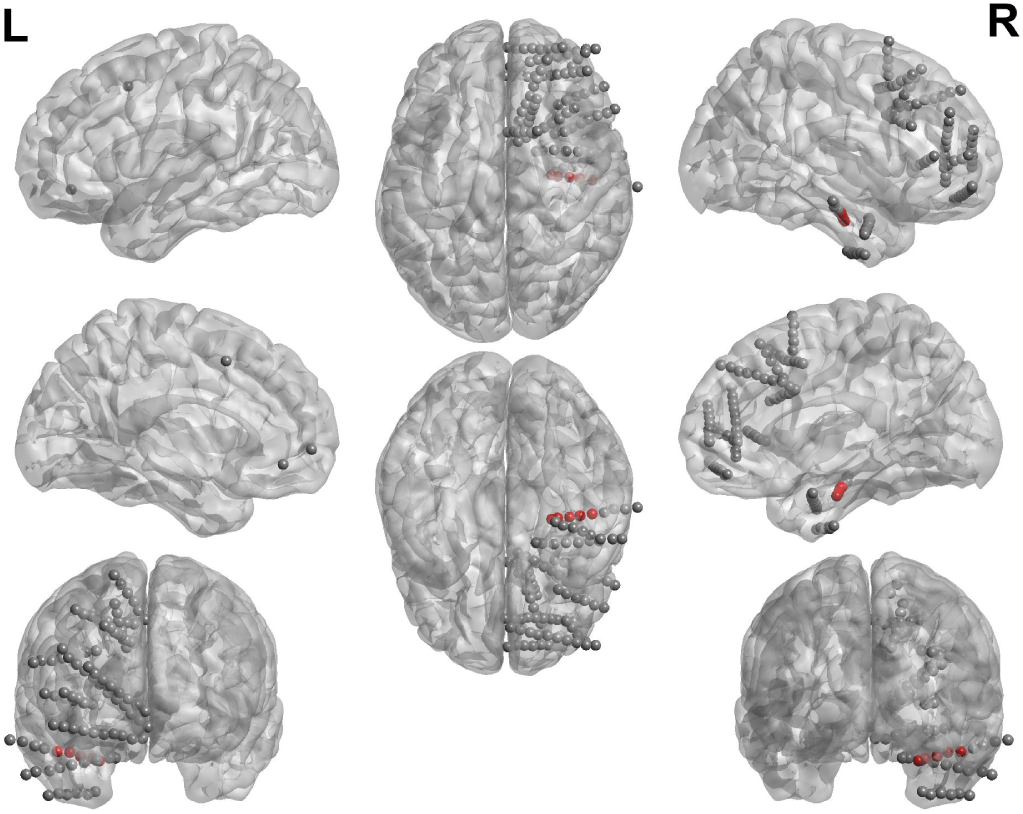
Electrode locations Patient 1. Analysed contacts in red.

**Fig. S5.**
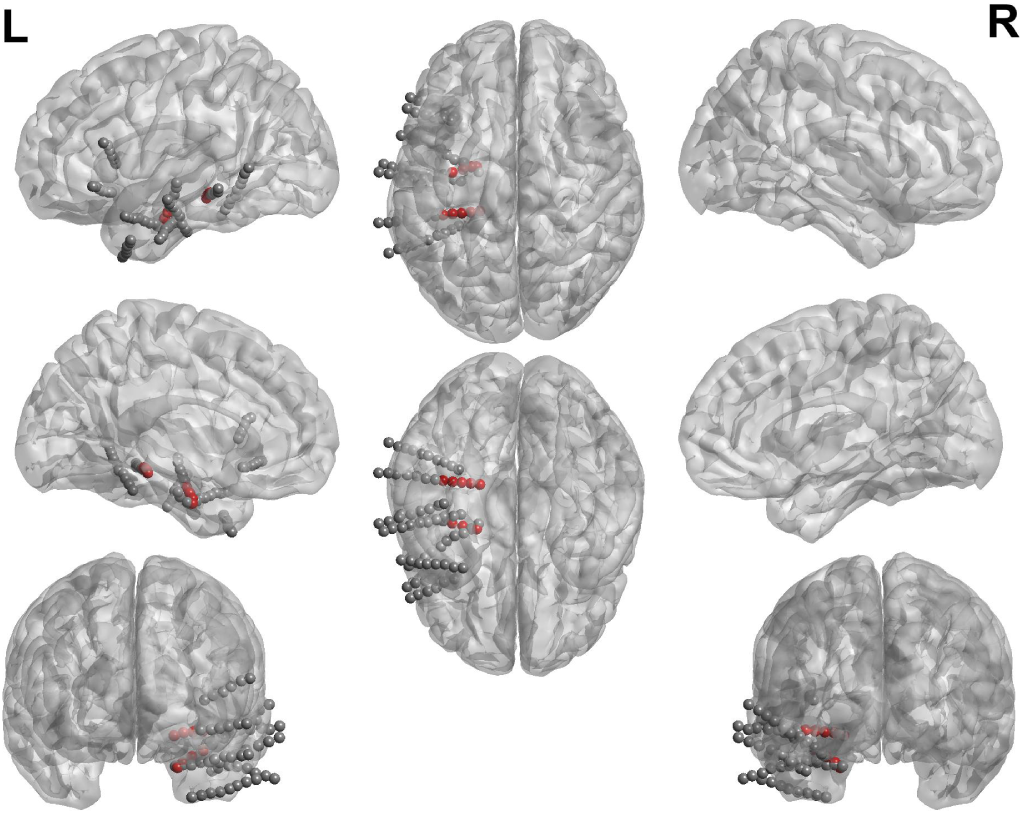
Electrode locations Patient 2. Analysed contacts in red.

**Fig. S6.**
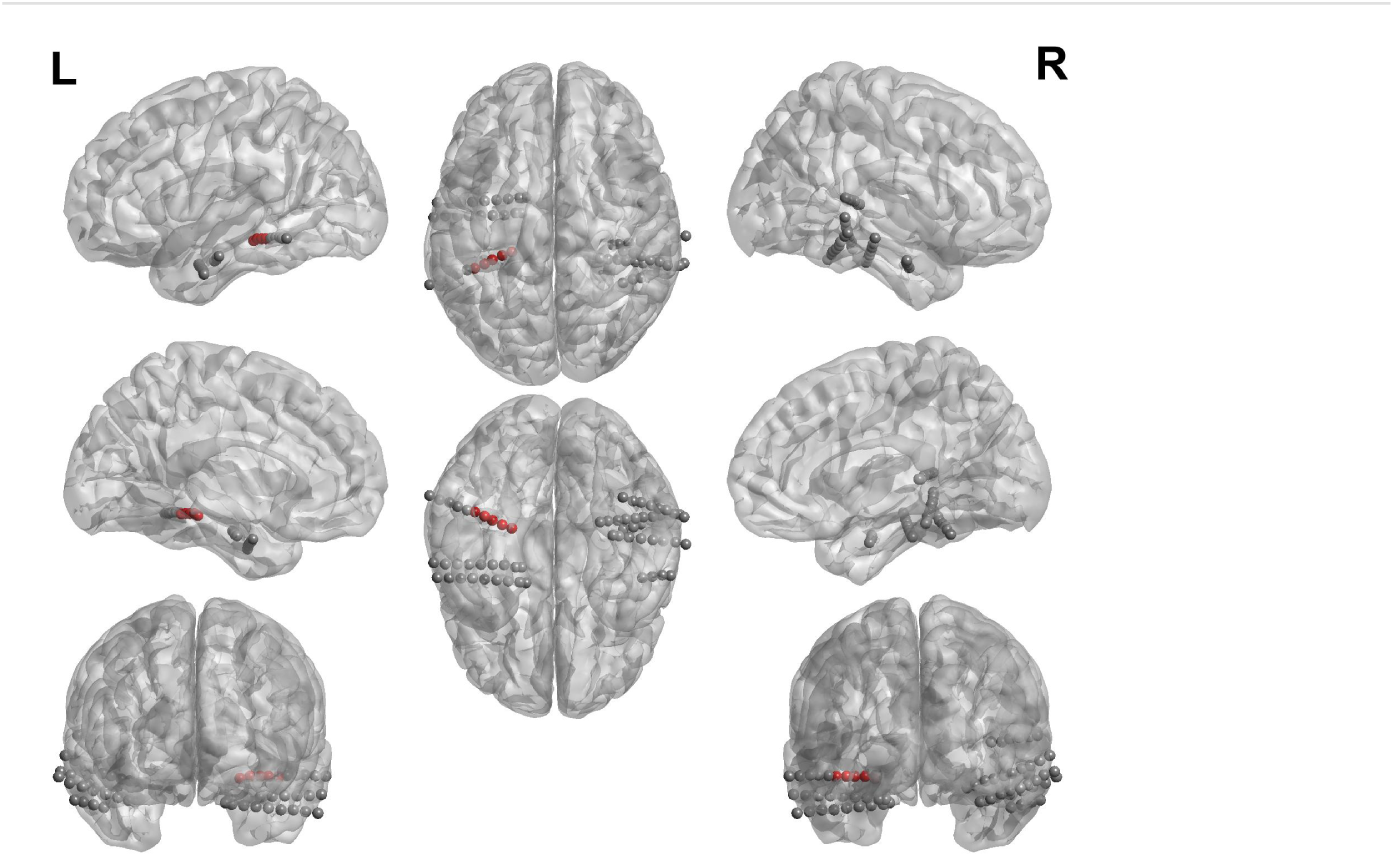
Electrode locations Patient 3. Analysed contacts in red.

**Fig. S7.**
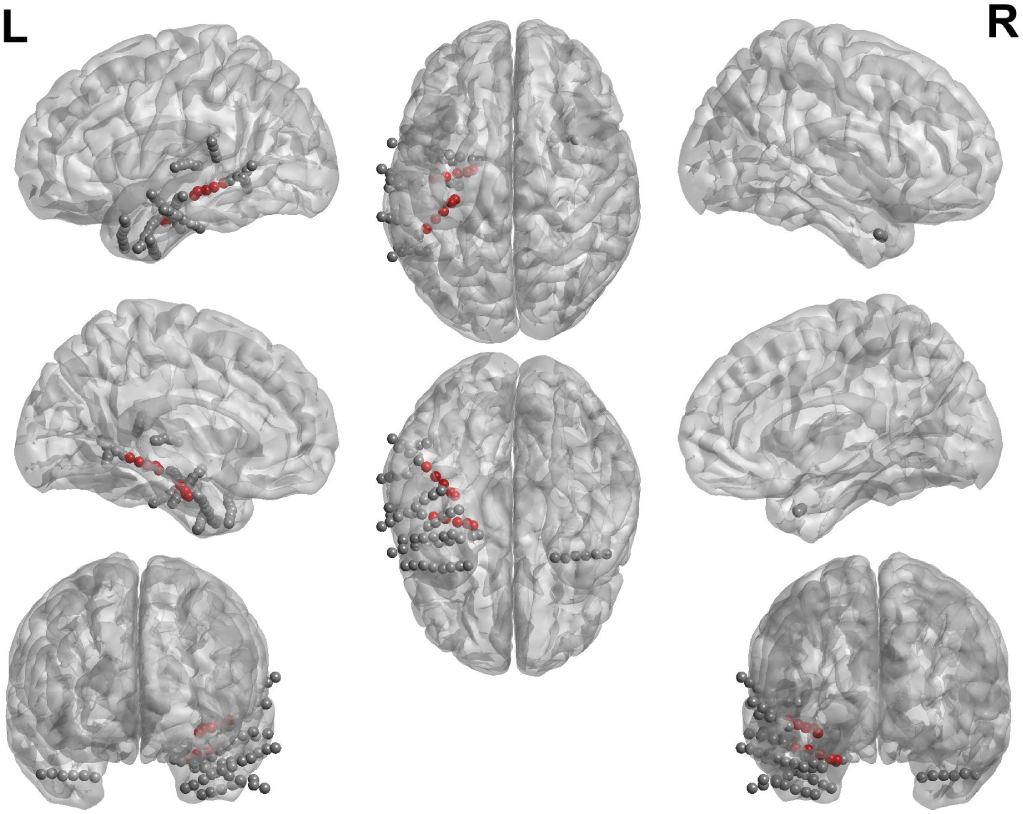
Electrode locations Patient 4. Analysed contacts in red.

**Fig. S8.**
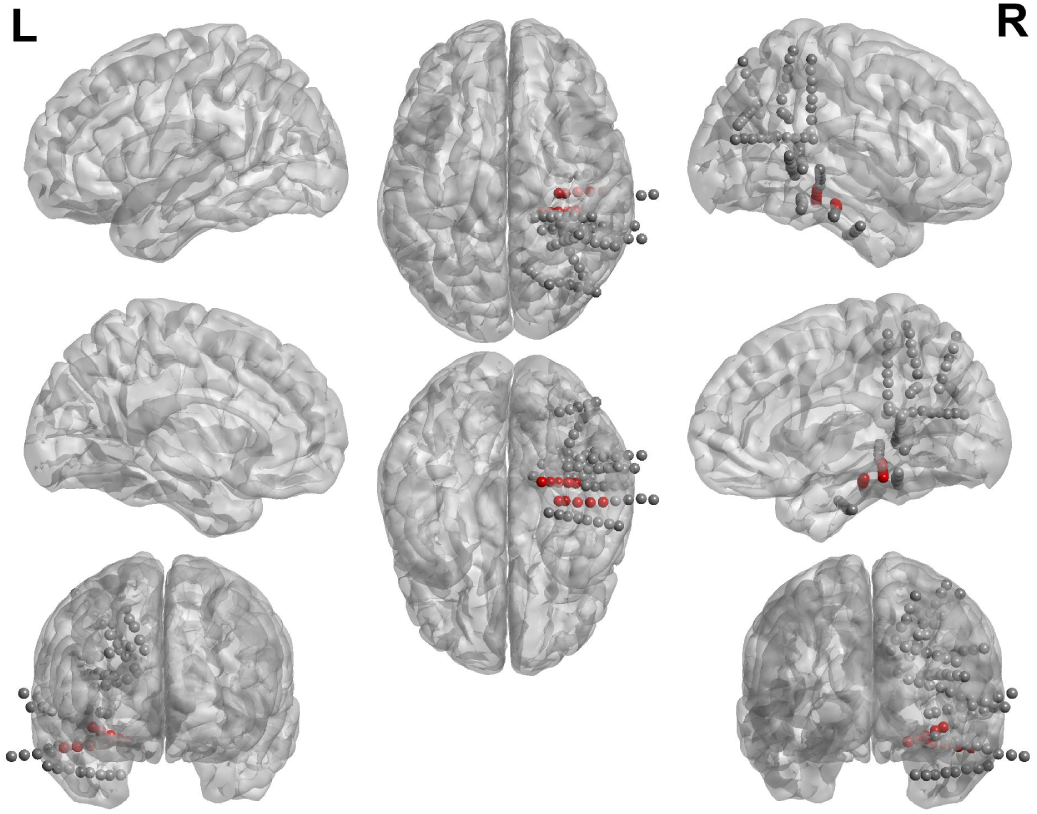
Electrode locations Patient 5. Analysed contacts in red.

**Fig. S9.**
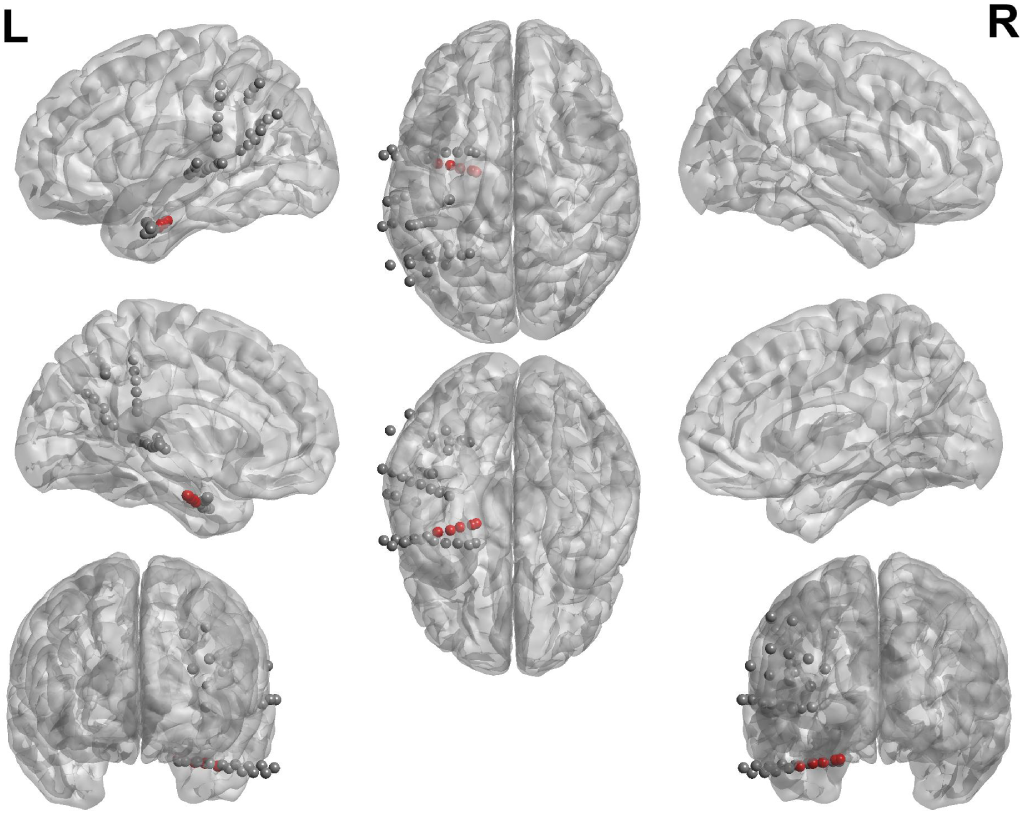
Electrode locations Patient 6. Analysed contacts in red.

**Fig. S10.**
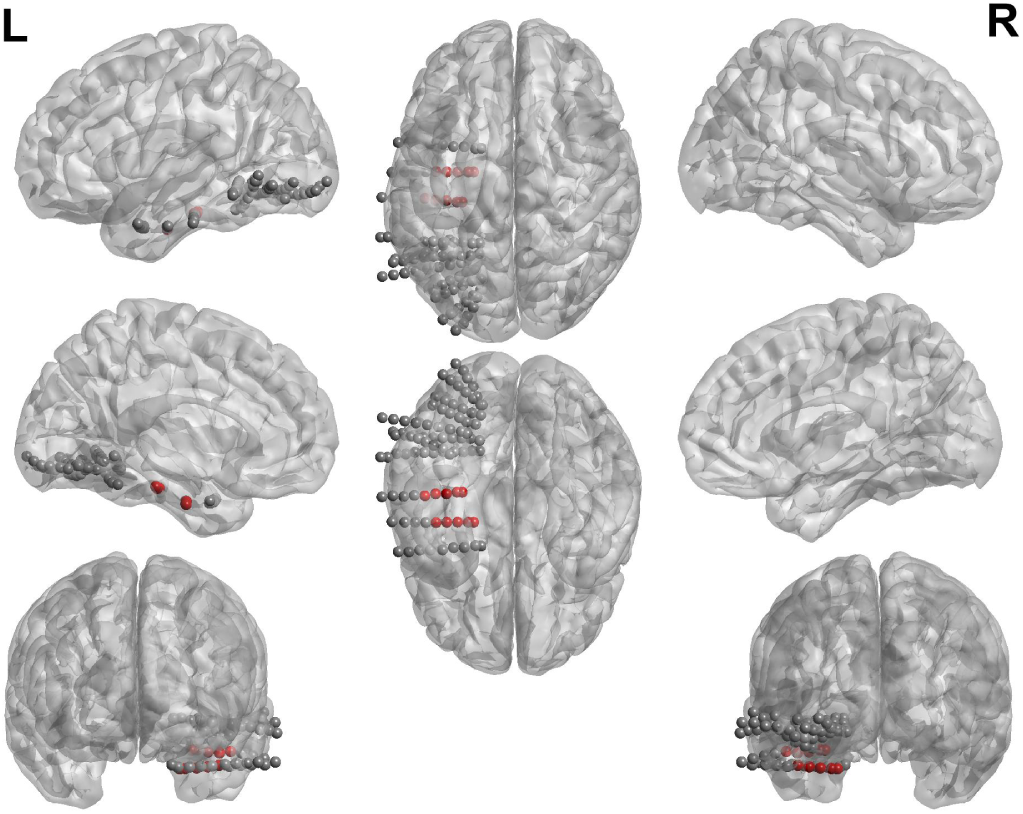
Electrode locations Patient 7. Analysed contacts in red.

**Fig. S11.**
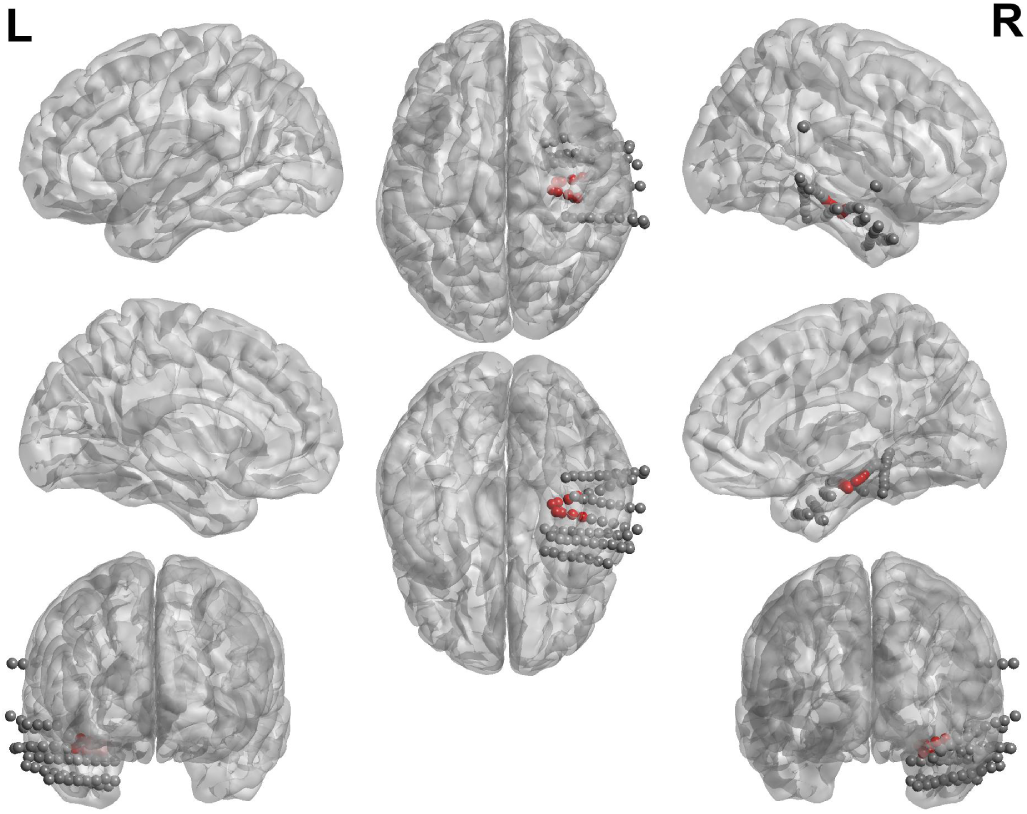
Electrode locations Patient 8. Analysed contacts in red.

**Fig. S12.**
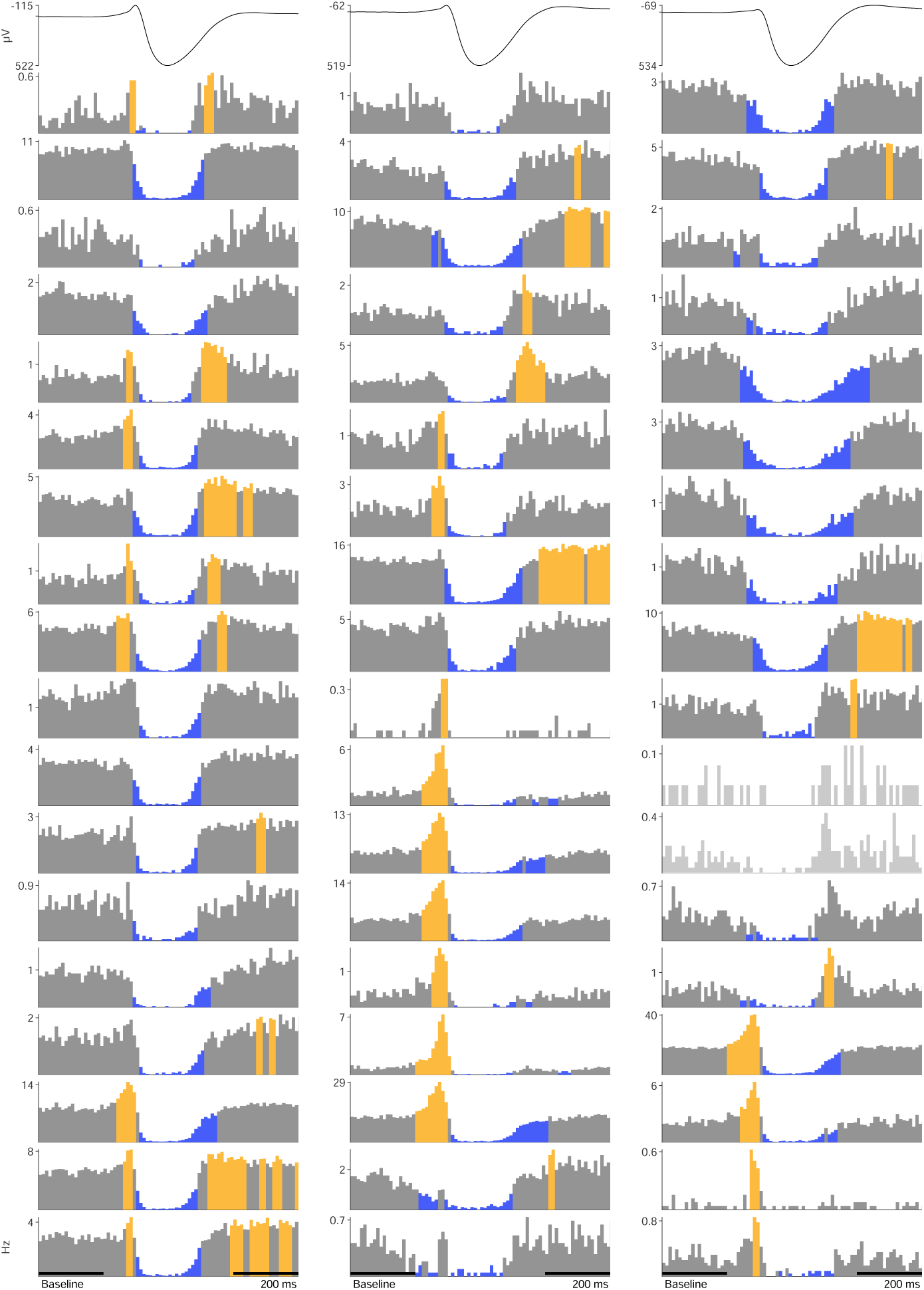
Patient 1 (1/3): PSTH of each unit. Top row shows time-locked LFP. Units are organized over nights (columns). The yellow color indicates significant increases and the blue color decreases compared with the average firing rate during baseline, indicated in the x-axis of bottom left plot.

**Fig. S13.**
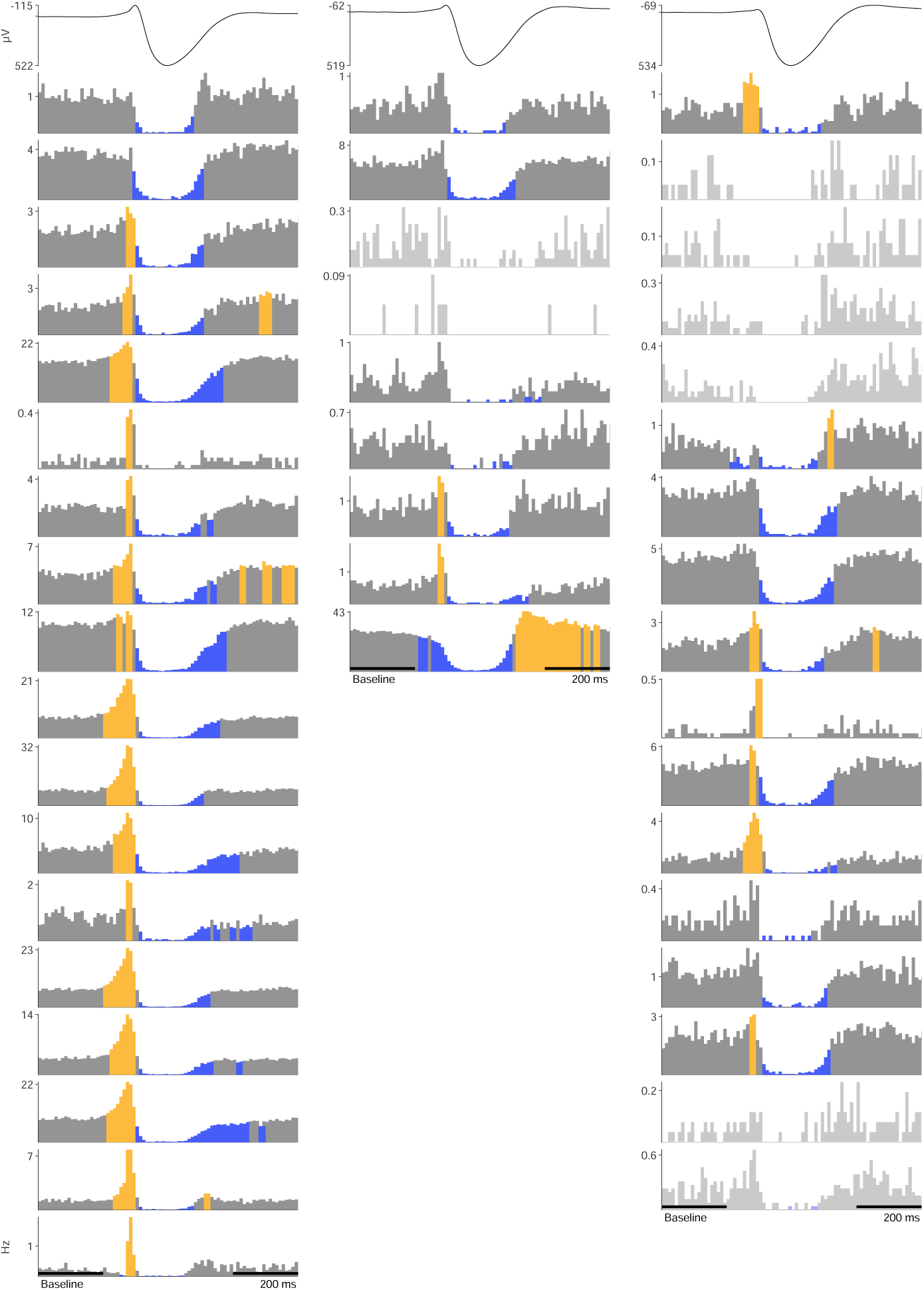
Patient 1 (2/3): PSTH of each unit. Top row shows time-locked LFP. Units are organized over nights (columns). The yellow color indicates significant increases and the blue color decreases compared with the average firing rate during baseline, indicated in the x-axis of bottom left plot.

**Fig. S14.**
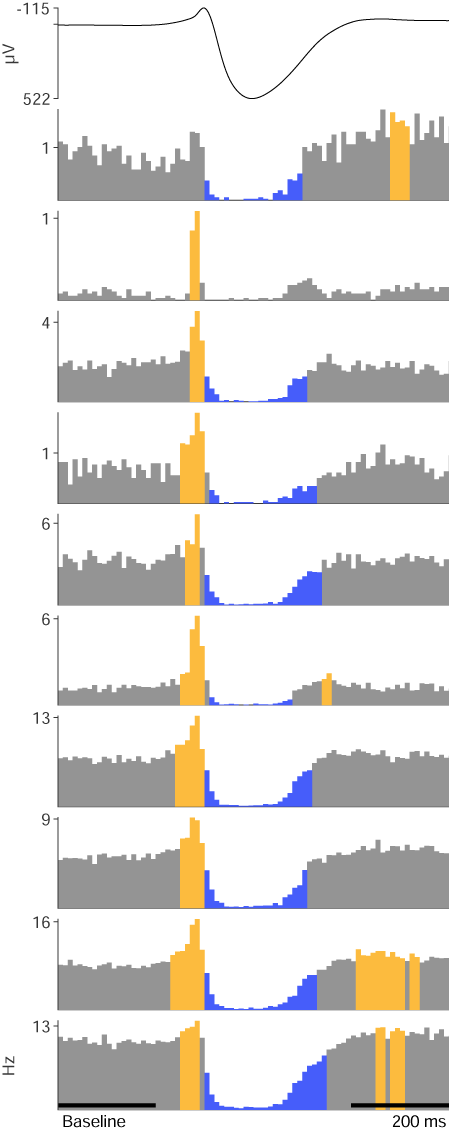
Patient 1 (3/3): PSTH of each unit. Top row shows time-locked LFP. Units are organized over nights (columns). The yellow color indicates significant increases and the blue color decreases compared with the average firing rate during baseline, indicated in the x-axis of bottom left plot.

**Fig. S15.**
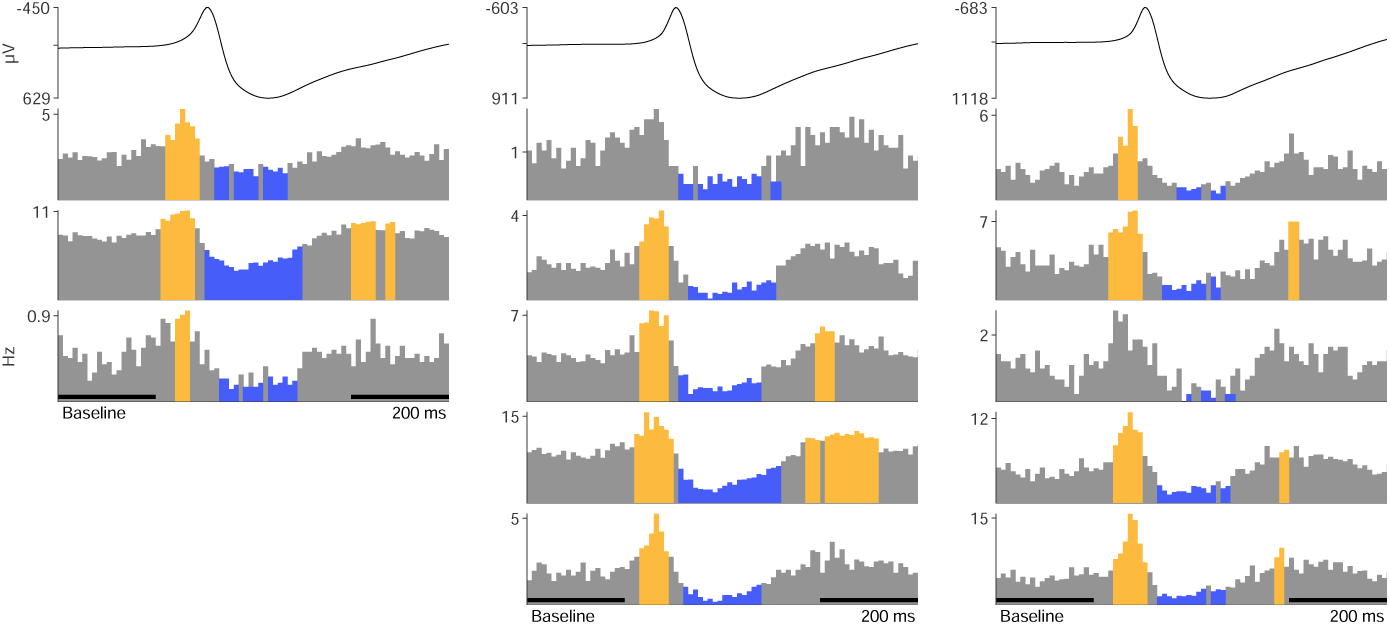
Patient 2: PSTH of each unit. Top row shows time-locked LFP. Units are organized over nights (columns). The yellow color indicates significant increases and the blue color decreases compared with the average firing rate during baseline, indicated in the x-axis of bottom left plot.

**Fig. S16.**
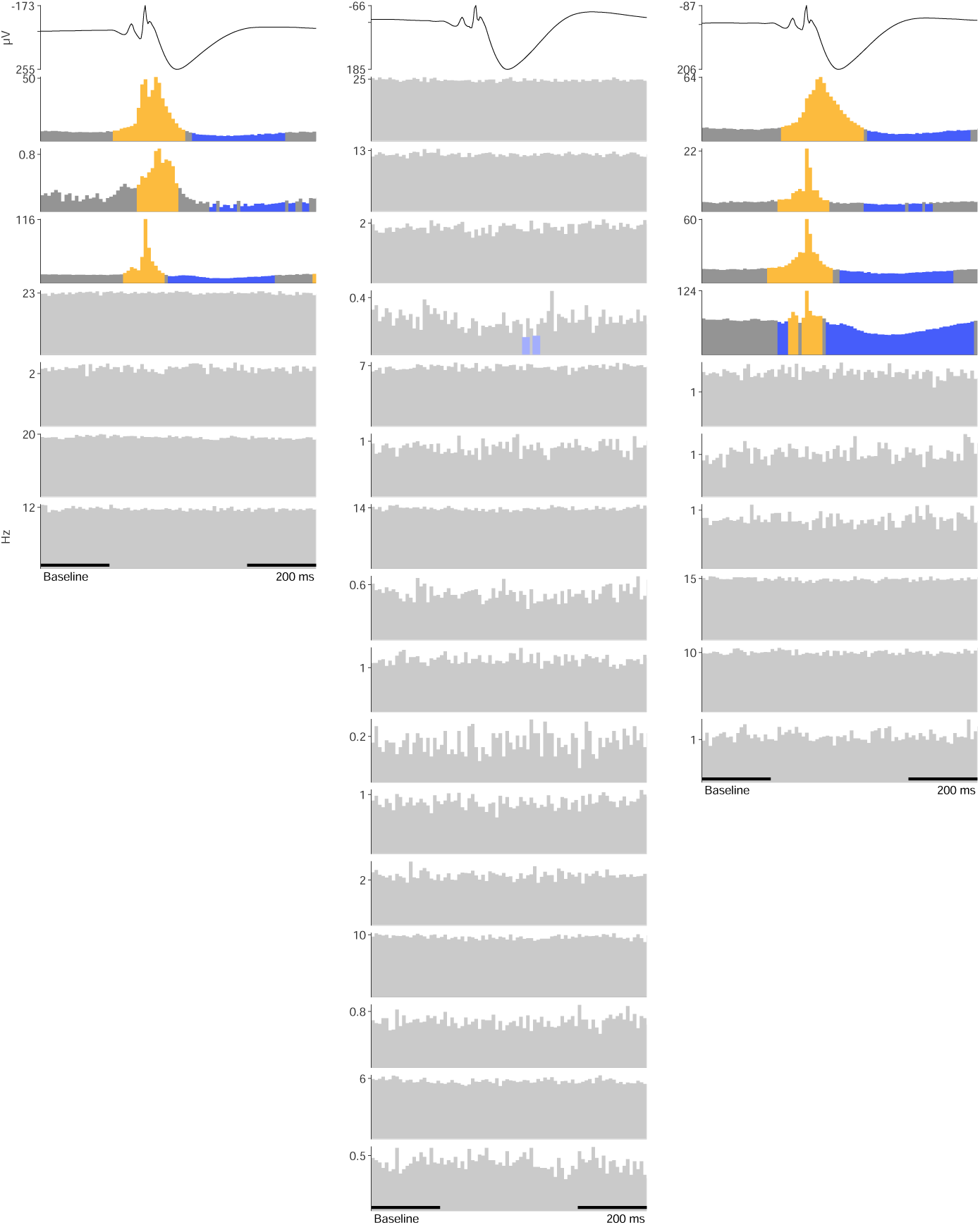
Patient 3: PSTH of each unit. Top row shows time-locked LFP. Units are organized over nights (columns). The yellow color indicates significant increases and the blue color decreases compared with the average firing rate during baseline, indicated in the x-axis of bottom left plot.

**Fig. S17.**
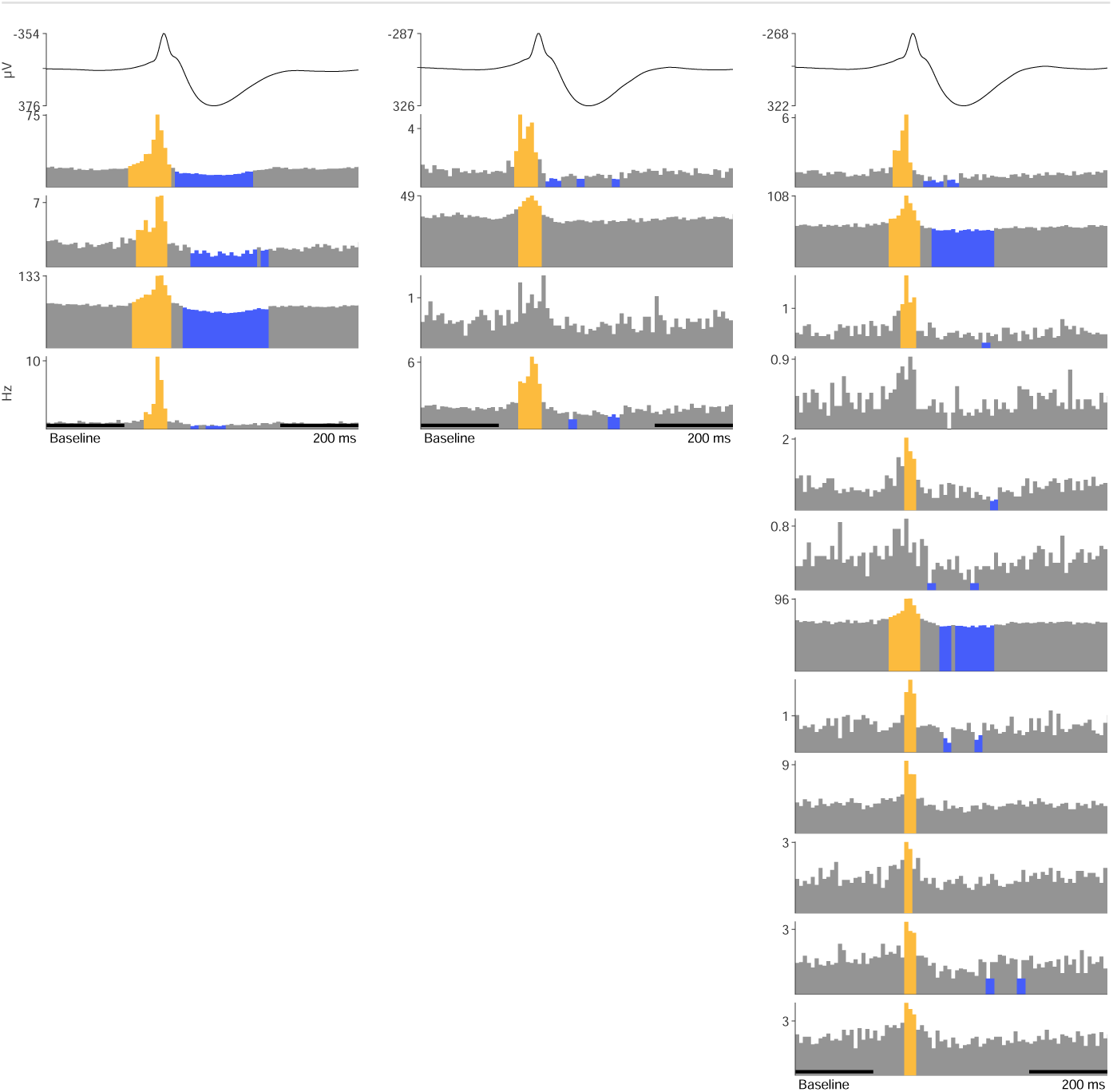
Patient 4: PSTH of each unit. Top row shows time-locked LFP. Units are organized over nights (columns). The yellow color indicates significant increases and the blue color decreases compared with the average firing rate during baseline, indicated in the x-axis of bottom left plot.

**Fig. S18.**
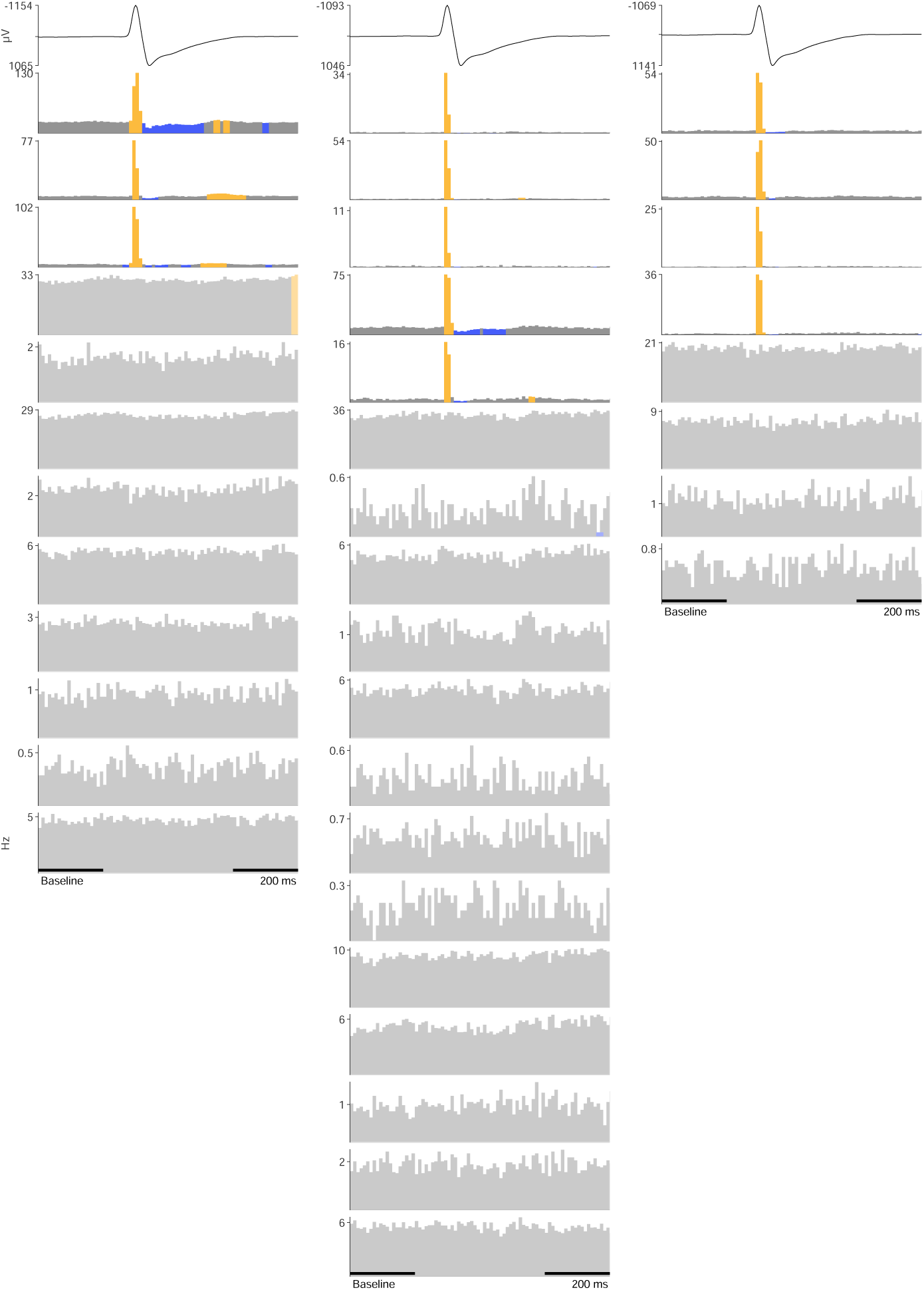
Patient 5: PSTH of each unit. Top row shows time-locked LFP. Units are organized over nights (columns). The yellow color indicates significant increases and the blue color decreases compared with the average firing rate during baseline, indicated in the x-axis of bottom left plot.

**Fig. S19.**
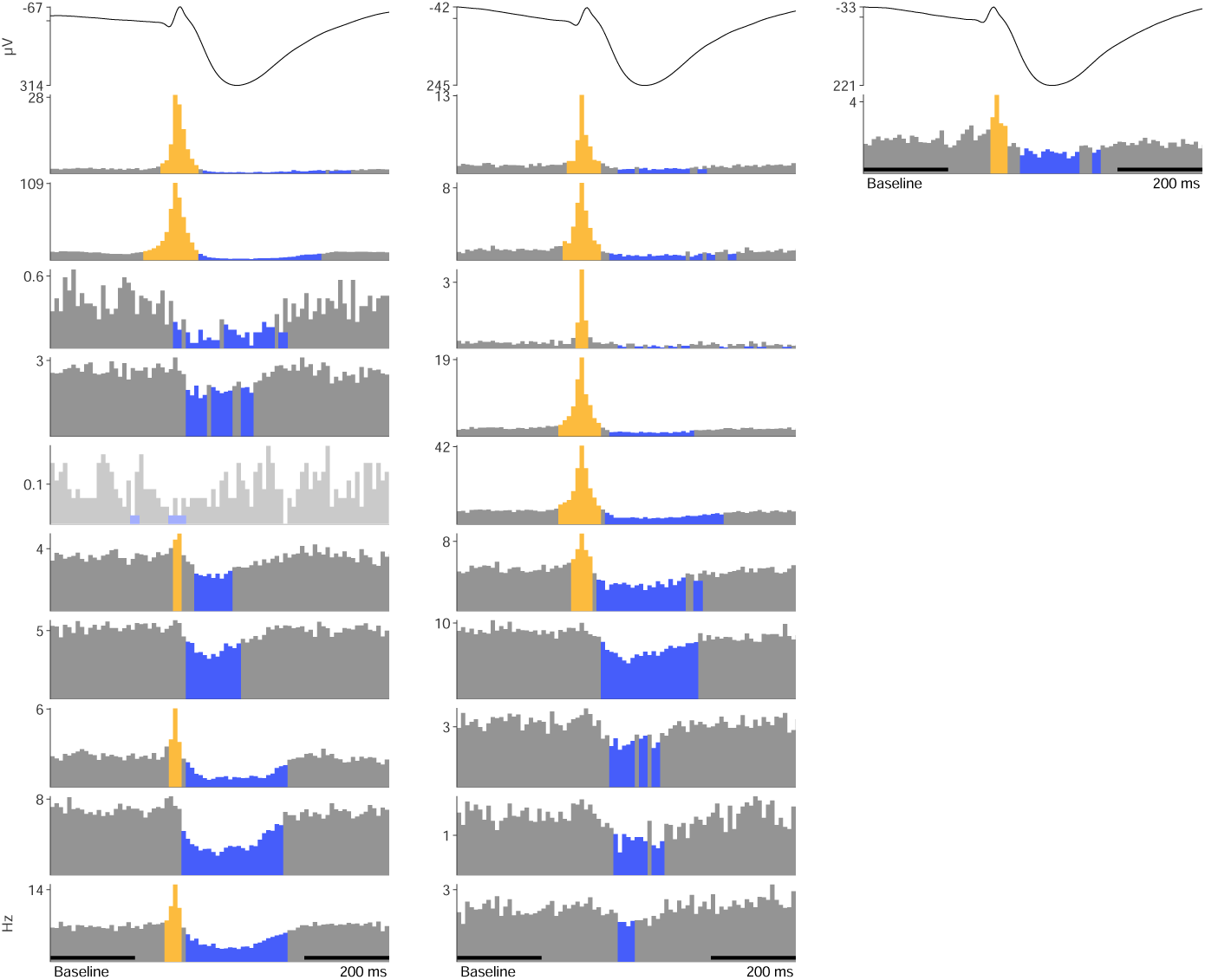
Patient 6: PSTH of each unit. Top row shows time-locked LFP. Units are organized over nights (columns). The yellow color indicates significant increases and the blue color decreases compared with the average firing rate during baseline, indicated in the x-axis of bottom left plot.

**Fig. S20.**
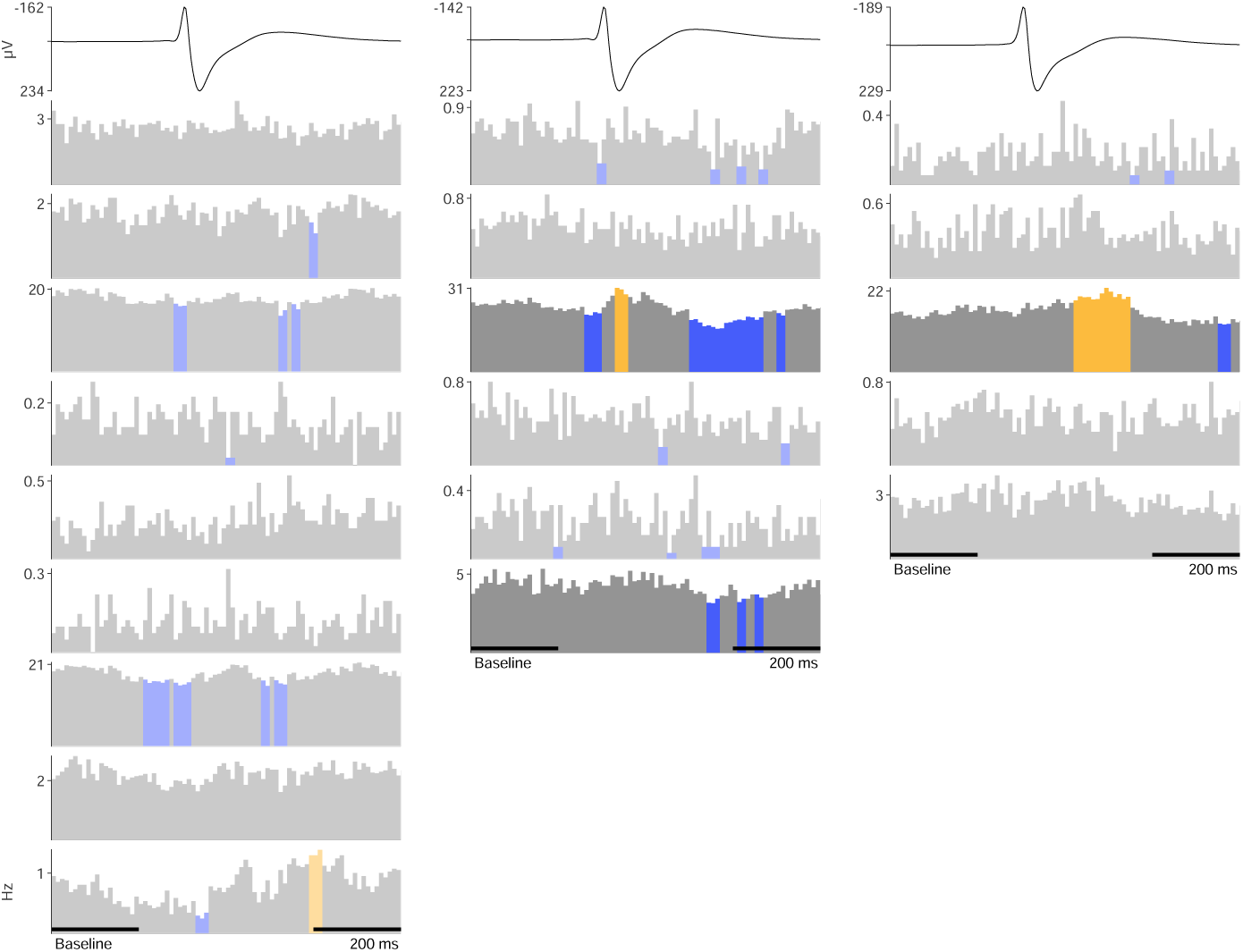
Patient 7: PSTH of each unit. Top row shows time-locked LFP. Units are organized over nights (columns). The yellow color indicates significant increases and the blue color decreases compared with the average firing rate during baseline, indicated in the x-axis of bottom left plot.

**Fig. S21.**
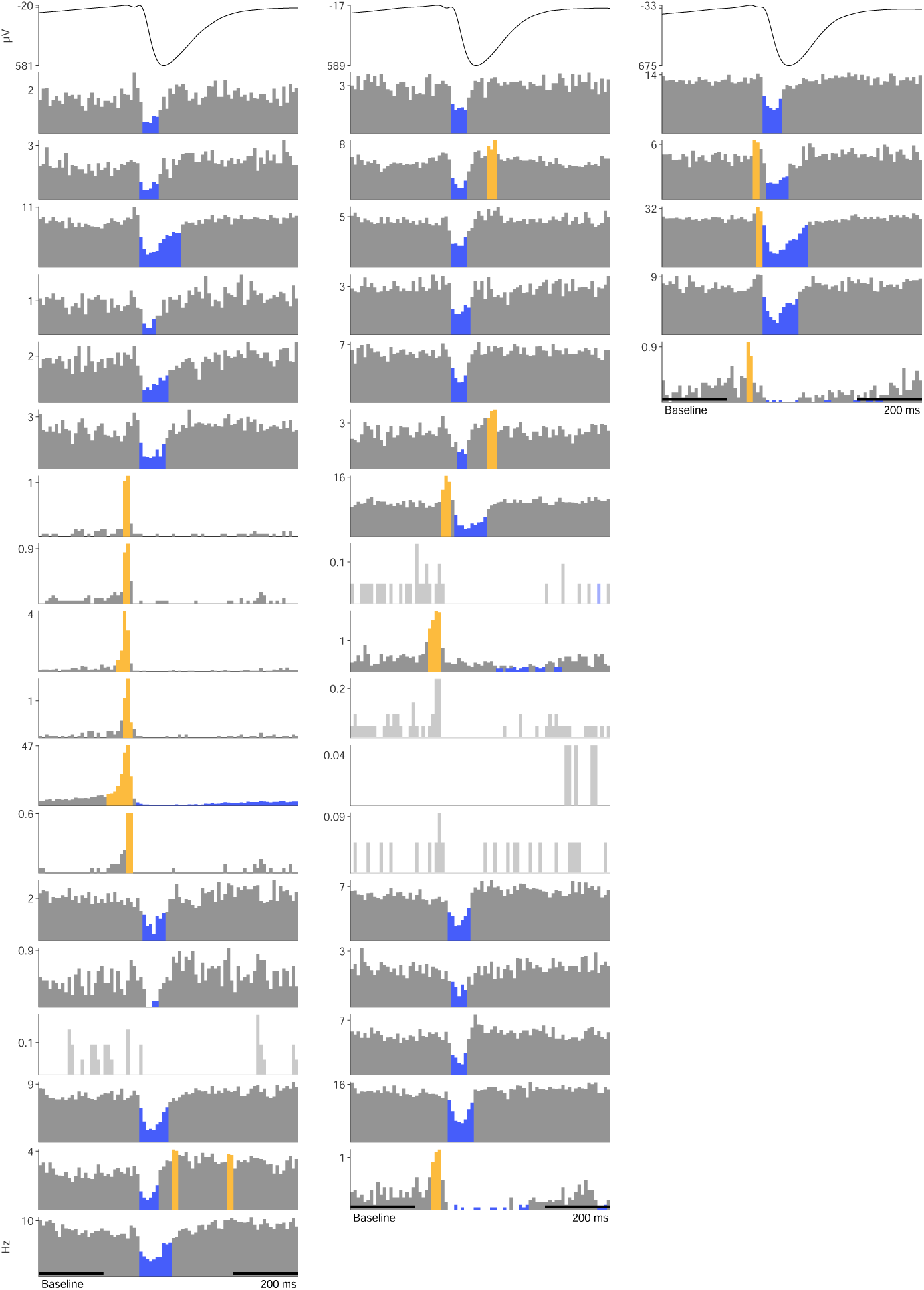
Patient 8: PSTH of each unit. Top row shows time-locked LFP. Units are organized over nights (columns). The yellow color indicates significant increases and the blue color decreases compared with the average firing rate during baseline, indicated in the x-axis of bottom left plot.

## References

1. Gowers W. Course of Epilepsy in Epilepsy and Other Chronic Convulsive Diseases: Their Causes, Symptoms and Treatment:157–164 New York, NY, USA: William Wood 1881.

2. Langdon-Down Mary, Russell Brain W.. Time of Day in Relation to Convulsions in Epilepsy The Lancet. 1929;213:1029–1032.

3. Griffiths Gwenvron M., Fox J. Tylor. Rhythm in Epilepsy The Lancet. 1938;232:409–416.

4. Janz D. The Grand Mal Epilepsies and the Sleeping-Waking Cycle Epilepsia. 1962;3:69–109.

5. Autret Alain, Lucas B., Hommet C., Corcia P., Toffol B. Sleep and the Epilepsies Journal of Neurology. 1997;244:S10–S17.

6. Bazil Carl W., Walczak Thaddeus S.. Effects of Sleep and Sleep Stage on Epileptic and Nonepileptic Seizures Epilepsia. 1997;38:56–62.

7. Shouse M.N., Farber P.R., Staba R.J. Physiological Basis: How NREM Sleep Components Can Promote and REM Sleep Components Can Suppress Seizure Discharge Propagation Clinical Neurophysiology. 2000;111:S9–S18.

8. Herman S.T., Walczak T.S., Bazil C.W. Distribution of Partial Seizures during the Sleep-Wake Cycle: Differences by Seizure Onset Site Neurology. 2001;56:1453–1459.

9. Jobst Barbara C., Williamson Peter D., Neuschwander Timothy B., Darcey Terrance M., Thadani Vijay M., Roberts David W.. Secondarily Generalized Seizures in Mesial Temporal Epilepsy: Clinical Characteristics, Lateralizing Signs, and Association With Sleep–Wake Cycle Epilepsia. 2001;42:1279–1287.

10. Méndez Melissa, Radtke Rodney A.. Interactions Between Sleep and Epilepsy: Journal of Clinical Neurophysiology. 2001;18:106–127.

11. Minecan Daniela, Natarajan Alamelu, Marzec Mary, Malow Beth. Relationship of Epileptic Seizures to Sleep Stage and Sleep Depth Sleep. 2002;25:56–61.

12. Malow Beth A.. Sleep and Epilepsy Neurologic Clinics. 2005;23:1127–1147.

13. Rocamora Rodrigo, Andrzejak Ralph G., Jiménez-Conde Jordi, Elger Christian E.. Sleep Modulation of Epileptic Activity in Mesial and Neocortical Temporal Lobe Epilepsy: A Study with Depth and Subdural Electrodes Epilepsy & Behavior. 2013;28:185–190.

14. Frucht Michael M, Quigg Mark, Schwaner Carl, Fountain Nathan B. Distribution of Seizure Precipitants Among Epilepsy Syndromes Epilepsia. 2000;41:6.

15. Ferlisi Monica, Shorvon Simon. Seizure Precipitants (Triggering Factors) in Patients with Epilepsy Epilepsy & Behavior. 2014;33:101–105.

16. Wassenaar Merel, Kasteleijn-Nolst Trenité Dorothée G. A., de Haan Gerrit-Jan, Carpay Johannes A., Leijten Frans S. S.. Seizure Precipitants in a Community-Based Epilepsy Cohort Journal of Neurology. 2014;261:717–724.

17. Cobabe Maurine M., Sessler Daniel I., Nowacki Amy S., O’Rourke Colin, Andrews Noah, Foldvary-Schaefer Nancy. Impact of Sleep Duration on Seizure Frequency in Adults with Epilepsy: A Sleep Diary Study Epilepsy & Behavior. 2015;43:143–148.

18. Reddy Doodipala Samba, Chuang Shu-Hui, Hunn Dayton, Crepea Amy Z., Magant Rama. Neuroendocrine Aspects of Improving Sleep in Epilepsy Epilepsy research. 2018;147:32–41.

19. Foldvary-Schaefer Nancy, Grigg-Damberger Madeleine. Sleep and Epilepsy: What We Know, Don’t Know, and Need to Know Journal of Clinical Neurophysiology. 2006;23:4–20.

20. Jain Sejal V., Kothare Sanjeev V.. Sleep and Epilepsy Seminars in Pediatric Neurology. 2015;22:86–92.

21. Kataria Lynn, Vaughn Bradley V. Sleep and Epilepsy Sleep Medicine Clinics. 2016;11:25–38.

22. Derry Christopher P., Duncan Susan. Sleep and Epilepsy Epilepsy & Behavior. 2013;26:394–404.

23. Staley Kevin J, Dudek F Edward. Interictal Spikes and Epileptogenesis Epilepsy Currents. 2006;6:199–202.

24. Gibbs E. L., Gibbs F. A. Diagnostic and Localizing Value of Electroencephalographic Studies in Sleep Res Publ Assoc Res Nerv Ment Dis. 1947;26:366–376.

25. Penfield Wilder, Jasper Herbert. Epilepsy and the Functional Anatomy of the Human Brain. Epilepsy and the Functional Anatomy of the Human Brain Oxford, England: Little, Brown & Co. 1954.

26. Lieb J. P., Joseph J. P., Engel J., Walker J., Crandall P. H. Sleep State and Seizure Foci Related to Depth Spike Activity in Patients with Temporal Lobe Epilepsy Electroencephalography and Clinical Neurophysiology. 1980;49:538–557.

27. Ferrillo F., Beelke M., De Carli F., et al. Sleep-EEG Modulation of Interictal Epileptiform Discharges in Adult Partial Epilepsy: A Spectral Analysis Study Clinical Neurophysiology. 2000;111:916–923.

28. Bagshaw Andrew P., Jacobs Julia, LeVan Pierre, Dubeau François, Gotman Jean. Effect of Sleep Stage on Interictal High-Frequency Oscillations Recorded from Depth Macroelectrodes in Patients with Focal Epilepsy Epilepsia. 2009;50:617–628.

29. Goncharova Irina I., Zaveri Hitten P., Duckrow Robert B., Novotny Edward J., Spencer Susan S.. Spatial Distribution of Intracranially Recorded Spikes in Medial and Lateral Temporal Epilepsies Epilepsia. 2009;50:2575–2585.

30. Karoly Philippa J., Freestone Dean R., Boston Ray, et al. Interictal Spikes and Epileptic Seizures: Their Relationship and Underlying Rhythmicity Brain. 2016;139:1066–1078.

31. Rossi G.F, Colicchio G, Pola P. Interictal Epileptic Activity during Sleep: A Stereo-EEG Study in Patients with Partial Epilepsy Electroencephalography and Clinical Neurophysiology. 1984;58:97–106.

32. Sammaritano Michele, Gigli Gian Luigi, Gotman Jean. Interictal Spiking during Wakefulness and Sleep and the Localization of Foci in Temporal Lobe Epilepsy Neurology. 1991;41:290–290.

33. Spencer David C., Sun Felice T., Brown Sarah N., et al. Circadian and Ultradian Patterns of Epileptiform Discharges Differ by Seizure-Onset Location during Long-Term Ambulatory Intracranial Monitoring Epilepsia. 2016;57:1495–1502.

34. Lambert Isabelle, Roehri Nicolas, Giusiano Bernard, et al. Brain Regions and Epileptogenicity Influence Epileptic Interictal Spike Production and Propagation during NREM Sleep in Comparison with Wakefulness Epilepsia. 2018;59:235–243.

35. Fouad Amal, Azizollahi Hamed, Le Douget Jean-Eudes, et al. Interictal Epileptiform Discharges Show Distinct Spatiotemporal and Morphological Patterns across Wake and Sleep Brain Communications. 2022;4:fcac183.

36. Anderson Christopher T., Tcheng Thomas K., Sun Felice T., Morrell Martha J.. Day–Night Patterns of Epileptiform Activity in 65 Patients With Long-Term Ambulatory Electrocorticography: Journal of Clinical Neurophysiology. 2015;32:406–412.

37. Baud Maxime O., Kleen Jonathan K., Mirro Emily A., et al. Multi-Day Rhythms Modulate Seizure Risk in Epilepsy Nature Communications. 2018;9.

38. Ng Marcus, Pavlova Milena. Why Are Seizures Rare in Rapid Eye Movement Sleep? Review of the Frequency of Seizures in Different Sleep Stages Epilepsy Research and Treatment. 2013;2013:1–10.

39. Malow Beth A., Lin Xihong, Kushwaha Ramesh, Aldrich Michael S. Interictal Spiking Increases with Sleep Depth in Temporal Lobe Epilepsy Epilepsia. 1998;39:1309–1316.

40. Fürbass Franz, Koren Johannes, Hartmann Manfred, Brandmayr Georg, Hafner Sebastian, Baumgartner Christoph. Activation Patterns of Interictal Epileptiform Discharges in Relation to Sleep and Seizures: An Artificial Intelligence Driven Data Analysis Clinical Neurophysiology. 2021;132:1584–1592.

41. McNamara J. O. Cellular and Molecular Basis of Epilepsy Journal of Neuroscience. 1994;14:3413–3425.

42. de Curtis Marco, Avanzini Giuliano. Interictal Spikes in Focal Epileptogenesis Progress in Neurobiology. 2001;63:541–567.

43. Ulbert Istvan, Heit Gary, Madsen Joseph, Karmos George, Halgren Eric. Laminar Analysis of Human Neocortical Interictal Spike Generation and Propagation: Current Source Density and Multiunit Analysis In Vivo Epilepsia. 2004;45:48–56.

44. Matsumoto H, Marsan C. Ajmone. Cortical Cellular Phenomena in Experimental Epilepsy: Interictal Manifestations Experimental Neurology. 1964;9:286–304.

45. Prince David A.. Inhibition in “Epileptic” Neurons Experimental Neurology. 1968;21:307–321.

46. Alvarado-Rojas Catalina, Lehongre Katia, Bagdasaryan Juliana, et al. Single-Unit Activities during Epileptic Discharges in the Human Hippocampal Formation Frontiers in Computational Neuroscience. 2013;7:140.

47. Keller Corey J., Truccolo Wilson, Gale John T., et al. Heterogeneous Neuronal Firing Patterns during Interictal Epileptiform Discharges in the Human Cortex Brain. 2010;133:1668–1681.

48. Fabó Dániel, Maglóczky Zsófia, Wittner Lucia, et al. Properties of in Vivo Interictal Spike Generation in the Human Subiculum Brain. 2008;131:485–499.

49. Nayak Chetan S., Mariyappa N., Majumdar Kaushik K., et al. NREM Sleep and Antiepileptic Medications Modulate Epileptiform Activity by Altering Cortical Synchrony Clinical EEG and Neuroscience. 2018;49:417–424.

50. Malow Beth A, Aldrich Michael S. Localizing Value of Rapid Eye Movement Sleep in Temporal Lobe Epilepsy Sleep Medicine. 2000;1:57–60.

51. Timofeev Igor, Grenier François, Steriade Mircea. Disfacilitation and Active Inhibition in the Neocortex during the Natural Sleep-Wake Cycle: An Intracellular Study Proceedings of the National Academy of Sciences of the United States of America. 2001;98:1924–1929.

52. Staba Richard J., Wilson Charles L., Fried Itzhak, Engel Jerome. Single Neuron Burst Firing in the Human Hippocampus during Sleep Hippocampus. 2002;12:724–734.

53. Vyazovskiy Vladyslav V, Olcese Umberto, Lazimy Yaniv M, et al. Cortical Firing and Sleep Homeostasis Neuron. 2009;63:865–878.

54. Aanestad Eivind, Brøgger Jan Christian. S84. Variability of Epileptiform Discharges by Age Using SCORE Clinical Neurophysiology. 2018;129:e173.

55. Aanestad Eivind, Gilhus Nils Erik, Brogger Jan. Interictal Epileptiform Discharges Vary across Age Groups Clinical Neurophysiology. 2020;131:25–33.

56. Beaumanoir A., Ballis T., Varfis G., Ansari K. Benign Epilepsy of Childhood With Rolandic Spikes Epilepsia. 1974;15:301–315.

57. Blom S., Heijbel J. Benign Epilepsy of Children with Centro-Temporal EEG Foci. Discharge Rate during Sleep Epilepsia. 1975;16:133–140.

58. Dalla Bernardina B., Sgrò V., Caraballo R., et al. Sleep and Benign Partial Epilepsies of Childhood: EEG and Evoked Potentials Study Epilepsy Research. Supplement. 1991;2:83–96.

59. Clemens B., Oláh R. Sleep Studies in Benign Epilepsy of Childhood with Rolandic Spikes. I. Sleep Pathology Epilepsia. 1987;28:20–23.

60. Hufnagel A., Dumpelmann M., Zentner J., Schijns O., Elger C. E. Clinical Relevance of Quantified Intracranial Interictal Spike Activity in Presurgical Evaluation of Epilepsy Epilepsia. 2000;41:467–478.

61. Frost James D., Hrachovy Richard A., Glaze Daniel G., McCully Michael I.. Sleep Modulation of Interictal Spike Configuration in Untreated Children with Partial Seizures Epilepsia. 1991;32:341–346.

62. Del Felice Alessandra, Storti Silvia Francesca, Manganotti Paolo. Sleep Affects Cortical Source Modularity in Temporal Lobe Epilepsy: A High-Density EEG Study Clinical Neurophysiology. 2015;126:1677–1683.

63. Liu Yung-Chun, Lin Chou-Ching K., Tsai Jing-Jane, Sun Yung-Nien. Model-Based Spike Detection of Epileptic EEG Data Sensors (Basel, Switzerland). 2013;13:12536–12547.

64. Ponce-Alvarez Adrián, Kilavik Bjørg Elisabeth, Riehle Alexa. Relating Firing Rate and Spike Time Irregularity in Motor Cortical Neurons BMC Neuroscience. 2009;10:P298.

65. Gotman J. Quantitative Measurements of Epileptic Spike Morphology in the Human EEG Electroencephalography and Clinical Neurophysiology. 1980;48:551–557.

66. Altafullah I, Halgren E, Stapleton J. M, Crandall P. H. Interictal Spike-Wave Complexes in the Human Medial Temporal Lobe: Typical Topography and Comparisons with Cognitive Potentials Electroencephalography and Clinical Neurophysiology. 1986;63:503–516.

67. Hofstra Wytske AElig, de Weerd Al Wytze. The Circadian Rhythm and Its Interaction with Human Epilepsy: A Review of Literature Sleep Medicine Reviews. 2009;13:413–420.

68. Daley Joseph T., DeWolfe Jennifer L.. Sleep, Circadian Rhythms, and Epilepsy Current Treatment Options in Neurology. 2018;20:47.

69. Khan Sofia, Nobili Lino, Khatami Ramin, et al. Circadian Rhythm and Epilepsy The Lancet Neurology. 2018.

70. Durazzo T. S., Spencer S. S., Duckrow R. B., Novotny E. J., Spencer D. D., Zaveri H. P. Temporal Distributions of Seizure Occurrence from Various Epileptogenic Regions Neurology. 2008;70:1265–1271.

71. Karafin Matthew, St. Louis Erik K., Zimmerman M. Bridget, Sparks Jon David, Granner Mark A. Bimodal Ultradian Seizure Periodicity in Human Mesial Temporal Lobe Epilepsy Seizure. 2010;19:347–351.

72. Hofstra Wytske A., Spetgens Willy P.J., Leijten Frans S.S., et al. Diurnal Rhythms in Seizures Detected by Intracranial Electrocorticographic Monitoring: An Observational Study Epilepsy & Behavior. 2009;14:617–621.

73. Hofstra Wytske A., Grootemarsink Bertine E., Dieker Rianneke, Palen Job Van Der, Weerd Al W. De. Temporal Distribution of Clinical Seizures over the 24-h Day: A Retrospective Observational Study in a Tertiary Epilepsy Clinic Epilepsia. 2009;50:2019–2026.

74. Pavlova Milena K., Shea Steven A., Scheer Frank A.J.L., Bromfield Edward B. Is There a Circadian Variation of Epileptiform Abnormalities in Idiopathic Generalized Epilepsy? Epilepsy & Behavior. 2009;16:461–467.

75. Badawy Radwa, Macdonell Richard, Jackson Graeme, Berkovic Samuel. The Peri-Ictal State: Cortical Excitability Changes within 24 h of a Seizure Brain. 2009;132:1013–1021.

76. Quigg Mark. Circadian Rhythms: Interactions with Seizures and Epilepsy Epilepsy Research. 2000;42:43–55.

77. Quigg Mark, Clayburn Hope, Straume Martin, Menaker Michael, Bertram Edward H. Effects of Circadian Regulation and Rest-Activity State on Spontaneous Seizures in a Rat Model of Limbic Epilepsy Epilepsia. 2000;41:502–509.

78. Nzwalo Hipólito, Menezes Cordeiro Inês, Santos Ana Catarina, Peralta Rita, Paiva Teresa, Bentes Carla. 24-Hour Rhythmicity of Seizures in Refractory Focal Epilepsy Epilepsy & Behavior. 2016;55:75–78.

79. Mirzoev Alexander, Bercovici Eduard, Stewart Lee S., Cortez Miguel A., Snead O. Carter, Desrocher Mary. Circadian Profiles of Focal Epileptic Seizures: A Need for Reappraisal Seizure. 2012;21:412–416.

80. Smyk Magdalena K., van Luijtelaar Gilles. Circadian Rhythms and Epilepsy: A Suitable Case for Absence Epilepsy Frontiers in Neurology. 2020;0.

81. Steriade M, Contreras D. Relations between Cortical and Thalamic Cellular Events during Transition from Sleep Patterns to Paroxysmal Activity The Journal of Neuroscience. 1995;15:623–642.

82. Steriade Mircea, Amzica Florin, Neckelmann Dag, Timofeev Igor. Spike-Wave Complexes and Fast Components of Cortically Generated Seizures. II. Extra– and Intracellular Patterns Journal of Neurophysiology. 1998;80:1456–1479.

83. Steriade Mircea. Sleep, Epilepsy and Thalamic Reticular Inhibitory Neurons Trends in Neurosciences. 2005;28:317–324.

84. Halász Peter. How Sleep Activates Epileptic Networks? Epilepsy Research and Treatment. 2013;2013.

85. Busek Petr, Buskova Jitka, Nevsimalova Sona. Interictal Epileptiform Discharges and Phasic Phenomena of REM Sleep Epileptic Disorders: International Epilepsy Journal with Videotape. 2010;12:217–221.

86. Okanari Kazuo, Baba Shiro, Otsubo Hiroshi, et al. Rapid Eye Movement Sleep Reveals Epileptogenic Spikes for Resective Surgery in Children with Generalized Interictal Discharges Epilepsia. 2015;56:1445–1453.

87. Scarlatelli-Lima Aline Vieira, Sukys-Claudino Lucia, Watanabe Nancy, Guarnieri Ricardo, Walz Roger, Lin Katia. How Do People with Drug-Resistant Mesial Temporal Lobe Epilepsy Sleep? A Clinical and Video-EEG with EOG and Submental EMG for Sleep Staging Study eNeurologicalSci. 2016;4:34–41.

88. Janca Radek, Krsek Pavel, Jezdik Petr, et al. The Sub-Regional Functional Organization of Neocortical Irritative Epileptic Networks in Pediatric Epilepsy Frontiers in Neurology. 2018;9:184.

89. Conrad Erin C, Tomlinson Samuel B, Wong Jeremy N, et al. Spatial Distribution of Interictal Spikes Fluctuates over Time and Localizes Seizure Onset Brain. 2020;143:554–569.

90. Kang Xuan, Boly Melanie, Findlay Graham, et al. Quantitative Spatio-Temporal Characterization of Epileptic Spikes Using High Density EEG: Differences between NREM Sleep and REM Sleep Scientific Reports. 2020;10:1673.

91. Asano Eishi, Mihaylova Temenuzhka, Juhász Csaba, Sood Sandeep, Chugani, Harry T. Effect of Sleep on Interictal Spikes and Distribution of Sleep Spindles on Electrocorticography in Children with Focal Epilepsy Clinical neurophysiology: official journal of the International Federation of Clinical Neurophysiology. 2007;118:1360–1368.

92. Ishijima B, Hori T, Yoshimasa N, Fukushima T, Hirakawa K, Sekino H. Neuronal Activities in Human Epileptic Foci and Surrounding Areas Electroencephalography and Clinical Neurophysiology. 1975;39:643–650.

93. Ishijima B. Unitary Analysis of Epileptic Activity in Acute and Chronic Foci and Related Cortex of Cat and Monkey Epilepsia. 1972;13:561–581.

94. Prince D. A, Futamachi K. J. Intracellular Recordings from Chronic Epileptogenic Foci in the Monkey Electroencephalography and Clinical Neurophysiology. 1970;29:496–510.

95. Wyler Allen R., Ojemann George A., Ward Arthur A.. Neurons in Human Epileptic Cortex: Correlation between Unit and EEG Activity Annals of Neurology. 1982;11:301–308.

96. Frazzini Valerio, Whitmarsh Stephen, Lehongre Katia, et al. Human Periventricular Nodular Heterotopia Shows Several Interictal Epileptic Patterns, Associated with Hyperexcitability of Neuronal Firing bioRxiv. 2022:816173.

97. Cohen Ivan, Navarro Vincent, Clemenceau Stéphane, Baulac Michel, Miles Richard. On the Origin of Interictal Activity in Human Temporal Lobe Epilepsy in Vitro Science, New Series. 2002;298:1418–1421.

98. Wittner Lucia, Huberfeld Gilles, Clémenceau Stéphane, et al. The Epileptic Human Hippocampal Cornu Ammonis 2 Region Generates Spontaneous Interictal-like Activity in Vitro Brain. 2009;132:3032–3046.

99. de Curtis M., Radici C., Forti M. Cellular Mechanisms Underlying Spontaneous Interictal Spikes in an Acute Model of Focal Cortical Epileptogenesis Neuroscience. 1999;88:107–117.

100. Traub R D, Miles R, Jefferys J G. Synaptic and Intrinsic Conductances Shape Picrotoxin-Induced Synchronized after-Discharges in the Guinea-Pig Hippocampal Slice. The Journal of Physiology. 1993;461:525–547.

101. Staba Richard J., Wilson Charles L., Bragin Anatol, Fried Itzhak, Engel Jerome. Sleep States Differentiate Single Neuron Activity Recorded from Human Epileptic Hippocampus, Entorhinal Cortex, and Subiculum The Journal of Neuroscience. 2002;22:5694–5704.

102. Calvin William H., Sypert George W., Ward Arthur A. Structured Timing Patterns within Bursts from Epileptic Neurons in Undrugged Monkey Cortex Experimental Neurology. 1968;21:535–549.

103. Wyler A. R., Fetz E. E., Ward A. A. Spontaneous Firing Patterns of Epileptic Neurons in the Monkey Motor Cortex Experimental Neurology. 1973;40:567–585.

104. Connors Barry W.. Initiation of Synchronized Neuronal Bursting in Neocortex Nature. 1984;310:685–687.

105. Prince David A., Wong Robert K.S.. Human Epileptic Neurons Studied in Vitro Brain Research. 1981;210:323–333.

106. Hofer Katharina T., Kandrács Ágnes, Tóth Kinga, et al. Bursting of Excitatory Cells Is Linked to Interictal Epileptic Discharge Generation in Humans Scientific Reports. 2022;12:6280.

107. Gast Heidemarie, Niediek Johannes, Schindler Kaspar, et al. Burst Firing of Single Neurons in the Human Medial Temporal Lobe Changes before Epileptic Seizures Clinical Neurophysiology. 2016;127:3329–3334.

108. Colder Brian W., Frysinger Robert C., Wilson Charles L., Harper Ronald M., Engel Jerome. Decreased Neuronal Burst Discharge Near Site of Seizure Onset in Epileptic Human Temporal Lobes Epilepsia. 1996;37:113–121.

109. Wilson Scott B., Emerson Ronald. Spike Detection: A Review and Comparison of Algorithms Clinical Neurophysiology. 2002;113:1873–1881.

110. Halford Jonathan J.. Computerized Epileptiform Transient Detection in the Scalp Electroencephalogram: Obstacles to Progress and the Example of Computerized ECG Interpretation Clinical Neurophysiology. 2009;120:1909–1915.

111. Halford Jonathan J., Schalkoff Robert J., Zhou Jing, et al. Standardized Database Development for EEG Epileptiform Transient Detection: EEGnet Scoring System and Machine Learning Analysis Journal of Neuroscience Methods. 2013;212:308–316.

112. da Silva Lourenço Catarina, Tjepkema-Cloostermans Marleen C., van Putten Michel J. A. M.. Machine Learning for Detection of Interictal Epileptiform Discharges Clinical Neurophysiology. 2021;132:1433–1443.

113. Brown Merritt W., Porter Brenda E., Dlugos Dennis J., et al. Comparison of Novel Computer Detectors and Human Performance for Spike Detection in Intracranial EEG Clinical Neurophysiology. 2007;118:1744–1752.

114. Nonclercq Antoine, Foulon Martine, Verheulpen Denis, et al. Cluster-Based Spike Detection Algorithm Adapts to Interpatient and Intrapatient Variation in Spike Morphology Journal of Neuroscience Methods. 2012;210:259–265.

115. Scheuer Mark L., Bagic Anto, Wilson Scott B.. Spike Detection: Inter-reader Agreement and a Statistical Turing Test on a Large Data Set Clinical Neurophysiology. 2017;128:243–250.

116. Reus E. E. M., Cox F. M. E., van Dijk J. G., Visser G. H. Automated Spike Detection: Which Software Package? Seizure. 2022;95:33–37.

117. Krishnan Balu, Vlachos Ioannis, Faith Aaron, et al. A Novel Spatiotemporal Analysis of Peri-Ictal Spiking to Probe the Relation of Spikes and Seizures in Epilepsy Annals of biomedical engineering. 2014;42:1606.

118. Janca Radek, Jezdik Petr, Cmejla Roman, et al. Detection of Interictal Epileptiform Discharges Using Signal Envelope Distribution Modelling: Application to Epileptic and Non-Epileptic Intracranial Recordings Brain Topography. 2015;28:172–183.

119. Peter-Derex Laure, Klimes Petr, Latreille Véronique, Bouhadoun Sarah, Dubeau François, Frauscher Birgit. Sleep Disruption in Epilepsy: Ictal and Interictal Epileptic Activity Matter Annals of Neurology. 2020;88:907–920.

120. Lourenço Catarina, Tjepkema-Cloostermans Marleen C., Teixeira Luís F., van Putten Michel J. A. M.. Deep Learning for Interictal Epileptiform Discharge Detection from Scalp EEG Recordings in XV Mediterranean Conference on Medical and Biological Engineering and Computing – MEDICON 2019 (Henriques Jorge, Neves Nuno, de Carvalho Paulo., eds.) IFMBE Proceedings(Cham):1984–1997 Springer International Publishing 2020.

121. Constantino Alexander C., Sisterson Nathaniel D., Zaher Naoir, Urban Alexandra, Richardson R. Mark, Kokkinos Vasileios. Expert-Level Intracranial Electroencephalogram Ictal Pattern Detection by a Deep Learning Neural Network Frontiers in Neurology. 2021;12:603868.

122. Fukumori Kosuke, Thu Nguyen Hoang Thien, Yoshida Noboru, Tanaka Toshihisa. Fully Data-driven Convolutional Filters with Deep Learning Models for Epileptic Spike Detection in ICASSP 2019 – 2019 IEEE International Conference on Acoustics, Speech and Signal Processing (ICASSP):2772–2776 2019.

123. Tjepkema-Cloostermans Marleen C., de Carvalho Rafael C.V., van Putten Michel J.A.M.. Deep Learning for Detection of Focal Epileptiform Discharges from Scalp EEG Recordings Clinical Neurophysiology. 2018;129:2191–2196.

124. Fürbass Franz, Kural Mustafa Aykut, Gritsch Gerhard, Hartmann Manfred, Kluge Tilmann, Beniczky Sándor. An Artificial Intelligence-Based EEG Algorithm for Detection of Epileptiform EEG Discharges: Validation against the Diagnostic Gold Standard Clinical Neurophysiology. 2020;131:1174–1179.

125. Antoniades Andreas, Spyrou Loukianos, Martin-Lopez David, et al. Detection of Interictal Discharges With Convolutional Neural Networks Using Discrete Ordered Multichannel Intracranial EEG IEEE Transactions on Neural Systems and Rehabilitation Engineering. 2017;25:2285–2294.

126. Medvedev A. V., Agoureeva G. I., Murro A. M. A Long Short-Term Memory Neural Network for the Detection of Epileptiform Spikes and High Frequency Oscillations Scientific Reports. 2019;9:19374.

127. Geng David, Alkhachroum Ayham, Bicchi Manuel A. Melo, Jagid Jonathan R., Cajigas Iahn, Chen Zhe Sage. Deep Learning for Robust Detection of Interictal Epileptiform Discharges Journal of Neural Engineering. 2021;18:056015.

128. Tanaka Fabio Henrique Kiyoiti dos Santos, Aranha Claus. Data Augmentation Using GANs arXiv:1904.09135 [cs, stat]. 2019.

129. Bagheri Elham, Jin Jing, Dauwels Justin, Cash Sydney, Westover M. Brandon. A Fast Machine Learning Approach to Facilitate the Detection of Interictal Epileptiform Discharges in the Scalp Electroencephalogram Journal of Neuroscience Methods. 2019;326:108362.

130. Wang Qitong, Whitmarsh Stephen, Navarro Vincent, Palpanas Themis. iEDeaL: A Deep Learning Framework for Detecting Highly Imbalanced Interictal Epileptiform Discharges Proceedings of the VLDB Endowment. 2023;16.

131. Horak Peter C., Meisenhelter Stephen, Testorf Markus E., Connolly Andrew C., Davis Kathryn A., Jobst Barbara C.. Implementation and Evaluation of an Interictal Spike Detector in SPIE Optical Engineering + Applications (Bones Philip J., Fiddy Michael A., Millane Rick P.., eds.)(San Diego, California, United States):96000N 2015.

132. Hao Yongfu, Khoo Hui Ming, von Ellenrieder Nicolas, Zazubovits Natalja, Gotman Jean. DeepIED: An Epileptic Discharge Detector for EEG-fMRI Based on Deep Learning NeuroImage: Clinical. 2018;17:962–975.

133. Lodder Shaun S., Putten Michel J. A. M.. A Self-Adapting System for the Automated Detection of Inter-Ictal Epileptiform Discharges PLOS ONE. 2014;9:e85180.

134. Quon Robert J., Meisenhelter Stephen, Camp Edward J., et al. AiED: Artificial Intelligence for the Detection of Intracranial Interictal Epileptiform Discharges Clinical Neurophysiology. 2022;133:1–8.

135. Lehongre Katia, Lambrecq Virginie, Whitmarsh Stephen, et al. Long-Term Deep Intracerebral Microelectrode Recordings in Patients with Drug-Resistant Epilepsy: Proposed Guidelines Based on 10-Year Experience NeuroImage. 2022:119116.

136. Fried I., Wilson C. L., Maidment N. T., et al. Cerebral Microdialysis Combined with Single-Neuron and Electroencephalographic Recording in Neurosurgical Patients. Technical Note Journal of Neurosurgery. 1999;91:697–705.

137. Mathon Bertrand, Clemenceau Stéphane, Hasboun Dominique, et al. Safety Profile of Intracranial Electrode Implantation for Video-EEG Recordings in Drug-Resistant Focal Epilepsy Journal of Neurology. 2015;262:2699–2712.

138. Pérez-García Fernando, Lehongre K., Bardinet Eric, et al. Automatic Segmentation Of Depth Electrodes Implanted In Epileptic Patients: A Modular Tool Adaptable To Multicentric Protocols Epilepsia. 2015;56:227.

139. Fedorov Andriy, Beichel Reinhard, Kalpathy-Cramer Jayashree, et al. 3D Slicer as an Image Computing Platform for the Quantitative Imaging Network Magnetic Resonance Imaging. 2012;30:1323–1341.

140. Xia Mingrui, Wang Jinhui, He Yong. BrainNet Viewer: A Network Visualization Tool for Human Brain Connectomics PLOS ONE. 2013;8:e68910.

141. Berry Richard B., Brooks Rita, Gamaldo Charlene, et al. AASM Scoring Manual Updates for 2017 (Version 2.4) Journal of clinical sleep medicine: JCSM: official publication of the American Academy of Sleep Medicine. 2017;13:665–666.

142. Dauvilliers Y, Billiard M. Aspects Du Sommeil Normal in Les Troubles de Sommeil:5–17 Paris: Masson 2005.

143. Oostenveld Robert, Fries Pascal, Maris Eric, Schoffelen Jan-Mathijs. FieldTrip: Open Source Software for Advanced Analysis of MEG, EEG, and Invasive Electrophysiological Data Computational Intelligence and Neuroscience. 2011;2011:1–9.

144. Hirsch L J, LaRoche S M, Gaspard N, et al. American Clinical Neurophysiology Society’s Standardized Critical Care EEG Terminology: 2012 Version Journal of Clinical Neurophysiology. 2013;30:27.

145. Hirsch Lawrence J., Fong Michael W. K., Leitinger Markus, et al. American Clinical Neurophysiology Society’s Standardized Critical Care EEG Terminology: 2021 Version Journal of Clinical Neurophysiology. 2021;38:1–29.

146. Yger Pierre, Spampinato Giulia LB, Esposito Elric, et al. A Spike Sorting Toolbox for up to Thousands of Electrodes Validated with Ground Truth Recordings in Vitro and in Vivo eLife. 2018;7.

147. Maris Eric, Oostenveld Robert. Nonparametric Statistical Testing of EEG– and MEG-data Journal of Neuroscience Methods. 2007;164:177–190.

148. Steriade M., Nuñez A., Amzica F.. A Novel Slow (< 1 Hz) Oscillation of Neocortical Neurons in Vivo: Depolarizing and Hyperpolarizing Components The Journal of Neuroscience: The Official Journal of the Society for Neuroscience. 1993;13:3252–3265.

149. Massimini Marcello, Rosanova Mario, Mariotti Maurizio. EEG Slow (∼1 Hz) Waves Are Associated With Nonstationarity of Thalamo-Cortical Sensory Processing in the Sleeping Human Journal of Neurophysiology. 2003;89:1205–1213.

